# Transcriptional regulation of disease-relevant microglial activation programs

**DOI:** 10.1101/2025.10.12.681832

**Authors:** Amanda McQuade, Reet Mishra, Venus Hagan, Weiwei Liang, Peter J. Colias, Vincent Cele Castillo, Bianca Gonzalez, Justin P. Lubin, Verena Haage, Victoria Marshe, Masashi Fujita, Thomas Ta, Layla Gomes, Olivia Teter, Xia Han, Nathaniel Robichaud, Sarah E. Chasins, Jessica E. Rexach, Philip L. De Jager, James K. Nuñez, Martin Kampmann

## Abstract

Microglia, the brain’s innate immune cells, can adopt a wide variety of activation states relevant to health and disease. Dysregulation of microglial activation occurs in numerous brain disorders, and driving or inhibiting specific states could be therapeutic. To discover regulators of microglia activation states, we conducted CRISPR interference screens in iPSC-derived microglia for inhibitors and activators of six microglial states. We characterized 31 regulators at the single-cell transcriptomic and cell-surface proteome level in two distinct iPSC-derived microglia models, uncovering new protein markers of relevant states. We functionally characterized several multi- state regulators. *ZNF532* and *PRDM1* knockdown drive disease-associated, lipid-rich signatures and enhance phagocytosis while showing opposing effects on antigen-presentation signatures. *DNMT1* knockdown results in widespread loss of methylation, activating negative regulators of interferon signaling. These findings provide a framework to direct microglial activation to selectively enrich microglial activation states, define their functional outputs, and inform future therapies.

**Highlights:**

- CRISPRi screening reveals novel regulators of six microglia activation states
- Multi-modal single-cell screens highlight new markers of microglial states
- Different iPSC-microglia models show different landscapes of activation states at baseline
- Loss of DNMT1 leads to widespread DNA demethylation, promoting some states but limiting the interferon-response state
- Loss of PRDM1 or ZNF532 drives the microglial disease-associated state

## Introduction

Microglia are highly plastic brain-resident macrophages that continuously monitor the parenchyma to support cognitive function and brain health^1,2^. In response to insult or injury, microglia rapidly trigger transcriptional activation programs, or states, that allow them to restore homeostasis. Advances in single-cell technologies have markedly expanded our understanding of the diversity of microglial activation states^3^. Along with the discovery of new activation states, these data enabled identification of specific states that are enriched or depleted in disease conditions^4–7^. In rare cases, these activation states have been linked directly to expression of disease risk-associated genes, highlighting the importance of these states for brain health^4,5,8,9^.

The disease-associated microglial (DAM) activation state has been shown to increase during aging and is further expanded in neurodegenerative diseases including Alzheimer’s disease, Parkinson’s disease, Frontotemporal dementia, and Amyotrophic Lateral Sclerosis^5,6,10–12^. This activation state has been most deeply studied in the context of Alzheimer’s disease where DAM cells cluster around beta-amyloid plaques and contribute to their compaction and clearance^13–17^. The Alzheimer’s disease risk gene Triggering Receptor Expressed on Myeloid cells 2 (*TREM2*) is a key regulator of this activation state. Limiting DAM activation is hypothesized to be the mechanism whereby TREM2 mutations affect disease risk^16,18–21^.

Lipid-rich microglia have also been associated with Alzheimer’s disease as well as multiple sclerosis, neuropathic pain, and stroke^8,22–26^. Microglia containing lipid droplets are thought to represent a highly phagocytic state, clearing myelin and dead cell debris. Whether the lipid droplets themselves represent a beneficial sequestration of excess lipids or induce dysfunction is still debated and may differ depending on the disease, though there is evidence that reducing lipid droplets may be beneficial^25,27,28^.

Interferon-responsive microglia are activated in response to viral infections such as HIV-associated neurocognitive disorders^29^, but are also present in sterile conditions, particularly during developmental windows of synaptic refinement^30^. Interferon-responsive microglia have been associated with developmental conditions including autism spectrum disorder^29–31^, but are also present in aging, multiple sclerosis, and tauopathies, where these cells induce aberrant synaptic pruning and contribute to neuroinflammation^32–36^.

Antigen-presenting microglia interact closely with T cells, a disease mechanism increasingly recognized in Alzheimer’s disease and related dementias^37–40^. Chemokine-high microglia are less well understood but have been profiled in Alzheimer’s disease, FTD/ALS, and autism spectrum disorders^41–43^. Increased chemokine secretion may impact infiltration of peripheral immune cells into the brain^38,44^ and has also been shown to increase microglial surveillance^18,45–47^. Lastly, the activation of many of these states is concordant with decreases in homeostatic microglia markers, though it is not known whether this decrease is necessary to permit activation or a separate but often convergent pathway. Homeostatic microglia express high levels of purinergic receptors that are used to surveille the brain environment.

Despite the critical importance of microglial activation states to their physiological function, we do not understand the transcriptional regulation of these activation states. While regulators of individual states have been predicted, we currently only have validated regulators of the DAM state. Even for these regulators, we lack insights into their global effect on microglial activation programs as multi-state regulators have not been comprehensively identified. The ability to selectively drive or inhibit microglial states will enable their functional characterization, support our understanding of how specific states contribute to disease, and enable the reprogramming of these states for therapeutic purposes.

Gene regulatory network (GRN) prediction algorithms such as SCENIC^48^ have been applied to nominate regulators of specific states with some success^49–55^. However, GRN prediction algorithms are still being refined to optimize accuracy, and all predictions need to be functionally validated. Furthermore, these algorithms often rely on predicted transcription factor binding motifs. Many of these motifs have never been functionally validated. Additionally, for ∼40% of transcription factors, DNA consensus sequences have not been identified^56^, limiting the power for GRN prediction algorithms to nominate these regulators. Thus, there is a critical need to directly perturb and discover transcription factor drivers of microglial activation.

We directly perturbed transcription factors (TF) and transcriptional regulators to determine regulators of six microglial activation states: homeostatic, disease-associated (DAM), lipid-rich, antigen-presenting, interferon-responsive, and chemokine. We performed these screens in iPSC-derived microglia because this model recapitulates a heterogeneous population of microglial activation states at baseline allowing for discovery of both drivers and inhibitors of each state^57^. The results of our screens showed some overlap with predicted regulators^51,54,55^, and additionally uncovered many novel regulators of microglial activation states, including TFs that target complex combinations of state signatures.

We selected 31 dynamic regulators from the six large-scale CRISPRi screens to investigate more deeply at the single-cell level. While most descriptions of microglial states are based purely on transcriptomic profiling, we aimed to obtain a deeper characterization by also monitoring the cell surface proteome in the same individual cells. For this purpose, we combined CRISPR-droplet RNA-sequencing (CROP-seq) with Cellular Indexing of Transcriptomes and Epitopes sequencing (CITE-seq). To increase the robustness of our findings, we performed these single-cell based studies using two microglia differentiation protocols^57,58^, and observed that each model exhibited distinct activation profiles at baseline. This observation highlights the importance of selecting appropriate models that recapitulate the disease state or activation trajectories of interest.

Lastly, we validated our findings for selected regulators, Zinc Finger Protein 532 (*ZNF532*), PR-domain zinc finger protein 1 (*PRDM1)*, Signal Transducer and Activator of Transcription 2 knockdown (*STAT2*), and DNA methyltransferase I (*DNMT1),* by orthogonal CRISPRi and RNAi approaches. *ZNF532* and *PRDM1* each negatively regulate the DAM signature but have opposing effects on the antigen-presentation and homeostatic signatures. *STAT2* and *DNMT1* KD microglia show marked decreases in interferon-responsive microglia and lysosomal function, though in *DNMT1* KD this response is paired with increases in homeostatic and lipid-rich signatures. We highlight a potential mechanism of interferon-response inhibition through ISG15 by enzymatic methylation sequencing.

Together, our data represent the first large-scale perturbation study to determine regulators of microglial activation states, including multi-state regulators. We highlight relationships between activation trajectories, discuss differences across two microglial models, and provide multi-modal validation of the impact of these perturbations on microglial gene expression, protein expression, and function. We anticipate that this multi-modal dataset will provide a valuable resource to isolate and enrich microglial activation states, explore their downstream effects, and guide future mechanistic and translational research.

## Results

### CRISPRi screens uncover regulators of microglial activation states

To determine transcriptional regulators of microglial activation states, we performed six fluorescence-associated cell sorting (FACS)-based CRISPR interference (CRISPRi) screens using a single-guide RNA (sgRNA) library targeting ∼1600 transcription factors and transcriptional regulators (Figure 1A). We selected markers for six activation states that have been reported^3,40,41,54,57,59–61^ to impact brain homeostasis and disease risk: Homeostatic (marked by P2RY12), disease-associated (DAM, marked by CD9), antigen-presenting (marked by HLA-DMB), lipid-rich (marked by BODIPY, neutral lipid dye), chemokine (marked by CCL13), and interferon-responsive (marked by IFIT1).

**Figure 1.**
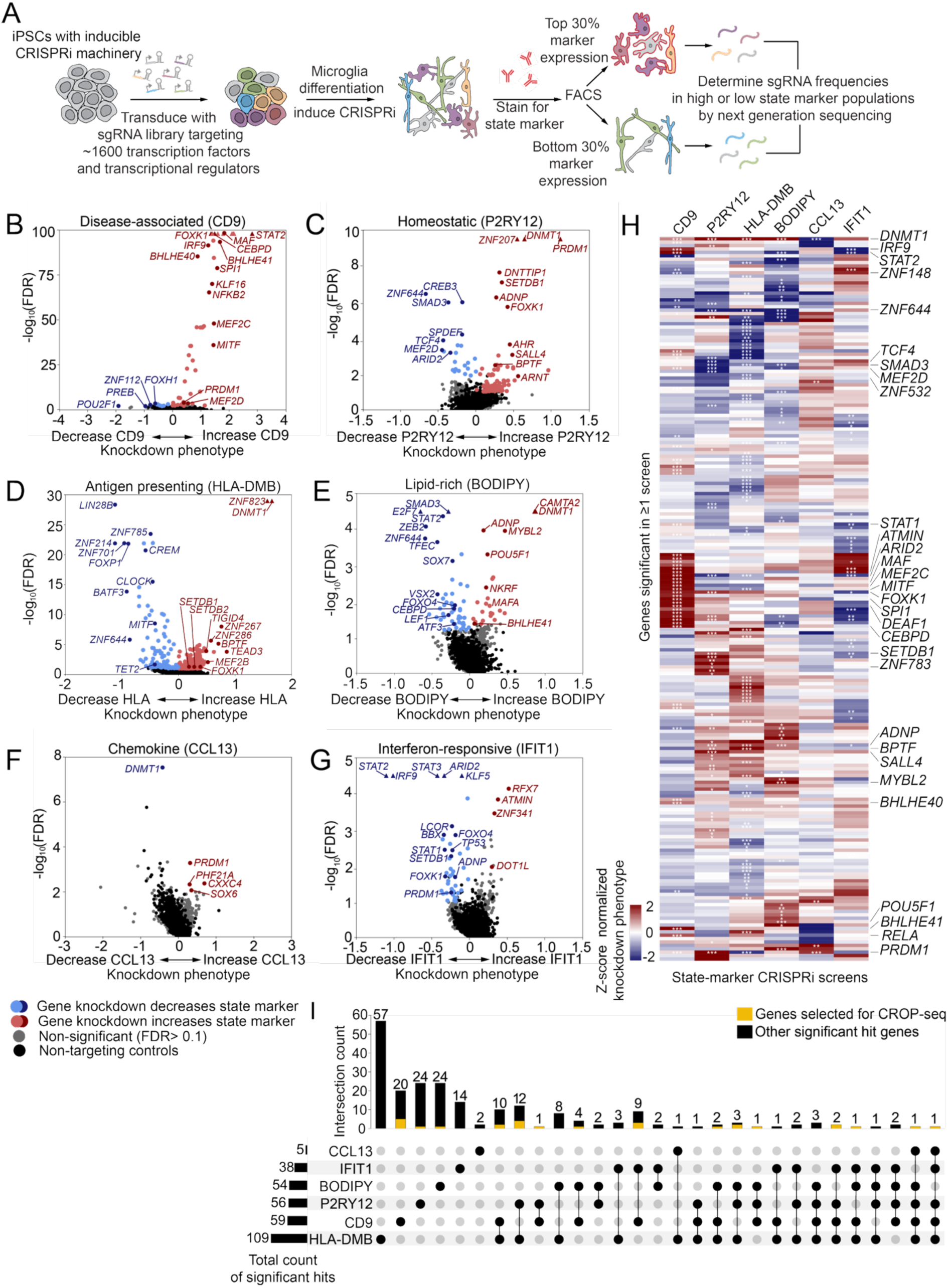
CRISPRi screens uncover transcriptional regulators of microglial activation states. (A) Schematic of FACS-based CRISPRi screens. iPSCs expressing inducible CRISPRi machinery were transduced with a pooled library of sgRNAs targeting ∼1600 transcription factors and transcriptional regulators. iPSCs were differentiated into microglia (iTF-MG protocol) and CRISPRi knockdown was induced with trimethoprim. Microglia were stained with markers for six activation states. The 30% of cells with the lowest and highest marker expression were separated by FACS. Genomic DNA was isolated from each population and the cassette encoding sgRNAs was amplified and sequenced using next-generation sequencing. Comparison of the frequencies of cells expressing each sgRNA to those expressing non-targeting control sgRNAs was used to determine hits. (B-G) Volcano plots of screening results for each state. Knockdown phenotype is calculated as a log_2_ ratio of counts in the high and low marker expression populations normalized to the standard deviation of non-targeting control cells. CRISPRi targets that significantly (FDR < 0.01) increase or decrease state marker expression are colored in red or blue, respectively. Gene knockdowns that do not significantly alter state marker expression (FDR ≥ 0.01) are shown in grey. Quasi-genes generated from random sets of non-targeting control sgRNAs to model a null distribution are shown in black (see also Table S1). (H) Heatmap representing screen results for all genes (rows) that were significant in one or more of the screens shown in H. Each column represents an individual screen. Color scale represents the knockdown phenotype and significance represents FDR. Labelled genes were selected for secondary screens with deep single-cell phenotyping. *FDR < 0.05, **FDR < 0.01, ***FDR < 0.001 FDR. (I) Upset plot representing the number of significant hits (FDR < 0.05) for each screen that are unique to a specific screen or shared across multiple screens. Shading in gold represents genes that were selected for secondary screens with deep single-cell phenotyping, same as labels in (H).

We previously developed a transcription factor (TF)-directed microglia differentiation protocol (iTF-MG) that reduces the bottlenecking of sgRNA libraries that occurs in more complex differentiation protocols^57^. iTF-MG were differentiated and stained for each of the state markers described above independently (P2RY12, CD9, HLA-DMB, BODIPY, CCL13, IFIT1) and sorted using FACS for high (top 30%) or low (bottom 30%) expression of each marker (Figure 1A). Enrichment of cells expressing each sgRNA in the high and low populations was quantified and compared against cells expressing non-targeting control sgRNAs.

The screen based on expression of CD9 (DAM) uncovered a strong skew of positive regulators with 50 genes for which knockdown increased CD9 expression and 8 genes for which knockdown decreased CD9 expression (Figure 1B, Table S1). This suggests that many transcriptional regulators actively suppress the disease-associated (DAM) signature. Indeed, we found that many of our positive hits had been previously identified to be enriched in human postnatal microglia over fetal microglia (*PRDM1, SPI1, IRF1, KLF2, KLF6, KLF8, KLF16*)^62^. The DAM state is not highly expressed in healthy adult microglia, so this overlap supports our hypothesis that the DAM state may be actively suppressed. Comparing our functional data to previous predicted regulators of the DAM state^14,53,55^, we report that perturbation of the following predicted regulators does indeed alter expression of CD9: *BLHLE40, BHLHE41, SPI1, MAF, MEF2C, MITF, STAT1, CEBPD, KLF6*.

Screening based on P2RY12 expression (homeostatic state) identified 80 genes for which knockdown increased P2RY12 and 20 genes for which knockdown reduced P2RY12 (Figure 1C, Table S1). This bias towards discovery of genes for which knockdown increases P2RY12 is not surprising, given that the iTF-MG microglia model used for these screens has relatively low levels of P2RY12 at baseline, although an updated media composition we developed for these experiments increased P2RY12 expression (Figure S1A-B). These results highlight additional transcription factors that could be activated to further increase homeostatic markers including SMAD3, which functions downstream of TGFB1, a key homeostatic signal for microglia^63^ (discussed below).

Screening based on HLA-DMB expression (antigen-presenting state) produced the highest number of hits, identifying 197 genes for which knockdown increased HLA-DMB and 88 genes for which knockdown reduced HLA-DMB (Figure 1D, Table S1). Of interest, this screen revealed a potential connection between regulators of DNA methylation and expression of HLA-DMB. One DNA methyltransferase (*DNMT1*) and several readers of DNA methylation (*SETDB1, SETDB2, MECP2*) were all positive hits in the screen. Additionally the transcription factor *YY1*, that stabilizes DNA methylation^64^, was also a positive hit. Further corroborating this relationship, a DNA demethylase (*TET2)* was a negative hit. Together, this data suggests that hypo-methylation of DNA in microglia may increase antigen presentation. Studies of various cancer tissues have also highlighted this relationship using chemical inhibitors of DNMT1^65,66^.

Screening based on BODIPY staining (lipid-rich state) revealed 46 negative hits for which knockdown reduced lipid load and 23 positive hits for which knockdown increased lipid load (Figure 1E, Table S1). Interestingly, we found 14 overlapping hits between the CD9 and BODIPY screens with 11 of these impacting these states in opposing directions (Figure 1H,I, Table S1). This may suggest a relationship between the two activation states, possibly reflecting increased lipid droplet clearance in disease-associated microglia. Expression of *CEBPD*, for example, is known to promote lipid accumulation in macrophages^67^, which our knockdown screen confirmed in microglia, and was also determined to be a negative regulator of the disease-associated state in our CD9 screen (Figure 1B).

The screen based on CCL13 expression (chemokine state) revealed only 4 hits (Figure 1F), suggesting a limited ability to alter this state by transcription factor perturbation, which is compatible with previous reports that chemokine expression is tightly regulated by post-translational signaling^68,69^. Interestingly, three out of the four genes for which knockdown increased CCL13 (CXXC4, PHF21A, SOX6) have been linked to neurodevelopmental disorders, though their impact is primarily hypothesized to occur in non-microglial cell types^70–72^.

Screening based on expression of IFIT1 (interferon-responsive state) was performed with a pre-stimulation of IFNβ to induce microglia to enter the interferon-responsive state. This screen revealed an enrichment of negative regulators with 44 hits for which knockdown decreased interferon-response and 4 hits for which knockdown increased interferon response (Figure 1G, Table S1). The iTF-MG microglia model is known to have high levels of interferon signaling at baseline^57,73^; however, our recent genome-wide screen for this state produced a more even distribution of positive and negative regulators^73^. As expected, we found that the interferon-stimulated gene factor 3 (ISGF3) complex consisting of *STAT1, STAT2,* and *IRF9* are all strong negative hits in this screen as these TFs are key amplifiers of type I interferon signaling. We also confirmed our previous discovery^74^ that knockdown of the ASD-risk gene Activity-Dependent Neuroprotector Homeobox (*ADNP*) biases microglia away from the interferon-responsive state. Furthermore, we uncovered Tumor Protein 53 (*TP53*) as a regulator of interferon-responsive microglia. The relationship between interferon signaling and TP53 has been explored in the context of cancer^75,76^ and in models of Down syndrome and Alzheimer’s disease, where blocking interferon signaling has been shown to limit senescence^77^. Our data suggests a reciprocal relationship where *TP53* expression also regulates interferon responses.

Considering the hits across all six screens collectively, we find that 65% of the significant regulators impacted an individual activation state marker, while 34% impacted two or more state markers (Figure 1C). Intriguingly, two factors impacted 5 out of 6 states: DNA methyltransferase I (*DNMT1*) and PR domain zinc finger protein 1 (*PRDM1*) (Figure 1H,I).

### Single-cell RNA-sequencing highlights distinct state landscapes across iPSC-microglia models

While the six marker-based screens are informative to nominate regulators of microglial activation states, an individual protein marker is often not sufficient to represent the full state. It is possible that some of the regulators in our screens increase only the state marker protein and not the full signature of proteins associated with a given state. To address this caveat, we conducted secondary screens in which we combined CRISPR perturbations with single-cell RNA sequencing (using CRISPR-droplet RNA-sequencing (CROP-seq) approach^78^ to reveal the full transcriptomic changes downstream of 31 regulators selected from the primary screen hits (labelled in Figure 1H,I). We focused in on complex regulators that impact multiple microglia activation states, as our study was specifically designed to reveal such nuanced mechanisms. For this targeted library, we included two sgRNAs for each gene target and five non-targeting sgRNAs (knockdown validation in Figure S2). As there are clear differences between mRNA and protein expression, particularly in microglia^79^, and surface receptor expression is an important indicator of activation, we also profiled the cell surfaceome of cells in this screen by using antibodies coupled to oligonucleotide barcodes that were read out in parallel with single cell transcriptomes. We used the Cellular Indexing of Transcriptomes and Epitopes sequencing (CITE-seq) approach with a curated antibody library for the study of immune cells and microglia^80^ (Figure 2A, Table S2).

**Figure 2.**
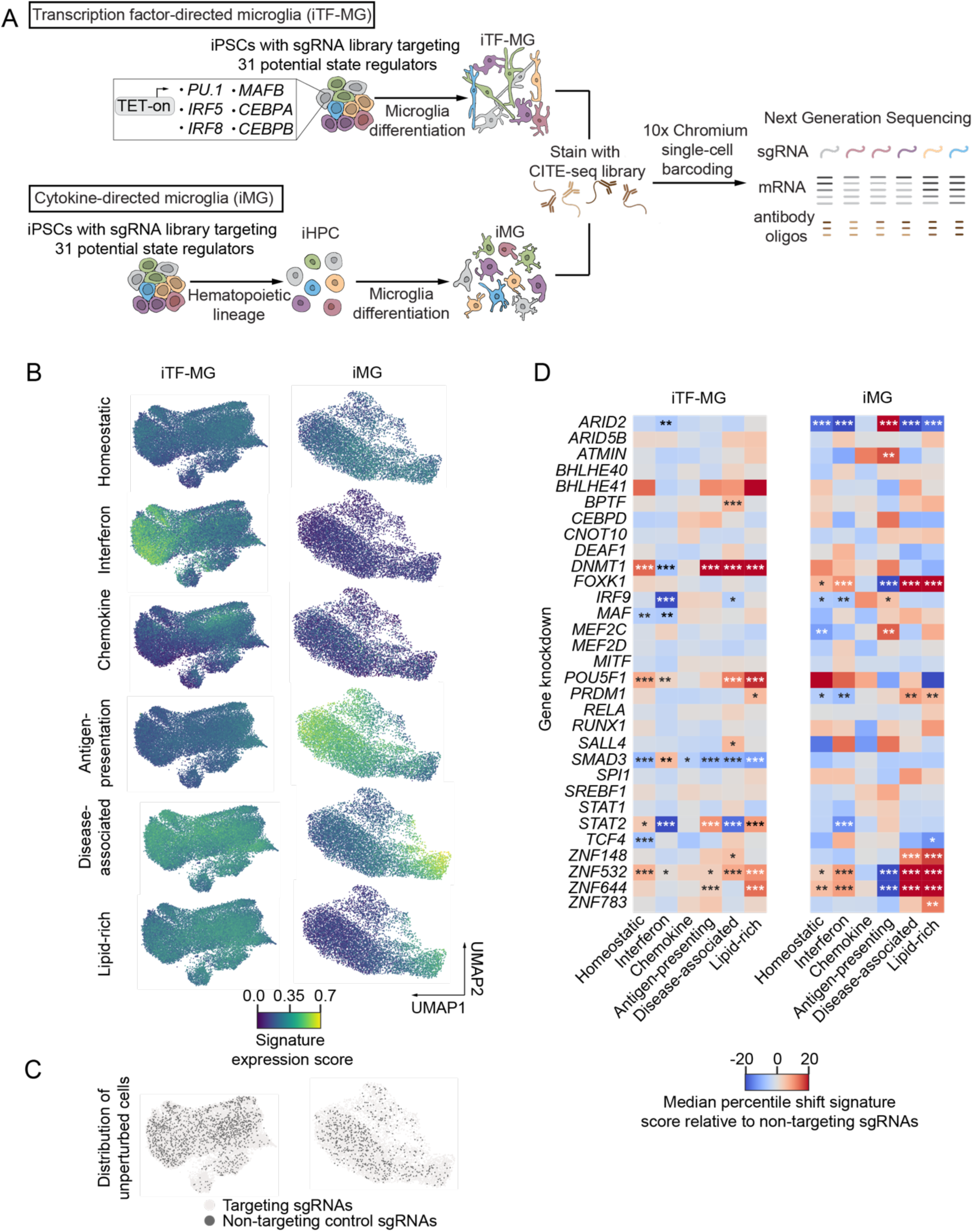
Single-cell RNA-sequencing highlights distinct state landscapes across iPSC-microglia models. (A) Schematic of the paired CROP-seq and CITE-seq experiments. iPSCs expressing inducible CRISPRi machinery were transduced with a pooled library of sgRNAs targeting 31 transcription factors and transcriptional regulators. 2 sgRNAs per gene, 5 non-targeting control sgRNAs. iPSCs were differentiated into microglia using the iTF-MG protocol from Dräger et al. 2022^57^ (top) or iMG protocol from McQuade et al. 2018^58^ (bottom). After differentiation, microglia were stained with a pooled oligo-tagged antibody library from Haage et al. 2025^83^ before single-cell barcoding with 10x Chromium and next generation sequencing of the sgRNA protospacer, mRNA, and antibody oligonucleotides (see also Table S2). (B) UMAP representation of iTF-MG (left) and iMG (right) mRNA space. Cells are colored by state signature expression scores calculated by UCell (see methods, see Figure S3). (C) UMAP representation of iTF-MG (left) and iMG (right) mRNA space, colored to show cells with non-targeting control sgRNAs (dark grey, see also Figure S3). (D) Heatmaps of median percentile shift in signature expression scores relative to non-targeting control cells calculated for each gene knockdown (rows) for each state (columns). *FDR < 0.05, **FDR < 0.01, ***FDR < 0.001 Mann Whitney U-test.

Our large-scale transcription factor screens (Figure 1) revealed asymmetric discovery rates for positive and negative regulators that appear to be dependent on the baseline activation state of the iTF-MG model (low homeostatic, high interferon). For this reason, we performed our CROP-seq and CITE-seq analysis in two distinct iPSC-derived microglia protocols: transcription factor-directed differentiation (iTF-MG)^57^ and developmental cytokine-directed differentiation (iMG)^58^. While both models express microglia identity genes (Figure S3A-B), each differentiation protocol resulted in a unique composition of baseline activation states (Figure 2B,C, Figure S3C,D).

Traditional clustering algorithms assume that individual cells must occupy one distinct state, however this is not aligned with our current understanding of microglial activation. Modeling microglial states as independent factors that can combine modularly is a better representation of the underlying biology^80,81^. Thus, we utilized an activation state signature curated from the literature to define gene expression programs for each state that can be measured independently across cells (Figure S3C, Table S2). Expression enrichment scores for these signature gene sets were calculated for every cell using UCell. Using these signature scores, we found that iTF-MG robustly express the interferon-responsive and chemokine signature genes (Figure 2B) corroborating our previous single-cell studies using this model^57,74^. Additionally, we find that disease-associated (DAM) and lipid-high signatures are high across most iTF-MG cells while antigen-presenting genes are uniformly low (Figure 2B). In the cytokine-directed iMG model, we find cells with high levels of genes in the disease-associated (DAM) and lipid-high signatures we well as antigen-presenting signatures. In the absence of stimulation, the iMG model shows exceptionally low levels of interferon-responsive and chemokine state signatures (Figure 2B), however these states can be activated with stimulation (Figure S1C).

As a complementary analysis and to report the relevance of our model systems in human brain-derived microglia, we aligned our CROP-seq dataset to a 26-factor framework developed de-novo from microglia from 161 diverse human brain samples^80^ (Figure S4). These factors reflect activation programs that are non-orthogonal and can be overlapping. Mirroring our analysis of curated state signatures, iTF-MG showed high expression of factors related to interferon-response (factor 20) and disease-associated factors (factor 8, 11, 26). iMG showed high expression of antigen-presenting (15) and oxidative phosphorylation factors (factor 8, 18, 21).

To further understand differences between iTF-MG and iMG, we analyzed the secretome of microglia from each protocol by nELISA^82^ for 978 proteins (Figure S2E, Table S3). We found iTF-MG secreted higher levels of chemoattractants including RARRES2, VEGF-R1, CXCL6, CXCL12α and CXCL12β often seen in perivascular macrophages. iMG secreted higher levels of proteins associated with the DAM state including TREM2, GPNMB, OPN, C3, Galectin-1, and Galectin-3 (Figure S2E).

### Perturbation of microglial activation states at the transcriptional level reveals complex state regulators

Since our initial screens were based on a single marker readout, we expected that a subset of the hits from these screens would not affect the full transcriptomic state signature. Indeed, knockdown of several hits did not significantly alter any of the states we define in this study (Figure 2C, comparison of FACS-based and CROP-seq screen results in Figure S5). These genes may be responsible for partial transcriptomic signatures (Table S2) or perhaps act on the state marker proteins through mechanisms unrelated to microglial activation. In some cases, this may also be due to low cell counts for specific perturbations, which decreases statistical power (Table S2).

Interestingly, many transcription factor perturbations produced distinct activation profiles in the two microglia models. For example, while *STAT2* knockdown decreased the interferon-response state in both iTF-MG and iMG, *STAT2* knockdown also had broader effects on multiple states in iTF-MG, but not in iMG (Figure 2D). Similarly, while *SMAD3* knockdown decreased homeostatic state genes in iTF-MG, this trend was not significant in iMG (Figure 2D). We also report transcription factor perturbations that are consistent across iTF-MG and iMG including regulators of the lipid-rich state (*PRDM1, ZNF148, ZNF532, ZNF644*).

Our RNA-sequencing data highlighted better separation for chemokine and interferon-responsive states in the iTF-MG model and better separation for DAM and lipid-rich states in the iMG model. We further tested the ability of each microglia protocol to drive the DAM state by perturbation of a canonical DAM regulator, Triggering Receptor Expressed on Myeloid Cells 2 (TREM2). While both microglia protocols show reduced DAM markers and increased homeostatic markers after knockdown of *TREM2*, results in iMG were more robust (Figure S1D-K). Based on these findings, follow up studies into specific regulators were primarily performed using the microglia model that best recapitulates each state (iTF-MG for interferon-responsive and chemokine, iMG for DAM, lipid-rich, and antigen-presentation).

In both microglia models, we found high concordance of the disease-associated (DAM) and lipid-rich mRNA signatures (Figure 2D). For iTF-MG, three of the six gene knockdowns that induced DAM also increased the lipid-rich signature. For iMG, five out of the six gene knockdowns that induced DAM also increased the lipid-rich signature. In our original transcription factor-wide regulator screens, we found a marked opposition between the results of the anti-CD9 (DAM) and BODIPY (lipid) screens (Figure 1B, E). Together, this suggests that the lipid-rich gene signature may not always be correlated with high levels of lipid droplets. This data highlights the importance of confirming functional outcomes that are expected to be correlated with gene signatures as the lipid-rich microglia state is often assumed to represent cells with high lipid droplet load.

### Microglial activation states are correlated with distinct surface protein signatures

The microglia “sensome” is a collection of surface receptors microglia use to detect and react to environmental changes. Cell-surface protein expression is dynamically regulated as microglia activate and alter the functional properties of the cell. Overall, we found good concordance between expression of individual proteins and their cognate mRNA, however some proteins showed no correlation with mRNA expression of the same gene or even a small negative correlation (Figure S6). Negative correlations for the same surface proteins were also seen in Haage et al. 2025^83^ which reports the design and deployment of the CITE-seq panel also used here. The fact that transcriptomes are not fully predictive of the cell surfaceome highlights the importance of our multi-omic profiling strategy.

A subset of our strongest transcription factor perturbations that impacted microglial activation also differentially altered specific surface protein abundances (iTF-MG: Figure 3A, D-F; iMG: Figure 3G, J-L). These surface proteins further support our understanding of the perturbed gene’s impact on microglial function. For example, the integrins increased in *DNMT1* and *STAT2* KD iTF-MG have been shown to promote beta-amyloid clearance and limit neuroinflammation^84,85^.

**Figure 3.**
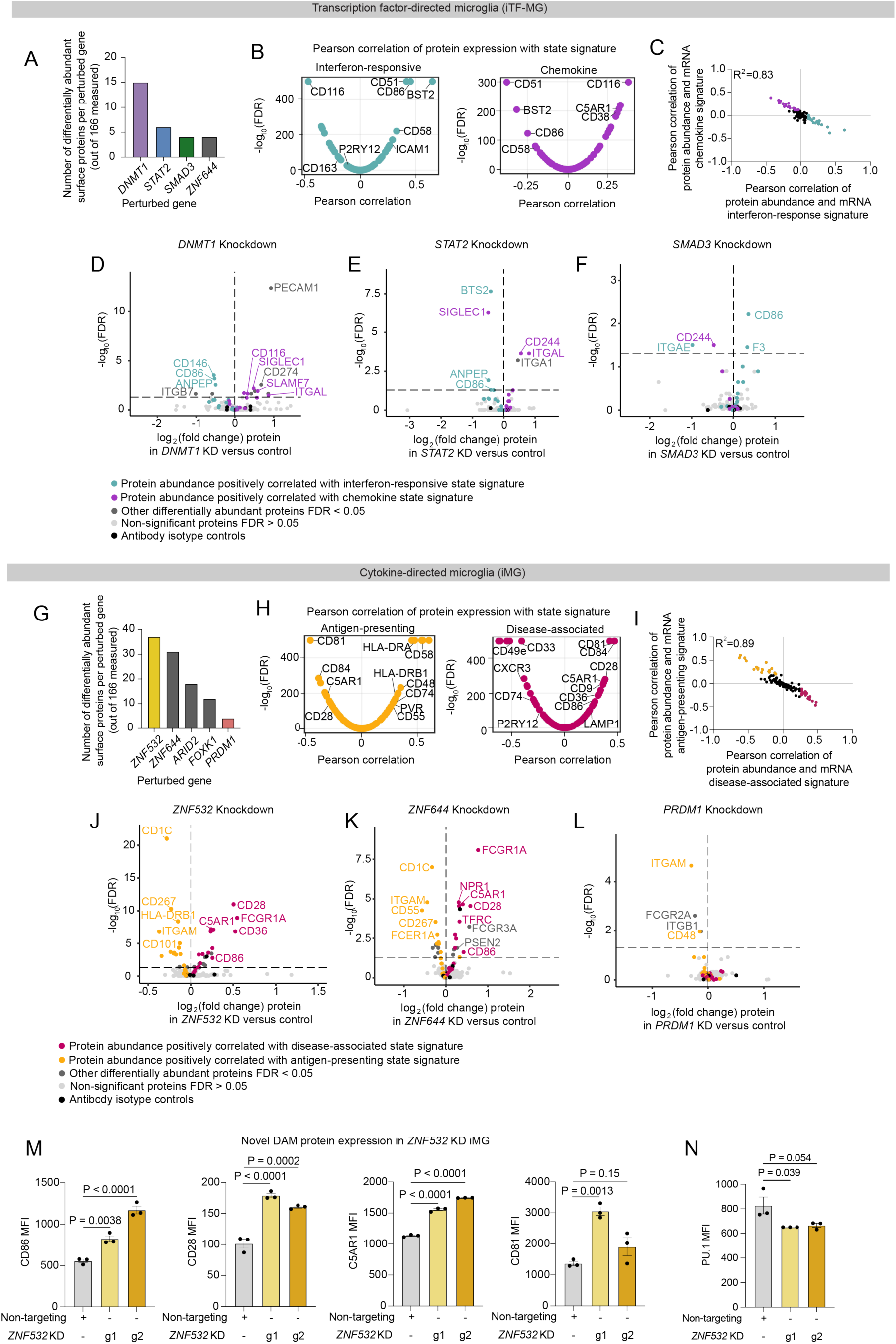
Paired CROP-seq and CITE-seq identifies new surface proteins marking microglial activation states. (A-F) Analysis in iTF-MG. (G-N) Analysis in iMG. (A) Bar graph displaying all perturbations with significant differentially abundant proteins compared to protein abundance in the non-targeting control cells. (B) Pearson correlation of protein expression from CITE-seq analysis and mRNA state signature gene expression from CROP-seq analysis. Correlation with interferon-responsive state is shown on left. Correlation with chemokine state is shown on right. (C) Negative correlation of protein abundance associated with interferon-responsive state (X-axis, top 20 associated proteins in teal) and chemokine state (Y-axis, top 20 associated proteins in purple). (D-F) Volcano plot of differentially abundant proteins in (D) *DNMT1* KD, (E) *STAT2* KD, or (F) *SMAD3* KD versus non-targeting control. FDR ≤ 0.05. Top 20 DAPs that are associated with interferon-responsive and chemokine state are colored teal and purple respectively. CITE-seq antibodies that are antibody isotype controls are colored black. (G) Bar graph displaying all perturbations with significant differentially abundant proteins compared to protein abundance in the non-targeting control cells. (H) Pearson correlation of protein expression from CITE-seq analysis and mRNA state signature gene expression from CROP-seq analysis. Correlation with antigen-presenting state is shown on left. Correlation with disease-associated state is shown on right. (I) Negative correlation of protein abundance associated with disease-associated state (X-axis, top 20 associated proteins in pink) and antigen-presenting state (Y-axis, top 20 associated proteins in yellow). (J-L) Volcano plot of differentially abundant proteins in (J) *ZNF532* KD, (K) *ZNF644* KD, or (L) *PRDM1* KD versus non-targeting control. FDR ≤ 0.05. Top 20 DAPs that are associated with disease-associated and antigen-presenting state are colored pink and yellow respectively. CITE-seq antibodies that are antibody isotype controls are colored black. (M) Median fluorescence intensity (MFI) of cell surface proteins positively correlated with the disease-associated state by flow cytometry in *ZNF532* KD microglia (two separate CRISPRi gRNAs (g1 and g2); yellow) versus non-targeting controls (grey). Each point represents an independent well, n ≥ 10,000 cells analyzed per well. P values from one way ANOVA. (N) Median fluorescence intensity (MFI) of PU.1 in *ZNF532* KD microglia (yellow) versus non-targeting controls (grey). Each point represents an independent well, n ≥ 10,000 cells analyzed per well. P values from one way ANOVA.

Beyond analysis of specific perturbations, we merged our CROP-seq and CITE-seq analyses to correlate abundance of 166 surface proteins with each microglia state (Figure 3B,C,H,I, Table S2). Intriguingly, these results corroborated the mRNA analysis, where iTF-MG showed distinct separation of interferon-responsive and chemokine states. At the protein level, proteins positively correlated with the interferon-response state (CD51, BST2, CD86, CD58, ICAM1, etc.) were negatively correlated with the chemokine state (Figure 3B,C). We found a similar relationship between the antigen-presenting and disease-associated states in iMG, with CD81, C5AR1, CD9, CD28 being positively correlated with the DAM state and negatively correlated with the antigen-presenting state (Figure 3H,I). The antigen-presenting state is also positively correlated with known antigen-presenting proteins including HLA-DRB1, HLA-DRA, CD58, CD48, CD74 etc. (Figure 3H, Table S2). These relationships were mirrored in primary human microglia isolated from surgical and post-mortem specimens^80^ which supports the robustness of our *in vitro* microglia models in discovering relationships between surface proteins and activation that reflect *ex vivo* relationships.

Next, we validated surface expression of our novel DAM markers (CD81, C5AR1, CD86, and CD28) in *ZNF532* KD iMG (Figure 3M). Knockdown of Zinc Finger Protein 532 (ZNF532) was identified in our CROP-seq as a putative driver of the DAM state (Figure 2D, explored below). *ZNF532* KD iMG also showed increased levels of our novel DAM markers from CITE-seq (Figure 3J) which we confirmed with two independent CRISPRi sgRNAs (Figure 3M). Of special interest, CD28 was recently confirmed as a DAM marker *in vivo* and is associated with a protective PU.1^low^ microglia subtype^86^. *ZNF532* KD iMG similarly showed a modest decrease in PU.1 (Figure 3N). Together, these results highlight the power of combining single-cell transcriptomics, CITE-seq, and pooled CRISPR perturbations to deeply profile, nominate, and validate functional regulators of disease-relevant states.

### Loss of ZNF532 promotes the disease-associated microglia signature and limits the antigen-presentation signature

Knockdown (KD) of *ZNF532* robustly promoted disease-associated (DAM), lipid-rich, and homeostatic activation signatures across both iTF-MG and iMG (Figure 2D). Since the iMG model shows higher fidelity to the DAM state (Figure S1, S3), we chose to study ZNF532 in iMG. Deeper investigation into the transcriptomic signature of *ZNF532* KD iMG revealed an increase in genes associated with the DAM and lipid-rich states and a strong decrease in antigen-presenting genes (Figure 4A-D, Table S2).

**Figure 4.**
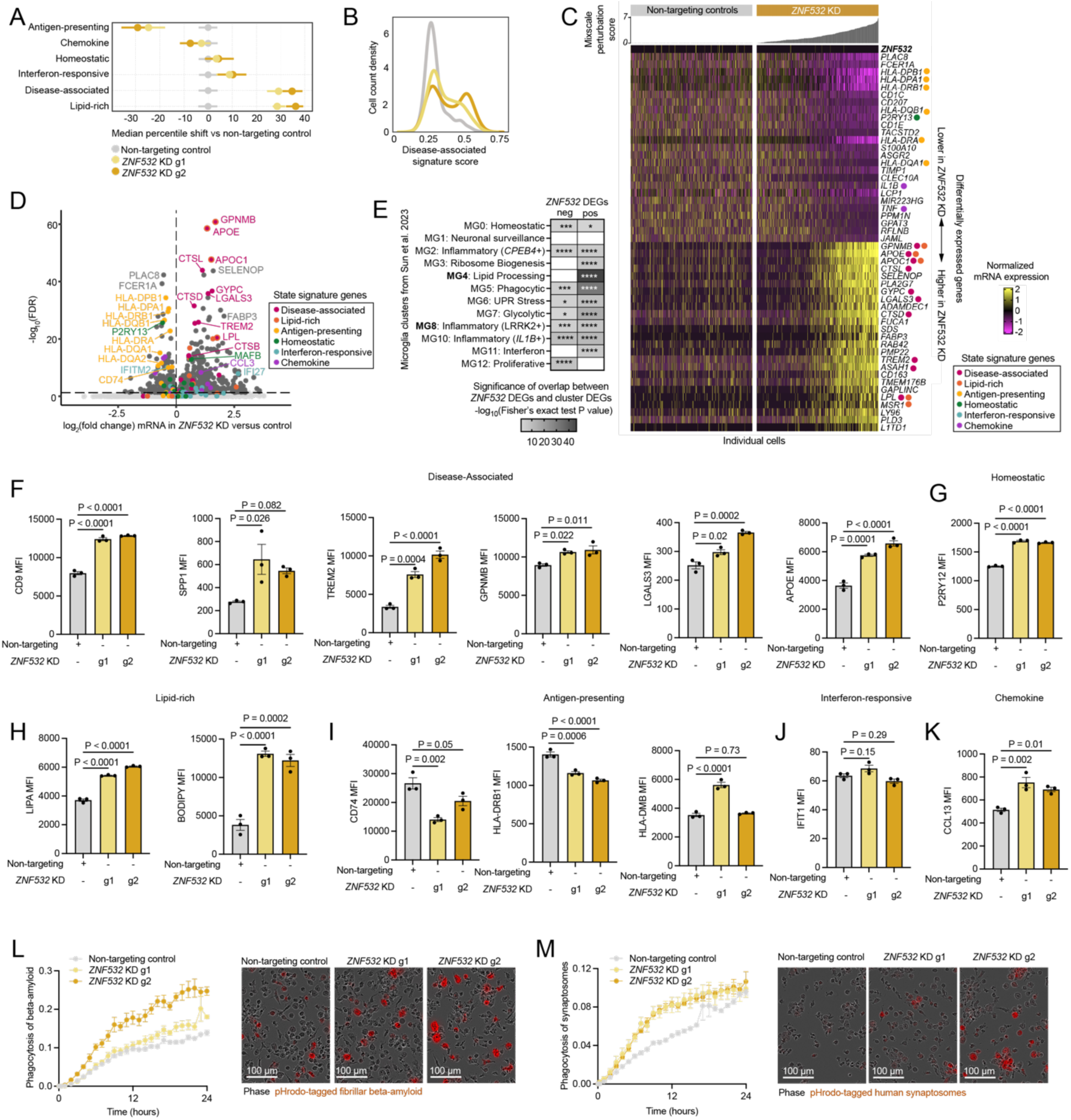
Loss of ZNF532 promotes the disease-associated microglia signature and limits the antigen-presenting signature in iMG. (A) Median percentile shift of state signature scores in *ZNF5321* KD (yellow) versus non-targeting controls (grey). Separate sgRNAs are represented as g1 and g2. Error bars represent 95% confidence interval. (B) Density plot of disease-associated signature score percentile shift for *ZNF532* KD (yellow) versus non-targeting controls (grey). (C) Heatmap of top differentially expressed genes (DEGs) ranked by FDR (25 positive DEGs, 25 negative DEGs). Columns represent individual cells ordered by Mixscale perturbation score (see methods for details). Scores are graphed above the corresponding column (non-targeting control perturbation scores are all zero by definition). Color scale represents the log-normalized mRNA expression. DEGs that appear in our state signature scores are denoted with a colored dot (pink: disease-associated, orange: lipid-rich, yellow: antigen-presenting, green: homeostatic, teal: interferon-responsive, purple: chemokine). (D) Volcano plot of differentially expressed genes in *ZNF532* KD versus non-targeting control. FDR ≤ 0.05. DEGs that appear in our state signature scores colored as above. (E) Heatmap of gene signature overlap analysis of DEGs from *ZNF532* KD with human microglia clusters from Sun et al. 2023^54^. Bolded states are increased in AD patients. Color denotes significance of overlap by Fisher’s exact test P value. Non-significant overlap is white. *P < 0.05, **P < 0.01, ***P < 0.001 ****P < 0.0001. (F-K) Median fluorescence intensity (MFI) of state marker proteins by flow cytometry. *ZNF532* KD microglia (yellow) were compared to non-targeting controls (grey). Each point represents an independent well, n ≥ 10,000 cells analyzed per well. P values from one way ANOVA. (also see Figure S8) (L) Phagocytosis of pHrodo-tagged beta-amyloid in *ZNF5321* KD (yellow) and versus non-targeting controls (grey). n=4 independent wells, 4 images per well. Points represent mean signal per well +/- standard error. Representative images of phase and fluorescent human synaptosomes (right). (M) Phagocytosis of pHrodo-tagged human synaptosomes in *ZNF532* KD (yellow) versus non-targeting controls (grey). n=4 independent wells, 4 images per well. Representative images of phase and fluorescent human synaptosomes (right).

One feature of targeting regulators with CRISPR interference rather than CRISPR knockout is that individual cells in the knockdown population may produce different levels of knockdown. The presence of cells with different extends of knockdown enables us to investigate the dose-response relationship between the knockdown of a transcription factors and the resulting effects on specific signatures. Specifically, we performed weighted differential expression using Mixscale^87^ to identify minimally, moderately, and highly perturbed cells (Figure S7). This analysis revealed that the distribution of DAM gene expression is bimodal in *ZNF532* KD iMG (Figure 4B,C, Figure S7A). In *ZNF532* KD cells with low Mixscale score (corresponding to weaker perturbation), we see little difference in the state signatures compared to non-targeted cells. However, even moderate Mixscale scores for this perturbation produce a strong phenotype with a near maximal response (Figure S7A).

To assess how our perturbations align with transcriptional signatures from human brain-derived microglia, we analyzed the overlap of the DEGs resulting from *ZNF532* KD with genes defining clusters identified from AD patient and control tissue by Sun et al. 2023^54^ (Figure 4E, Table S2). This comparison revealed a strong overlap of DEGs increased in *ZNF532* KD with MG4, a lipid-processing state. In their manuscript, Sun et al. highlight MG3 and MG4 as having significant overlap with the original DAM signature discovered in mouse by Keren-Shaul et al. 2017^4^. By comparing genes increased after *ZNF532* KD directly to the Keren-Shaul DAM signature, we also report significant overlap (P value = 3.06×10^-29^, Fisher’s exact test, Table S2). Our up-regulated genes additionally showed significant overlap with human AD-patient gene signatures as defined by Gerrits et al. 2021^60^, specifically with their clusters 7, 8, 9, and 10, which are increased in AD patients and three of which correlated with beta-amyloid deposition in post-mortem tissue (P value for Cluster 7: 8.52×10^−30^, Cluster 8: 3.75×10^−75^ Cluster 9: 2.22×10^−51^, Cluster 10: 3.34×10^−96^ Fisher’s exact test, Table S2). Genes decreased in *ZNF532* KD vs control overlap with several microglia clusters from Sun et al. including proliferative and inflammatory states. This gene set also significantly overlaps with the negative DEGs from Keren-Shaul et al. DAM (P value = 0.006, Fisher’s exact test, Table S2). Together, these observations show a strong overlap of *ZNF532* KD DEGs with gene signatures relevant to Alzheimer’s disease progression and pathology.

Projecting *ZNF532* KD iMG onto the 26-factor framework from Marshe et al. 2025^80^ revealed a strong increase in the GPNMB^high^, OxPhos-1, and C1q^high^-phagocytic factors and a decrease in HLA^high^/APC, OxPhos-3, and PLCG2^high^ (Figure S4E).

To corroborate our screening data, we generated *ZNF532* CRISPRi knockdown cell lines using two independent sgRNAs (g1 and g2) (Figure S2D). Activation state markers were quantified at the protein level by flow cytometry (Figure 4F-K, Figure S8). This experiment confirmed the increase in DAM (CD9, SPP1, TREM2, GPNMB, LGALS3, APOE), homeostatic (P2RY12), and lipid-rich (LIPA, BODIPY) markers as well as the decreased antigen-presenting markers (CD74, HLA-DRB1; HLA-DMB is increased in g1). CCL13 was also increased at the protein level. IFIT1 showed no change in *ZNF532* KD microglia.

With constitutively active CRISPRi, *ZNF532* expression is reduced throughout the differentiation process. To address whether a shorter-term knockdown in mature microglia would show the same phenotypes, we used siRNA to knock down *ZNF532*. We found that the DAM, homeostatic, chemokine, and antigen-presenting phenotypes were consistent when using siRNA to knock down *ZNF532* in two independent iPSC lines (Figure 9A-N). We also showed that *ZNF532* KD has a similar phenotype using the iTF-MG differentiation protocol, though the effect sizes were minimal (Figure S9O-Q).

### *ZNF532* knockdown promotes phagocytosis of synaptosomes and beta-amyloid

Given the overlap of *ZNF532* KD DEGs with gene signatures that are associated with phagocytosis and correlated with beta-amyloid pathology in patients, we assessed the phagocytic capacity of *ZNF532* KD microglia after exposure to beta-amyloid or human brain-derived synaptosomes. We found that *ZNF532* KD microglia take up more beta-amyloid and synaptosomes than control microglia (Figure 4L, M).

Since the fluorescent signal we quantified could be due to increased uptake of fluorescent substrates or decreased clearance, we assessed lysosomal cathepsin activity by orthogonal measures. Uptake and cleavage of Dye-Quenched Bovine Serum Albumin (DQ-BSA) was also increased in *ZNF532* KD iMG, suggesting that lysosomal cathepsin activity is not impaired (Figure S8C). Analysis of lysosomal cathepsin activity directly also showed no changes in cathepsin B, D, or L (Fig. S7D). We also report no difference in lysosomal abundance as captured by lysotracker, lysosensor, and LAMP1 expression (Figure S8E,F). Together, these results suggest that the increased phagocytosis in *ZNF532* KD microglia is likely due to increased uptake of material and not due to impaired lysosomal degradation.

### Loss of PRDM1 promotes the disease-associated microglia signature

PR-domain zinc finger protein 1 (PRDM1) is another regulator that impacts multiple activation states. The most highly impacted gene signatures in *PRDM1* KD versus control iMG are the disease-associated (DAM) and lipid-processing activation states (Figure 5A-D). Mixscale-weighted differential expression shows a strong enrichment of disease-associated microglia genes up-regulated in *PRDM1* KD microglia (Figure 5C, D, Table S2). Interestingly, Mixscale analysis revealed that, unlike *ZNF532*, *PRDM1* knockdown is not a simple on/off switch, and that expression of DEGs can be tuned based on the level of knockdown achieved with some genes only perturbed in the cells with the highest *PRDM1* KD score (Figure 5C, Figure S7B, Table S2). We found that downregulation of negative DEGs already occurred at lower Mixscale scores (corresponding to weaker perturbation) than upregulated DEGs, which were observed only in cells with higher Mixscale scores (Figure 5C). This may suggest that these downregulated DEGs are direct targets of PRDM1, as it is an inhibitory TF.

**Figure 5.**
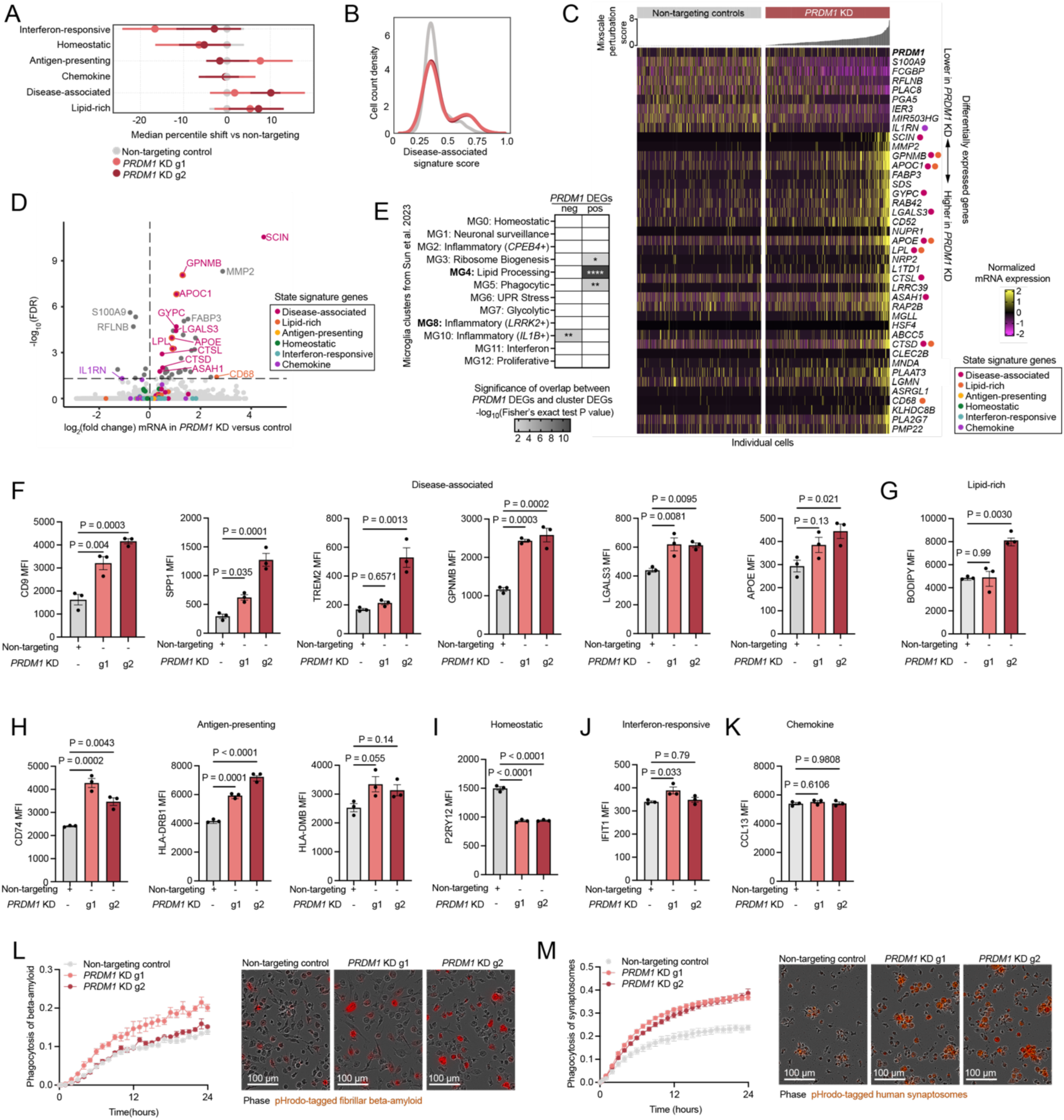
Loss of PRDM1 promotes the disease-associated microglia signature in iMG. (A) Median percentile shift of state signature scores in *PRDM1* KD (salmon, red) versus non-targeting controls (grey). Separate sgRNAs are represented as g1 and g2. Error bars represent 95% confidence interval. (B) Density plot of disease-associated signature score for *PRDM1* KD (salmon, red) versus non-targeting controls (grey). (C) Heatmap of top differentially expressed genes (DEGs) ranked by FDR (32 positive DEGs, 8 (all) negative DEGs). Columns represent individual cells ordered by Mixscale perturbation score (see Methods for details). Scores are graphed above the corresponding column (non-targeting control perturbation scores are all zero by definition). Color scale represents the log-normalized mRNA expression. DEGs that appear in our state signature scores are denoted with a colored dot (pink: disease-associated, orange: lipid-rich, yellow: antigen-presenting, green: homeostatic, teal: interferon-responsive, purple: chemokine). (D) Volcano plot of differentially expressed genes in *PRDM1* KD versus non-targeting control. FDR ≤ 0.05. DEGs that appear in our state signature scores colored as above. (E) Heatmap of gene signature overlap analysis of DEGs from *PRDM1* KD with human microglia clusters from Sun et al. 2023^54^. Bolded states are increased in AD patients. Color denotes significance of overlap by Fisher’s exact test P value. Non-significant overlap is white. *P < 0.05, **P < 0.01, ***P < 0.001 ****P < 0.0001. (F-K) Median fluorescence intensity (MFI) of state marker proteins by flow cytometry. *PRDM1* KD microglia (salmon, red) were compared to non-targeting controls (grey). Each point represents an independent well, n ≥ 10,000 cells analyzed per well. P values from one way ANOVA. (see also Figure S10) (L) Phagocytosis of pHrodo-tagged beta-amyloid in *PRDM1* KD (salmon, red) and versus non-targeting controls (grey). n=4 independent wells, 4 images per well. Points represent mean signal per well +/- standard error. Representative images of phase and fluorescent human synaptosomes (right). (M) Phagocytosis of pHrodo-tagged human synaptosomes in *PRDM1* KD (salmon, red) versus non-targeting controls (grey). n=4 independent wells, 4 images per well. Representative images of phase and fluorescent human synaptosomes (right).

Similarly to *ZNF532* KD, genes up-regulated in *PRDM1* KD overlap with MG3, MG4, and MG5 from Sun et al. 2023^54^ which represent their ribosome biogenesis, phagocytic, and lipid-processing states (Figure 5E, Table S2). We additionally find significant overlap with the Keren-Shaul et al. DAM signature (P value = 1.1×10^−4^, Fisher’s exact test, Table S5) and Gerrits et al. clusters associated with amyloid deposition. (P value for Cluster 7: 3.9×10^−6^, Cluster 8: 1.1×10^−10^ Cluster 9: 1.5×10^−6^, Cluster 10: 1.4×10^−10^ Fisher’s exact test, Table S2). We report very few genes with lower expression in *PRDM1* KD microglia (Figure 5D, Table S2). However, these genes are significantly overlapping with MG10 from Sun et al. 2023, an early inflammatory state high for IL-1β (Figure 5E, Table S2). Alignment of *PRDM1* KD iMG to the 26-factor framework from Marshe et al. 2025, showed higher levels of GPNMB^high^ and APOE^high^ factors and decreased levels of PLCG2^high^ and CX3CR1^high^ (Figure S4F).

We generated two independent *PRDM1* KD lines and found that one sgRNA (g2) reduced *PRDM1* expression to a greater extent than the other (g1) (Figure S2B). Using these lines, we found a strong increase in proteins associated with the DAM and lipid-processing states (Figure 5F-G). However, unlike *ZNF532* KD, *PRDM1* KD increased antigen-presenting states as well (Figure 5H). In many cases, sgRNA g2 increased expression of these markers to a greater extent than sgRNA g1, further supporting the notion that the downstream consequences *PRDM1* KD may be tunable. Additionally, we found no change in expression of IFIT1 and CCL13 at the protein level, and a modest decrease in P2RY12 (Figure 5I-K). Additional protein markers of these states showed consistent results (Figure S10). These findings remained consistent even when knocking down *PRDM1* in mature iMG using siRNA, in an additional iPSC background, and in iTF-MG (Figure S11).

### *PRDM1* knockdown promotes phagocytosis of synaptosomes and beta-amyloid

As with *ZNF532* KD, we found that *PRDM1* KD microglia have increased uptake of beta-amyloid and synaptosomes relative to control microglia (Figure 5L, M). In this case, we found no difference in cleavage and unquenching of DQ-BSA, no difference in lysotracker signal, a modest decrease in lysosensor signal, and no difference in the function of Cathepsin D or L (Figure S10D,E). Cathepsin B activity and LAMP1 expression was increased only in the more stringent *PRDM1* KD with g2 (Figure S10E,F). We conclude that the increased internalized pHrodo signal in *PRDM1* KD microglia is not a result of impaired lysosomal degradation. The functional conservation between *ZNF532* and *PRDM1* KD iMG suggests that this phenotype is not directly impacted by the antigen-presenting state, since ZNF532 and PRDM1 drive antigen-presenting genes in opposing directions.

### Loss of STAT2 inhibits the interferon-response state

Our marker-based primary screens found that Signal Transducer and Activator of Transcription 2 knockdown (*STAT2* KD) regulates lipid-rich, disease-associated, antigen-presenting and interferon-responsive states in the iTF-MG model. However, the full transcriptomic data highlights interferon-responsive genes as the main category of differentially expressed genes (DEGs) following *STAT2* knockdown (KD) (Figure 6A-D, Table S2). Interestingly, antigen-presenting and lipid-rich signatures were increased at the mRNA level in *STAT2* KD iTF-MG (Figure 6A) whereas staining for cell surface HLA-DMB protein levels or for lipid droplets using BODIPY confirmed the result of the original screens based on same antibody/dye (Figure 6J, K). This finding highlights the importance of multimodal analysis as opposed to identifying microglial activation profiles solely based on mRNA. Similarly, *STAT2* KD significantly decreased the disease-associated microglia mRNA signature, but only a subset of marker proteins (Figure 6G, Figure S13A).

**Figure 6.**
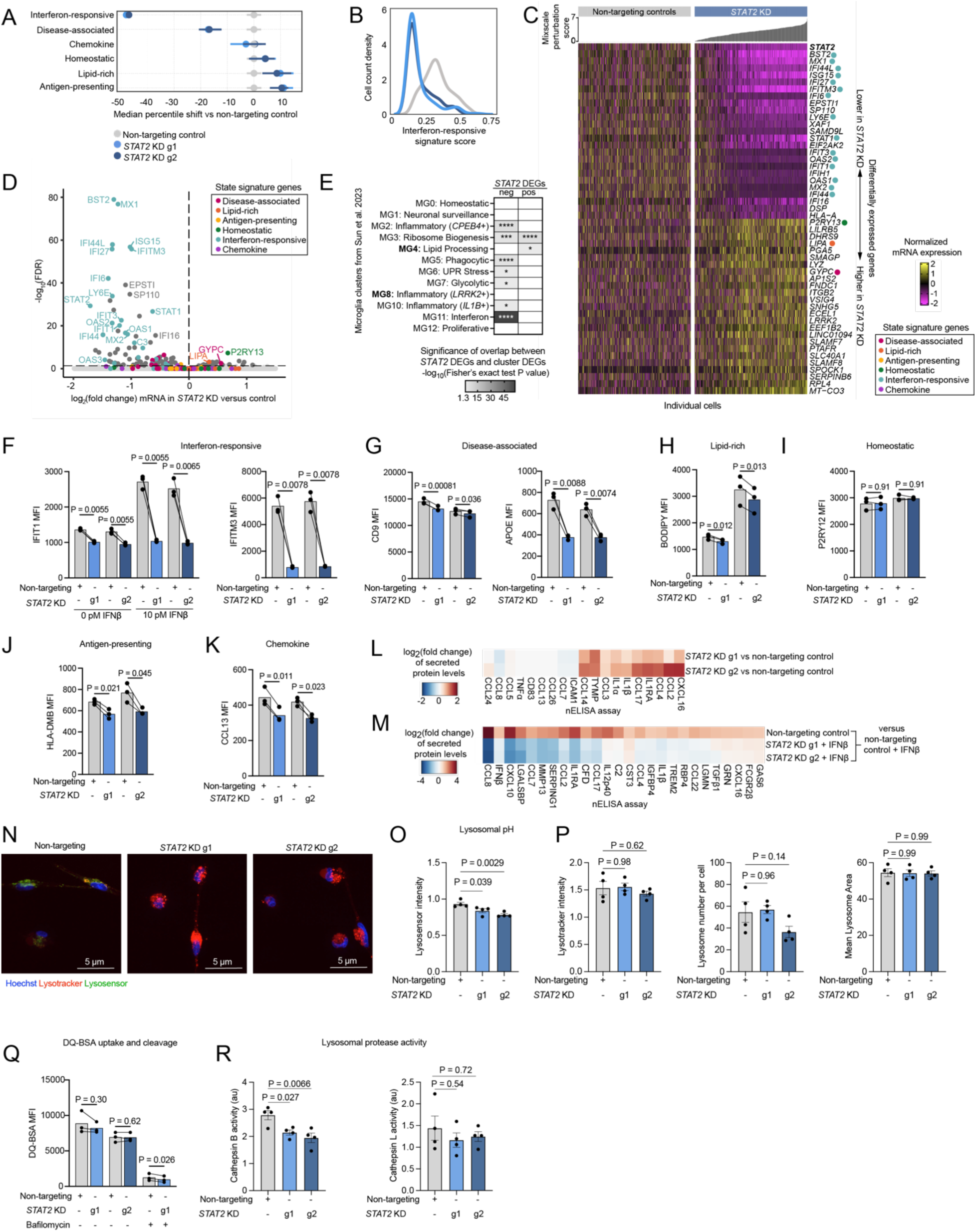
Loss of STAT2 inhibits the interferon-response state in iTF-MG. (A) Median percentile shift of state signature scores in *STAT2* KD (blue) versus non-targeting controls (grey). Separate sgRNAs are represented as g1 and g2. Error bars represent 95% confidence interval. (B) Density plot of interferon-responsive signature score for *STAT2* KD (blue) versus non-targeting controls (grey). (C) Heatmap of top differentially expressed genes (DEGs) ranked by FDR (25 positive DEGs, 25 negative DEGs). Columns represent individual cells ordered by Mixscale perturbation score (see methods for details). Scores are graphed above the corresponding column (non-targeting control perturbation scores are all zero by definition). Color scale represents the log-normalized mRNA expression. DEGs that appear in our state signature scores are denoted with a colored dot (pink: disease-associated, orange: lipid-rich, yellow: antigen-presenting, green: homeostatic, teal: interferon-responsive, purple: chemokine). (D) Volcano plot of differentially expressed genes in *STAT2* KD versus non-targeting control. FDR ≤ 0.05. DEGs that appear in our state signature scores colored as above. (E) Heatmap of gene signature overlap analysis of DEGs from *STAT2* KD with human microglia clusters from Sun et al. 2023^54^. Bolded states are increased in AD patients. Color denotes significance of overlap by Fisher’s exact test P value. Non-significant overlap is white. *P < 0.05, **P < 0.01, ***P < 0.001****P < 0.0001. (F-K) Median fluorescence intensity (MFI) of state marker proteins by flow cytometry. *STAT2* KD microglia (blue) were compared to in-well non-targeting controls (grey) distinguished by nuclear fluorescent proteins (see also Figure S12). Each point represents an independent well, n ≥ 10,000 cells analyzed per well. P values from paired Student’s T-test. (see also Figure S13) (L) Heatmap displaying differential abundance of secreted proteins that define the chemokine state signature. Differential abundance is shown between STAT2 *KD* microglia and non-targeting controls. Protein abundance was measured by nELISA. (M) Heatmap displaying differential abundance of secreted proteins measured by nELISA. Protein list displayed represents all differentially abundant proteins in non-targeting control cells after treatment with 10 pM IFNβ for 24 hrs. (N) Representative images of lysotracker (red), lysosensor (green), and Hoechst (blue) fluorescence in non-targeting control or *STAT2* KD iTF-MG. Scale bar = 5 µm. (O) Quantification of lysosomal pH using lysosensor intensity within lysosomes normalized to Hoechst intensity. N= 4 images per well in 4 separate wells. P value from One-way ANOVA. (P) Quantification of lysotracker intensity within lysosomes normalized to Hoechst intensity (left), lysosomal count (middle) and mean lysosomal area per lysosome (right). N= 4 images per well in 4 separate wells. P value from One-way ANOVA. (Q) Median fluorescence intensity (MFI) DQ-BSA in *STAT2* KD microglia and in-well non-targeting controls. DQ-BSA was added for two hours before analysis to allow for uptake. Bafilomycin A was added four hours before analysis to control wells. (R) Lysosomal protease activity in *STAT2* KD microglia versus non-targeting control microglia measured by fluorescence of MagicRed signal for Cathepsin B (left) or Cathepsin L (right). n=4 wells, 15,000 cells per well. P value from One-way ANOVA.

For genes decreased in *STAT2* KD microglia, we found strong overlap with clusters MG11, MG2, and MG5 from Sun et al. (Figure 6E, Table S2). Cluster MG11 represents an interferon-responsive microglia cluster. MG2 represents a *CPEB4*-high inflammatory state the authors predicted to be regulated by IRF8. MG5 represents high expression of phagocytosis-related genes. For genes increased in *STAT2* KD microglia, we found significant overlap with MG3 and MG4. MG3 represents microglia high in ribosome biogenesis. MG4 represents lipid processing high microglia and is increased in AD patients. Using the 26-factor model, *STAT2* KD iTF-MG show decreased levels of IFN-I response and C1q^high^/phagocytic state and increased levels of OxPhos-2 and OxPhos-3 (Figure S4B).

Protein markers of activation states were quantified by flow cytometry with two independent *STAT2* CRISPRi sgRNAs in iTF-MG (Figure 6G-K, Figure S2A). This confirmed that *STAT2* knockdown limits activation of interferon-responsive marker IFIT1 both at baseline and after stimulation with IFNβ (Figure 6F). We also find reduced levels of a separate marker of the interferon-responsive state, IFITM3 (Figure 6F). The cellular response to IFNβ includes release of cytokines to amplify immune responses. We challenged *STAT2* KD and control iTF-MG with IFNβ and found STAT2 KD iTF-MG limit this downstream effect of IFNβ response as well (Figure 6M). Beyond interferon, we found lower levels of APOE, BODIPY, HLA-DMB, and CCL13 staining in *STAT2* knockdown microglia (Figure 6G, H, J, K). P2RY12 and CD9 expression were not perturbed in *STAT2* KD microglia (Figure 6G, I). Additional protein markers of these states showed consistent results (Figure S13A-C). These findings are consistent when knocking down STAT2 in mature iTF-MG using siRNA, in a separate iPSC line, and in the iMG model (Figure S13D-M).

Although *STAT2* KD iTF-MG showed no change to chemokine signature genes at the RNA level and decreased levels of CCL13 in the cell, we analyzed cytokine levels in *STAT2* KD conditioned media from iTF-MG. We found increased abundance of ∼50% of the chemokines and cytokines that define our chemokine activation state (Figure 6L). This suggests that *STAT2* KD does indeed increase chemokine secretion despite the lack of evidence at the transcriptomic level, providing further support for the need for multi-modal analyses at the transcriptomic, protein, and functional (secretion) levels.

STAT2 modifies transcription downstream of IFNβ by forming the Interferon-Stimulated Gene Factor 3 Complex (ISGF3) with STAT1 and IRF9. While STAT1 and IRF9 had more modest knockdown at the transcriptional level (Figure S2O), we were still able to detect similar phenotypes in *IRF9* KD iTF-MG (Figure S14).

Additionally, we report that *SMAD3* KD in iTF-MG globally produced the opposite effect of *STAT2* or *IRF9* KD. *SMAD3* KD iTF-MG showed increased response to interferon, higher levels of chemokines within the cells, decreased P2RY12, and increased expression of markers for the lipid-rich and DAM states (Figure S14). SMAD3 is the primary effector of TGF-β signaling, a critical homeostatic signal for microglia, therefore, we are not surprised to find that microglia lacking SMAD3 lose homeostatic signals and increase markers of activation.

### *STAT2* KD decreases lysosomal acidity

Interferon-responsive microglia are responsible for pruning neuronal synapses in development and hypothesized to do the same in disease. In epithelial cells, the interferon response increases lysosomal acidification and protease activity^88^, which are critical for phagocytosis. We assessed lysosomal properties in *STAT2* KD microglia to determine if tonic low-level interferon-response contributes to lysosomal degradative capacity in microglia. We found that *STAT2* KD microglia have reduced levels of the pH-dependent lysosomal indicator, lysosensor, but found no difference in fluorescence of DQ-BSA, a BSA conjugate that increases fluorescence after cleavage of quenching dyes by lysosomal cathepsins (Figure 6N-Q). However, direct analysis of Cathepsin B showed lower activity in *STAT2* KD iTF-MG (Figure 6R). There were no significant differences in the number or size of lysosomes (Figure 6P).

### Loss of DNMT1 is generally permissive to activation excluding the interferon-responsive state

DNA-methyltransferase I (DNMT1) is an epigenetic regulator that we found to strongly perturb microglial activation signatures. While *DNMT1* knockout is lethal during embryonic development^89,90^, using inducible CRISPRi to reduce *DNMT1* expression modestly during microglial differentiation avoids any toxicity (Figure S15A). In iTF-MG, knockdown of *DNMT1* enriched expression of all states except chemokine and interferon-responsive (Figure 7A). The interferon-responsive state signature was significantly decreased in *DNMT1* KD microglia (Figure 7A-D, Table S2). Analysis of *DNMT1* KD by Mixscale score shows that even weak knockdown of *DNMT1* achieved dramatic changes in state signatures, which only minimally increased further with stronger knockdown (Figure S7D).

**Figure 7.**
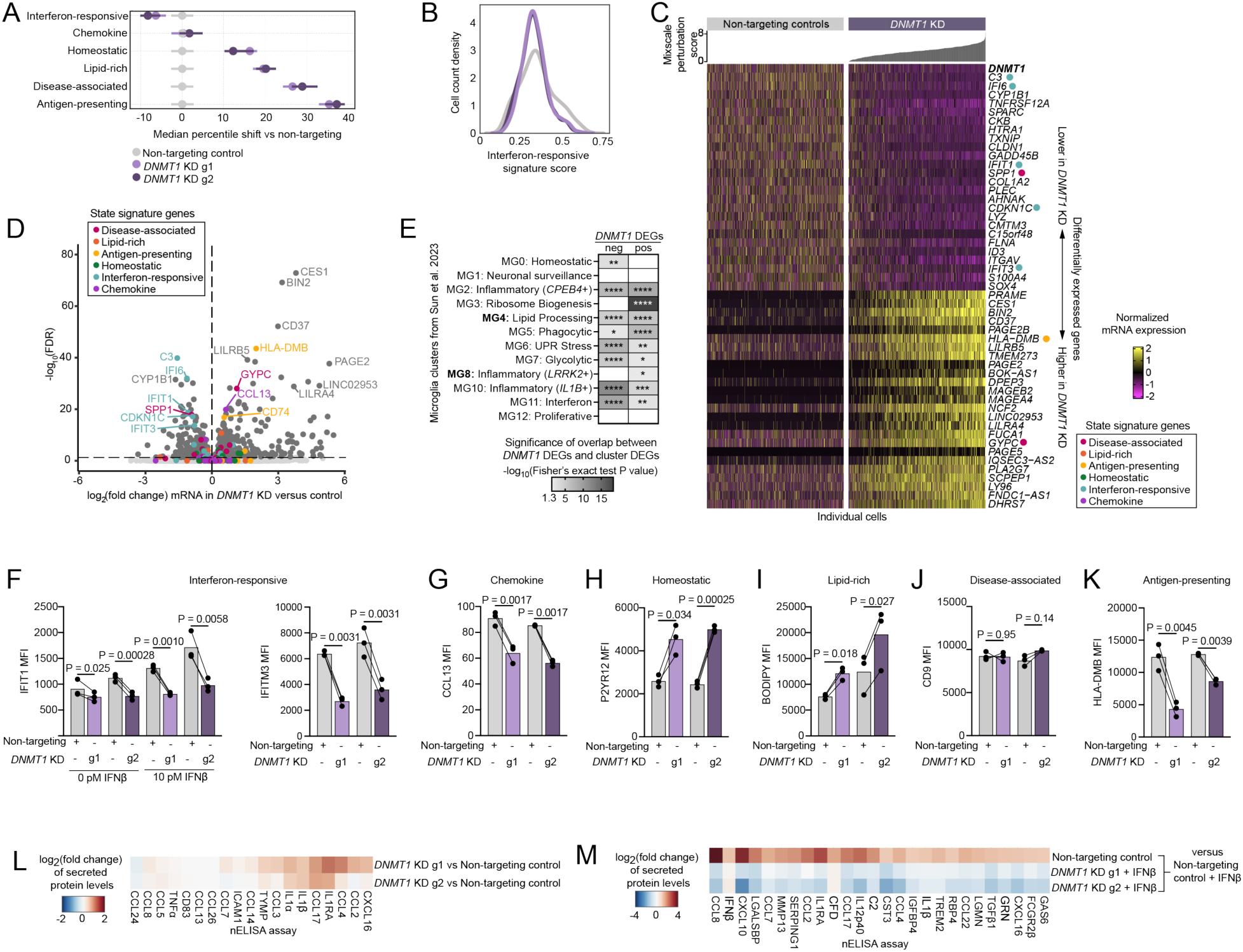
Loss of DNMT1 is generally permissive to activation excluding the interferon-responsive state in iTF-MG. (A) Median percentile shift of state signature scores in *DNMT1* KD (violet) versus non-targeting controls (grey). Separate sgRNAs are represented as g1 and g2. Error bars represent 95% confidence interval. (B) Density plot of interferon-responsive signature score for *DNMT1* KD (violet) versus non-targeting controls (grey). (C) Heatmap of top differentially expressed genes (DEGs) ranked by FDR (25 positive DEGs, 25 negative DEGs). Columns represent individual cells ordered by Mixscale perturbation score (see methods for details). Scores are graphed above the corresponding column (non-targeting control perturbation scores are all zero by definition). Color scale represents the log-normalized mRNA expression. DEGs that appear in our state signature scores are denoted with a colored dot (pink: disease-associated, orange: lipid-rich, yellow: antigen-presenting, green: homeostatic, teal: interferon-responsive, purple: chemokine). (D) Volcano plot of differentially expressed genes in *DNMT1* KD versus non-targeting control. FDR ≤ 0.05. DEGs that appear in our state signature scores colored as above. (E) Heatmap of gene signature overlap analysis of DEGs from *DNMT1* KD with human microglia clusters from Sun et al. 2023^54^. Bolded states are increased in AD patients. Color denotes significance of overlap by Fisher’s exact test P value. Non-significant overlap is white. *P < 0.05, **P < 0.01, ***P < 0.001 ****P < 0.0001. (F-K) Median fluorescence intensity (MFI) of state marker proteins by flow cytometry. *DNMT1* KD microglia (violet) were compared to in-well non-targeting controls (grey) distinguished by nuclear fluorescent proteins. Each point represents an independent well, n ≥ 10,000 cells analyzed per well. P values from paired Student’s T-test. (see also Figure S15) (L) Heatmap displaying differential abundance of secreted proteins that define the chemokine state signature. Differential abundance is shown between DNMT1 *KD* microglia and non-targeting controls. Protein abundance was measured by nELISA. (M) Heatmap displaying differential abundance of secreted proteins measured by nELISA. Protein list displayed represents all differentially abundant proteins in non-targeting control cells after treatment with 10 pM IFNβ for 24 hrs.

The down-regulated DEGs in *DNMT1* KD had significant overlap with multiple human patient-derived microglial states as defined by Sun et al. 2023^54^. The strongest enrichment was for clusters MG10 and MG11, which represent inflammatory and interferon-responsive states (Figure 7E, Table S2). The up-regulated DEGs in *DNMT1* KD also overlapped with many human microglial states as may be expected due to broad hypo-methylation after loss of DNMT1. The strongest overlap was found with cluster MG3, a ribosome biogenesis state (Figure 7E). This state was also enriched for positive DEGs in *STAT2* KD microglia (Figure 6E), so it is possible that this phenotype is related to the inhibition of interferon-response as well.

In two independent *DNMT1* knockdown lines (Figure S2C), we confirmed that *DNMT1* KD inhibits the interferon response using multiple interferon-responsive proteins, IFIT1 and IFITM3 (Figure 7F). We additionally found a decrease in three markers of the chemokine state, which confirms our original screening results (Figure 1B, Figure 7G, Figure S15E) but was not identified using mRNA-level analyses. We also confirmed increased levels of lipid-rich and homeostatic markers, consistent with both the original screens and mRNA analyses (Figure 7H, I). Markers of the DAM state were not consistently altered in *DNMT1* KD (Figure 7J, Figure S15C). Lastly, we found a surprising decrease in antigen-presentation proteins (Figure 7K, Figure S15F). We find these results consistent when knocking down DNMT1 with a small molecule inhibitor, in an additional iPSC line, delaying knockdown until maturation, or in the iMG model (Figure S15N-Y). *DNMT1* KD iTF-MG increased secretion of chemokines in conditioned media compared to control cells (Figure 7L), even though the effect was less strong than with *STAT2* KD. Surprisingly, *DNMT1* KD iTF-MG have a more pronounced effect on limiting interferon-induced cytokine secretion than *STAT2* KD iTF-MG (Figure 7M). *DNMT1* KD iTF-MG also mirror *STAT2* KD iTF-MG with decreased lysosomal pH and Cathepsin B activity (Figure S15G-M). This further supports our finding that loss of tonic interferon signaling impacts lysosomal pH.

### Hypo-methylation in *DNMT1* KD enables expression of interferon-response inhibitors

Although DNMT1 is commonly known as a general DNA methyltransferase, DNMT1 can act in a targeted manner though transcription factor binding^91,92^. To determine if our specific transcriptomic signature is conserved in human patient-derived microglia, we assessed areas of open chromatin in DNMT1-high and DNMT1-low microglia from human postmortem tissue (see methods). We found interferon-response related DEGs from *DNMT1* KD iTF-MG to also be differentially accessible in vivo (Figure 8A, teal), suggesting that the effect of DNMT1 on interferon is specific and conserved.

**Figure 8.**
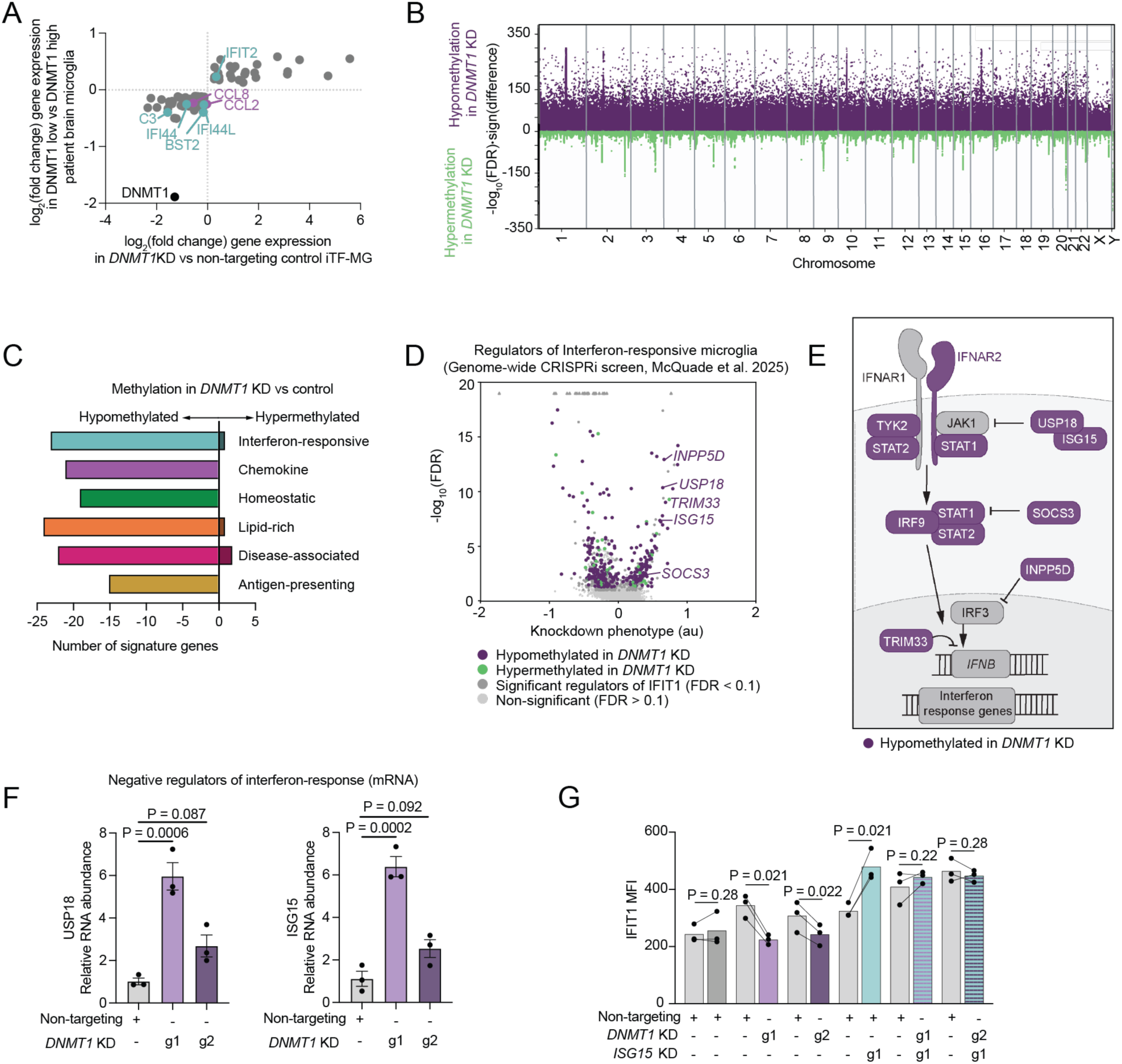
Loss of DNMT1 activates negative regulators of interferon signaling that mediate the interferon-response signature in iTF-MG. (A) Comparison of differentially expressed genes in iTF-MG and differentially accessible chromatin regions from ATAC-seq in human postmortem tissue microglia (see methods). *DNMT1* KD versus non-targeting control iTF-MG is shown on x- axis. Differentially accessible chromatin at gene promoters in human postmortem tissue derived microglia with DNMT1 high vs low (above vs below 70^th^ percentile DNMT1 expression) on y-axis. Genes from interferon-responsive and chemokine state signatures are shown in teal and purple respectively. DNMT1 is colored in black. (B) Manhattan plot showing CpGs of hypo- (purple) and hyper- (green) methylation from enzymatic methylation-sequencing. The x-axis is chromosomal location, and the y-axis is -log_10_(FDR) × sign(difference). (C) Classification of state signature genes for which the promoter region shows hypo- or hyper-methylation after *DNMT1* KD, the x-axis represents count of signature score genes in that category. (D) Volcano plot representing genome-wide CRISPRi screen for interferon-responsive microglia from McQuade et al. 2025^73^. Knockdown phenotype is calculated as a log_2_-ratio of counts in the high and low marker expression populations normalized to the standard deviation of the non-targeting control sgRNAs. Genes for which the promoter region shows hyper- or hypo- methylation are shown in green or purple respectively. Genes with no differential methylation pattern in *DNMT1* KD microglia are shown in grey. (E) Schematic of canonical Type I interferon signaling. Proteins colored in purple are hypo-methylated in *DNMT1* KD microglia and include three negative regulators: USP18, ISG15, SOCS3. (F) Reverse-transcription quantitative PCR analysis of *USP18* and *ISG15*. Points represent delta-delta Ct normalized to expression in non-targeting control cells. Error bars represent standard error. P values from One-way AVONA with Tukey post-hoc testing. (G) Median fluorescence intensity (MFI) of IFIT1 by flow cytometry. non-targeting control microglia (grey), *DNMT1* KD microglia (violet), *ISG15* KD microglia (teal), or *DNMT1* and *ISG15* double knockdown lines (striped) were compared to in-well non-targeting controls (light grey) distinguished by nuclear fluorescent proteins. Each point represents an independent well, n ≥ 10,000 cells analyzed per well. P values from paired Student’s T-test. (see also Figure S15).

To directly investigate the effect of DNMT1 on methylation, we performed whole-genome enzymatic methylation sequencing (EM-seq) in non-targeting control and *DNMT1* KD iTF-MG to identify differentially methylated regions (DMRs) and determine if differential methylation is the mechanism whereby DNMT1 alters microglial activation. As expected, we found robust, widespread hypo-methylation in the *DNMT1* KD microglia (Figure 8B, Table S4). Hypo-methylation was also present at promoters for the majority of genes comprising our state signatures (Figure 8C). This suggests that the decrease of interferon response in *DNMT1* KD is not due to widespread hyper-methylation at interferon response genes.

To nominate targets of DNMT1 that may be responsible for inhibition of the interferon-responsive state, we mapped genes with hypo- or hyper-methylation onto our previous genome-wide CRISPRi screen for interferon-responsive microglia^73^. We found hypo-methylation at promoters of several negative regulators of interferon signaling (including USP18 and ISG15) (Figure 8D, E). Depending on the methylation unit, it is possible for transcription to increase or decrease^93^. We confirmed that *DNMT1* KD microglia show increased mRNA levels of USP18 and ISG15 (Figure 8F). It is possible that *DNMT1* KD unleashes expression of interferon inhibitors leading to the robust decrease in interferon-responsive microglia that we reported. To test this, we generated iTF-MG lines with CRISPRi sgRNAs targeting *DNMT1* and *ISG15* in the same cell. Loss of ISG15 in DNMT1 knockdown background was sufficient to rescue WT interferon response (Figure 8G, Figure S15Z).

## Discussion

In this study, we identified regulators for six microglial activation states through CRISPRi-based functional genomic screens. We perturbed ∼1600 transcription factors and transcriptional regulators in iPSC-derived microglia and sorted microglia on expression of 6 activation state markers. For these large-scale, FACS-based screens, we relied on individual protein markers or dyes to classify microglial activation states. However, one individual protein does not capture the full state signature. Therefore, we selected 31 of these regulators to characterize at a deeper level by multimodal, single-cell analyses. Microglial activation states have been primarily defined at the transcriptomic level as advances in single-cell RNA-sequencing preceded other single-cell modalities. However, we know that there is often divergence between gene expression and protein expression. For this reason, we paired single-cell RNA-sequencing with a CITE-seq library of 166 surface proteins. This allowed us to correlate protein expression with the mRNA-based activation state signatures and uncover new surface markers for each state that we confirmed using our novel state drivers. Many of our state-associated proteins have been reported to be functionally relevant for that state, though future research is needed to understand if the novel correlations we report also contribute to state-specific functions.

We previously developed the iTF-MG microglia differentiation protocol to support large-scale CRISPR screening^57^. This model relies on doxycycline-inducible expression of PU.1, MAFB, CEBPA, CEBPB, IRF5, and IRF8 to directly drive microglial gene expression in twelve days. Importantly, using a directed differentiation approach avoids the bottlenecking events that occur in more complex cytokine-directed protocols that mimic developmental trajectories such as iMG^58^. When differentiating iMG containing the full 9246 element sgRNA library targeting transcription factors and transcriptional regulators, we recorded dropout of > 50% of sgRNAs throughout the differentiation. Thus, we performed the six large scale screens using the iTF-MG model. However, gene expression in the iMG model is known to align more closely with human patient-derived microglia samples. iMG also express higher levels of microglia identity genes and homeostatic markers (Figure 2B, Figure S2) than iTF-MG. Therefore, for our targeted single-cell analyses, we used both iPSC-microglia models in parallel. With this smaller library, the iMG protocol still skewed representation of the sgRNAs from bottlenecking (Table S2), with ∼ 50% of the sgRNAs assigned to less than 100 cells, although we were still able to perform meaningful statistical analyses for the majority of targeted genes by merging sgRNAs at the gene level. This study revealed distinct distributions of activation states present in iTF-MG and iMG. iTF-MG showed higher expression of the chemokine and interferon-responsive states at the population level, while iMG showed higher heterogeneity of disease-associated, lipid-rich, and antigen-presenting states. These results emphasize the importance of selecting the appropriate model system that reflects the biological states relevant to the research.

One outstanding question concerning microglial activation is whether activation states represent mutually exclusive programs or can be modularly combined. Our data supports the idea that individual cells can express multiple state signatures at once. We provide evidence of this using state signature gene programs from the literature or an unbiased scHPF human factor model^80^. For example, we identified many regulators that drive both disease-associated and lipid-rich signatures, but also have evidence that these signatures can move independently. We highlight two regulators of the DAM/lipid-rich states that have opposing levels of antigen-presentation and homeostatic markers as evidence that activation signatures can combine factorially. However, we also reported anti-correlation between the interferon-responsive and chemokine signatures at the mRNA level, suggesting competitively interaction. This type of interaction has been previously reported in our study of inflammatory states in astrocytes^94^. In future work, it will be important to investigate the functional relationships between microglial activation trajectories to understand the underlying molecular circuits that drive or limit activation plasticity. The regulators reported here build a foundational framework that would assist these deeper studies.

Following our multimodal sequencing analyses, we chose four regulators to study in independent knockdown lines: ZNF532, PRDM1, STAT2, and DNMT1. The transcription factor ZNF532 was identified as part of the Human Alzheimer’s Microglia signature with decreased expression in Alzheimer’s disease patients^95^. This direction matches our findings that lower expression of ZNF532 drives the activation states associated with disease. However, our study is the first to report a causal relationship between ZNF532 and DAM gene expression.

PRDM1 is a transcriptional repressor that functions by recruitment of histone deacetylases (HDACs) to target genes. PRDM1 has been most widely studied in B cells where it is required for B cell maturation into plasma cells^96,97^, although PRDM1 is widely expressed across cell types. In microglia, PRDM1 expression is temporally restricted: PRDM1 is absent in microglia during development and activated as part of a postnatal microglia signature^62,98^. Interestingly, three recent studies have identified FDA-approved HDAC inhibitors to mimic neuroprotective, DAM-like signatures^9,99,100^. As *PRDM1* relies on HDAC recruitment to repress targets, it is possible that *PRDM1* loss-of-function explains these findings.

In a study of APOE in iMG, *PRDM1* expression was shown to increase in response to neuronal conditioned media in APOE3/3 but not APOE4/4 microglia. These APOE3/3 microglia were also more responsive to stimuli and increased expression of DAM markers such as GPNMB^101^. PRDM1 was also highlighted as part of a neuroimmune activation module along with the DAM gene *TREM2* in a study of tauopathies^102^. Here, we identified PRDM1 as a functional inhibitor of the DAM state and showed that knockdown of *PRDM1* is sufficient to drive expression of the core DAM signature genes. This makes PRDM1 inhibition an interesting therapeutic potential strategy to drive the DAM state in mature microglia, which is hypothesized to be beneficial for amyloid clearance in the context of Alzheimer’s disease^21,103^. However, we note that the impact of DAM microglia during the tau phase of disease is less clear^104–107^.

The study of PRDM1 and ZNF532 in parallel provides an instructive comparison since knockdown of either drives DAM and lipid-rich states, but each has an opposing effect on antigen-presentation genes. This supports our model that microglia activation signatures can move independently and are best modeled in combinatorial state signatures rather than mutually exclusive, monolithic states. Of interest, we found similar responses to beta-amyloid and synaptosomes in *ZNF532* and *PRDM1* KD, suggesting that the antigen-presenting signature is not critical for this function. Future studies in *in vivo* systems will be required to assess the differential impacts of antigen-presenting-high and antigen-presenting-low DAM on brain health. We provide a well-characterized analysis of *PRDM1* and *ZNF532* KD to support these future studies.

STAT2 is a well-studied transcription factor known to be required for type I interferon responses. Thus, we were not surprised to find that *STAT2* KD decreases interferon response in both iTF-MG and iMG. However, STAT2 has not been widely investigated in microglia specifically, and we report for the first time that inhibition of *STAT2* causes lysosomal defects in iTF-MG. This is an important insight, since blocking microglial interferon responses is currently being investigated as a therapeutic mechanism for Alzheimer’s disease^35^. We also report a strong decrease in APOE expression in *STAT2* KD microglia, an important finding given that APOE is one of the strongest risk factors for Alzheimer’s disease^108^

DNMT1 is a DNA methyltransferase primarily responsible for maintaining DNA methylation patterns during replication. Interestingly, DNMT1 has been shown to be necessary for hematopoietic development. Global *DNMT1* knockout mice are not viable, and *DNMT1* knockout specifically in the hematopoietic system leads to deficient maturation of hematopoietic stem cells^109^. Of note, from the five independent sgRNAs targeting *DNMT1* used in our TF-wide screens, only three were present at the end of differentiation. It is possible that the two sgRNAs that dropped out had stronger knockdown of *DNMT1*, reducing viability. Indeed, the two sgRNAs targeting *DNMT1* that were used in our follow-up studies have a modest ∼50% knockdown of *DNMT1* and no toxicity.

Dysregulation of DNA methylation is a hallmark of aging and is known to contribute to age-related phenotypes including cognitive decline and AD^110–112^. In patients with Alzheimer’s disease, significant DNA hyper-methylation is specifically found in microglia ^113^. However, the impact of this hyper-methylation has not been investigated. Our functional genomic screens have identified that perturbing regulators of DNA methylation has a strong impact on microglial activation states. We specifically followed up on DNMT1, but determined that other regulators of DNA methylation also modify microglial activation (ex. TET2, SETDB1, SETDB2, CXXC4, MBD1, MBD6, Table S1). Guided by these results, we hypothesize that skewed microglial activation could explain the aging phenotypes correlated with altered DNA methylation in these microglia. Our study found that hypo-methylation reduced interferon-responsive microglia through activation of negative regulators of interferon responses. Further research is needed to understand if hyper-methylation increases interferon-responsive microglia. Since interferon-responsive microglia are known to remove synapses^30,34^, enriching this activation state could explain the correlation between hyper-methylation and Alzheimer’s disease. Additionally, a progressive neurodegenerative disease known as Hereditary sensory and autonomic neuropathy type 1E (HSAN1E) is caused by autosomal dominant mutations in DNMT1^114^. Microglia contributions to this disease have not yet been assessed. DNMT1 mutations have also been linked to Autosomal dominant cerebellar ataxia, deafness and narcolepsy (ADCA-DN) and with related mutations in P2RY11 and HLA-DQB1∗ 06:02^115^. These results are of particular interest given our findings that DNMT1 impacts both homeostatic markers and antigen-presenting markers in microglia. Microglia activation has not been assessed in ADCA-DN.

An important property of CRISPR interference is that knockdown populations capture a spectrum of perturbation rather than the binary on/off signal from CRISPR knockout. This is beneficial as it reduces lethality that may occur from the complete loss of important transcription factors. We additionally leveraged this feature by applying Mixscale^87^ to rank cells based on perturbation strength. This allows us to infer perturbation for genes that are expressed at too low a level to be detected by single-cell RNA-sequencing. Stratification of phenotypes by Mixscale perturbation scores revealed interesting relationships between our regulators of interest and their downstream changes in gene expression. For ZNF532, we found a binary switch where minimally perturbed cells showed no shifts of state signatures while moderately perturbed cells showed nearly the same extent of state signature shifts as highly perturbed cells. In contrast, *PRDM1* KD shows a more gradual change in state signature expression with increasing perturbation, suggesting that the impact of PRDM1 on the DAM and lipid-rich states may be tunable. We discovered a set of down-regulated genes (S100A9, FCGBP, RFLNB) that are significantly changed even in cells with low Mixscale perturbation scores. We hypothesize that these genes may be direct targets inhibited by PRDM1. We also discovered a set of positive DEGs that turn on exclusively in cells with the highest perturbation scores (SCIN, MMP2, NUPR1). Many of our DAM genes also follow this trajectory, with uniformly higher expression in the most highly perturbed cells. For *STAT2* KD cells, we observed a more binary response for inhibition of interferon, DAM states have a gradual increase across Mixscale scores, and changes in the chemokine signature are only present in the most highly perturbed *STAT2* KD cells. *DNMT1* KD is unique in that minimally, moderately, and maximally perturbed cells produce very similar state signature shifts. This may reflect the nature of widespread DNA hypo-methylation.

To contextualize our perturbations in iPSC-derived microglia with human brain-derived microglia, we performed overlap analysis of our differentially expressed genes from *ZNF532, PRDM1*, *STAT2,* and *DNMT1* knockdown with the human patient-derived microglia clusters identified in Sun et al. 2023^54^. In the original publication, the authors predicted regulators of their microglial clusters based on transcription factor motif enrichment analysis. They predicted STAT2 to be a regulator of MG11, an interferon-responsive cluster. Our data confirms that STAT2 does indeed drive the genes defining this cluster. In contrast, the authors predicted PRDM1 to regulate clusters MG6 and MG7, but our data shows that PRDM1 DEGs only overlap with clusters MG3, MG4, MG5, and MG11. Instead, MG4, a lipid-processing state, shows the most significant overlap with *PRDM1* KD DEGs. Interestingly, their TF-enrichment analysis did not uncover any significant regulators for MG3, a ribosome biogenesis cluster. Our data reveals *DNMT1* to be a strong regulator of this signature. Depending on the database, DNMT1 would not be represented with a searchable DNA motif that can be utilized in these types of TF motif enrichment analyses, highlighting the benefit of our CRISPR screening approach and a broader focus on epigenetic regulators of gene expression beyond transcription factors.

To further understand the discrepancies between predicted TF targets and the DEGs discovered here by TF perturbation, we generated a gene regulatory network from our own CROP-seq dataset and quantified the overlap of predicted gene targets and actual DEGs (Table S2). For well-described TFs like STAT2 and IRF9, we find approximately half of the DEGs described here are predicted targets of STAT2. The other half may represent microglia-specific targets of STAT2, or downstream effects of *STAT2* KD past the direct loss of TF binding. For other TFs including PRDM1 and DNMT1 we find almost no overlap between predicted targets and DEGs. Further work is required to understand whether this is caused by microglia specific co-factors or effects downstream of TF-binding. ZNF532 does not have a predicted binding motif in human, and thus could not be analyzed in this way further highlighting the need for functional perturbation experiments as presented here.

We also aligned our single-cell sequencing data to a recent factor-based model of microglia activation states derived from human microglia across a compendium of brain areas and diseases^80^ (Table S2). We found high levels of the GPNMB^high^ factor in *ZNF532* and *PRDM1* KD iMG. This factor is positively correlated with pathological AD, amyloid, and tau tangles across meta-analysis of 4 large datasets. As future studies align to a common factor-based model, we will gain a better understanding of the significance of these factors in brain health and microglia function.

Together, our data offer a unique lens to study microglial activation states. We importantly use multi-modal analyses of state perturbations at the mRNA, protein, and functional level and discuss key cases in which there is misalignment between these modalities. This data strongly highlights the need to validate findings past the mRNA profiling stage and cautions against assuming downstream function based solely on mRNA expression.

### Limitations of the study

One limitation of this study is the use of iPSC-derived microglia models. In the brain, microglia are constantly signaling to their surrounding cells and receiving feedback from their environment. Microglial activation occurs and is resolved as a response to these complex combinatorial signals. By using iPSC-derived microglia in monoculture, we are not modeling this intricate environment. However, this platform provides the highest level of control and removes the confounding variables of environmental homeostatic or proinflammatory signals that may occur in vivo. Because the regulators we identified act through transcriptional regulation, we expect faithful translation in vivo is likely. However, further work will be needed to test if our regulators can override environmental signals.

Another limitation is that we were unable to perform the high-resolution single-cell analyses on the full ∼1600 gene sgRNA library of transcription factors and transcriptional regulators due to the substantial cost of these experiments. For this reason, we opted to use a tiered approach with six unbiased screens for the ∼1600 gene library, paired CROP-seq and CITE-seq for 31 hits nominated from the larger library, and detailed analysis of four genes that were particularly interesting by single-cell sequencing. While this approach ensured that our deeper characterizations focused on particularly robust regulators of microglial activation states, our data is necessarily limited in scope by the high cost of transcriptomic analysis.

## Resource availability

### Lead contact and materials availability

The cell lines generated in this study are available on request upon the completion of a Material Transfer Agreement (MTA). Plasmids generated in this study will be deposited on AddGene. Further information and requests for resources and reagents should be directed to and will be fulfilled by the Lead Contact, Martin Kampmann.

### Data and code availability

Raw single-cell CROP-seq and CITE-seq data will be made available on Gene Expression Omnibus (GEO). Processed data and additional metadata will be made available by author request.

CRISPR-screening data will be posted on www.CRISPR-brain.org. Raw files will be made available upon request.

The code for the custom Mixscale (https://github.com/satijalab/Mixscale) analysis of using Benajmini-Hochberg correction in comparison to the standard Bonferroni correction can be found on Github (https://github.com/reetm09/mixscale). All remaining Python and R code for analysis and for figure generation is also available on Github (https://github.com/reetm09/mglia_regulators_paper).

## Acknowledgements

We acknowledge all members of the Kampmann lab for their technical expertise and advice. We thank the Innovation Core at the Weill Institute for Neurosciences for sharing their microscopy expertise and the Laboratory for Cell Analysis for sharing flow cytometry and FACS expertise. We also thank Steven Boggess for advice on CROP-seq sample preparation and Ya Zhang at Columbia for advice on the CITE-seq sample preparation. The UCSF Neurodegenerative Disease Brain Bank receives funding support from NIH grants P30AG062422, P01AG019724, U01AG057195, and U19AG063911, as well as the Rainwater Charitable Foundation and the Bluefield Project to Cure FTD. This work was supported by Alzheimer’s Association grants AARF-22-973222 (AM), ZEN-22-969903 (MK), and ADSF-21-831212-C (MK), Larry L. Hillblom Foundation grant 2022-A-016-FEL (AM), Innovative Genomics Institute Grant CA-0204195 (MK), CIRM EDUC2-12730 (TT), 1R01AG075802-01 (J.E.R), Rainwater Charitable Foundation (J.E.R), 1R01NS144499(J.E.R), National Science Foundation Graduate Research Fellowship under Grant No. 2034836 (OT) and Grant No. 17528 (JL), and Chan Zuckerberg Initiative grant CP2-1-0000000332 (MK). SEC is a Chan Zuckerberg Biohub Investigator. JKN acknowledges funding for this work from the National Institutes of Health (R35GM155044) and the Chan Zuckerberg Biohub San Francisco.

## Author contributions

A. McQuade, R. Mishra, and M. Kampmann designed and conducted and analyzed the overall research, created figures, and wrote the manuscript. A. McQuade, VC. Castillo, and V. Hagan performed the CRISPRi pooled screens. VC. Castillo and W. Liang analyzed the pooled CRISPRi screens. V. Hagan cloned the CROP-seq library. A. McQuade performed the CROP-seq/CITE-seq screens with advice from V. Haage. R. Mishra analyzed the CROP-seq and CITE-seq screens. R. Mishra, M. Fujita, and V. Marshe performed protein analysis of the CITE-seq data. N. Robichaud performed nELISA. O. Teter, V. Hagan, B. Gonzalez, and W. Liang generated individual sgRNA lines. L. Gomes performed RT-qPCR of DNMT1 targets. A. McQuade, O. Teter, and T. Ta updated the differentiation protocol. A. McQuade, V. Hagan, B. Gonzalez, and W. Liang performed functional studies in iTF-MG model. A. McQuade performed functional studies in iMG model. X. Han and J. Rexach analyzed human data. A. McQuade and P. Colias performed EM-seq. P. Colias, J. Lubin, and R. Mishra analyzed the EM-seq data under supervision from S. Chasins and J. Nuñez. All authors reviewed and approved the final manuscript.

---

**Figure S1.**
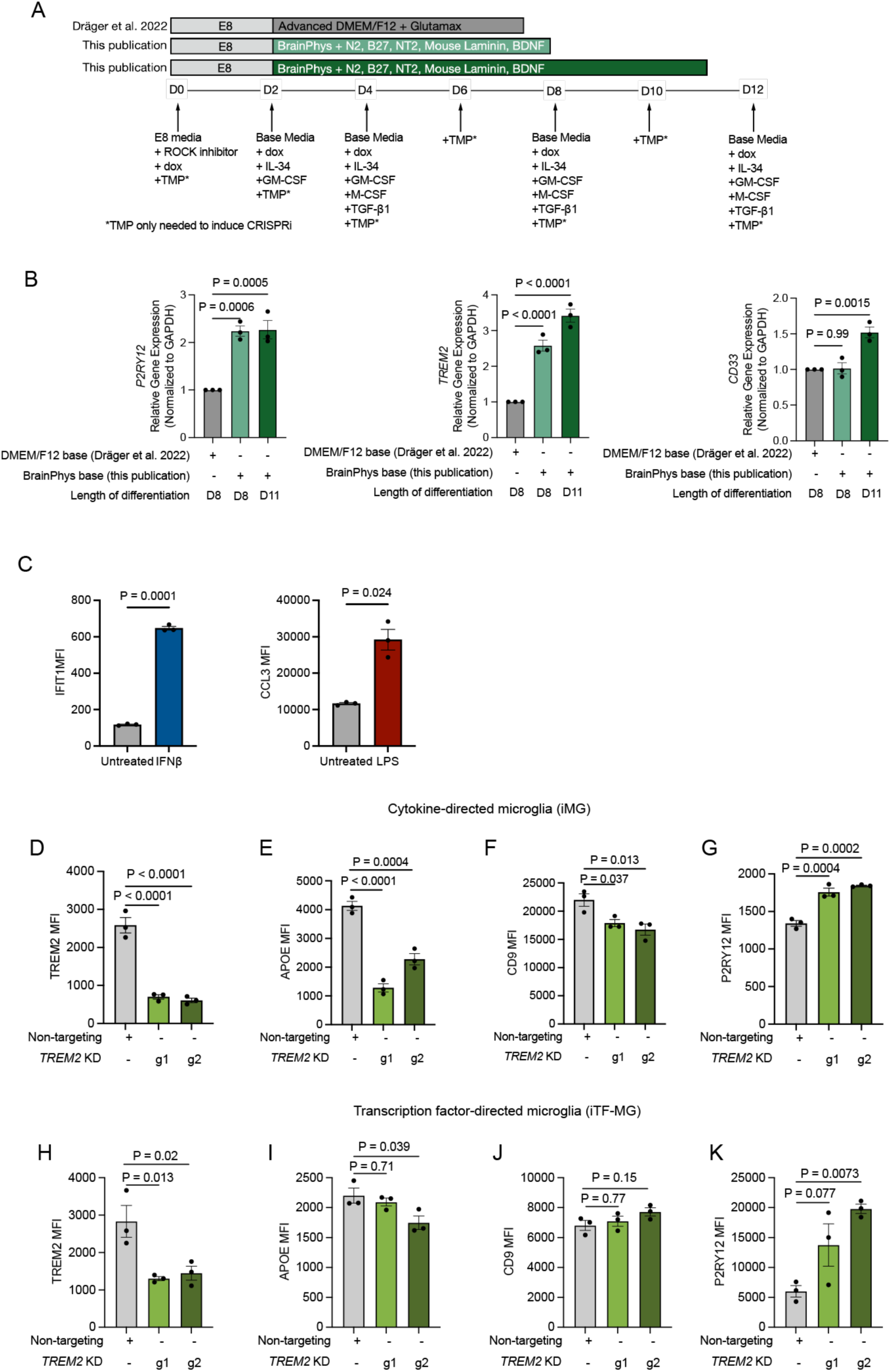
Updated differentiation protocol and perturbation of TREM2 in iMG and iTF-MG. (A) Schematic of the original and updated media composition and differentiation lengths. (B) Reverse-transcription quantitative PCR analysis of homeostatic microglia markers (P2RY12, TREM2, CD33) in the original and updated media compositions. P value one-way ANOVA with Tukey post-hoc testing. (C) Stimulation of iMG model with 50 pM IFNβ or 100 ng/mL LPS is sufficient to drive additional activation states that are not present at baseline. Median fluorescence intensity (MFI) of state marker proteins by flow cytometry 24 hours after stimulation. Points represent one well, n ≥ 10,000 cells analyzed per well. P value students T test. (D-K) iPSC-Microglia with CRISPRi targeting *TREM2* were differentiated using the iMG protocol (D-G) or iTF-MG protocol (H-K). Median fluorescence intensity (MFI) of state marker proteins for DAM (TREM2, CD9, APOE) and homeostatic (P2RY12) states by flow cytometry. *TREM2* KD microglia (green) were compared to non-targeting controls (grey). Points represent one well, n ≥ 10,000 cells analyzed per well. P value is One-way ANOVA.

**Figure S2.**
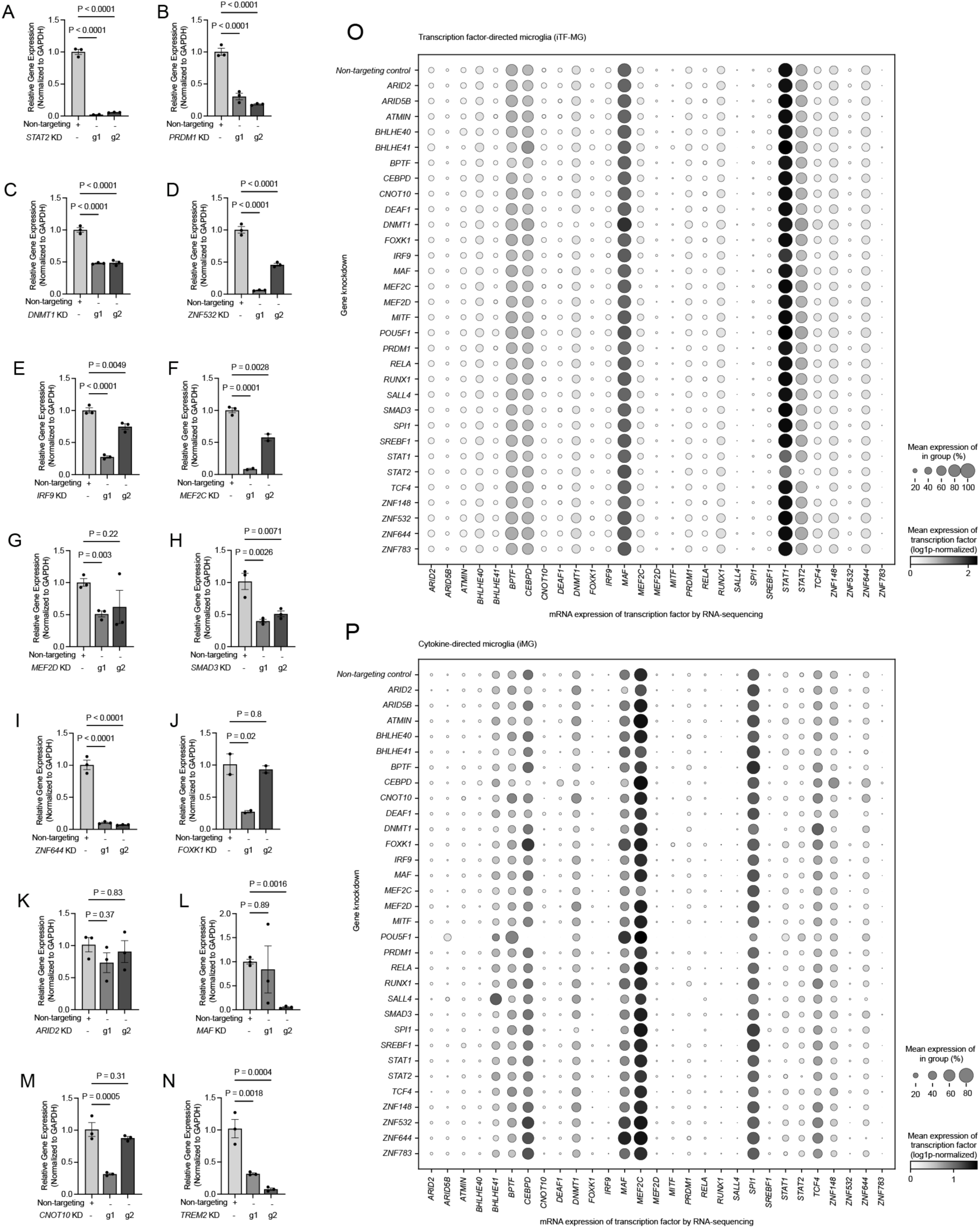
Gene expression knockdown validation for key sgRNAs. (A-N) Reverse-transcription quantitative PCR analysis of targeted genes after induction of CRISPRi. P value one-way ANOVA with Tukey post-hoc testing. *STAT2, DNMT1, IRF9, CNOT10, SMAD3, and TREM2* were performed in iTF-MG. *PRDM1, MEF2C, MEF2D, ZNF644, FOXK1, ARID2, ZNF532,* and *MAF* were quantified in iMG. (O,P) Mean mRNA expression of all genes perturbed in CROP-seq analysis separated by gene perturbation in (O) iTF-MG and (P) iMG. Columns represent the mean normalized gene expression. Rows represent the perturbed gene knocked down with CRISPRi. Each gene row is the average of two sgRNAs. The non-targeting control represents the average of four sgRNAs. Circle color represents mean expression of mRNA. Circle size represents the percentage of cells expressing the gene.

**Figure S3.**
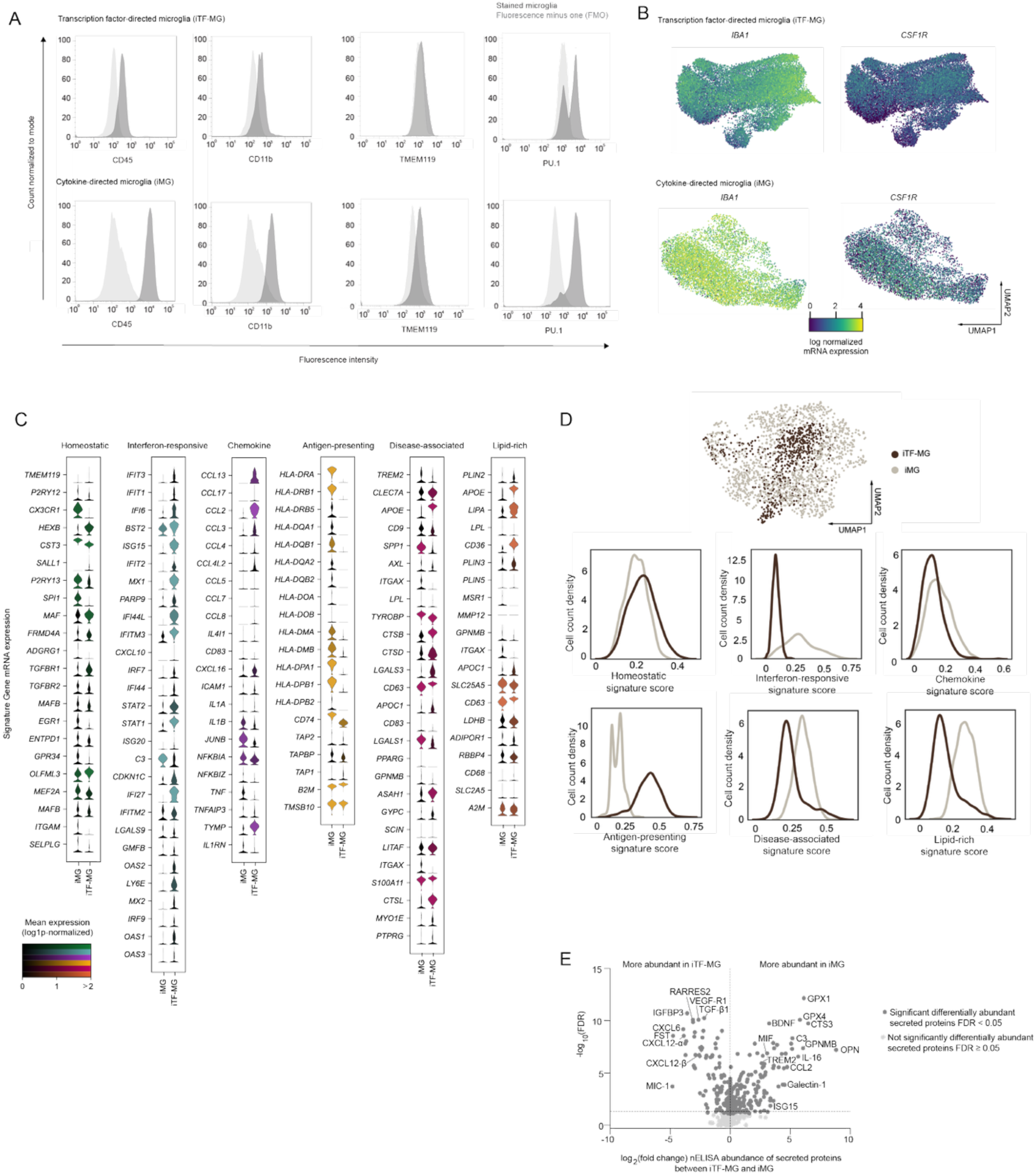
Expression of immune and microglia proteins in iTF-MG and iMG. (A) Histograms representing expression of relevant immune markers by flow cytometry in iTF-MG (top) and iMG (bottom). Antibody-stained cells (dark grey) are compared to a fluorescence minus one control (FMO, light grey) of the same cell type. Data shown from one representative well. Three wells analyzed per marker n ≥ 10,000 cells analyzed per well. (B) UMAP representation of iTF-MG (top) and iMG (bottom) mRNA space. Cells are colored by expression levels of *IBA1* and *CSF1R*. (C) Violin plots of microglia activation state signature genes in non-targeting control cells from iMG versus iTF-MG. Color intensity represents mean expression over all cells (log1p-normalized). (D) Integration of non-targeting control cells from iTF-MG (dark brown) and iMG (grey). UMAP representation of iMG and iTF-MG integration (top). Signature score gene expression in iMG and iTF-MG (bottom). iTF-MG has higher heterogeneity over interferon-responsive and chemokine signature scores. iMG has higher heterogeneity over antigen-presenting, disease-associated, and lipid-rich signature scores. (E) Volcano plot representing differential abundance of secreted proteins measured by nELISA. Positive fold change values are more abundant in conditioned media from iMG. Negative fold change values are more abundant in conditioned media from iTF-MG. n=6 wells iTF-MG, n=8 wells iMG. 10,000 cells per well. Media was conditioned for 24 hours.

**Figure S4.**
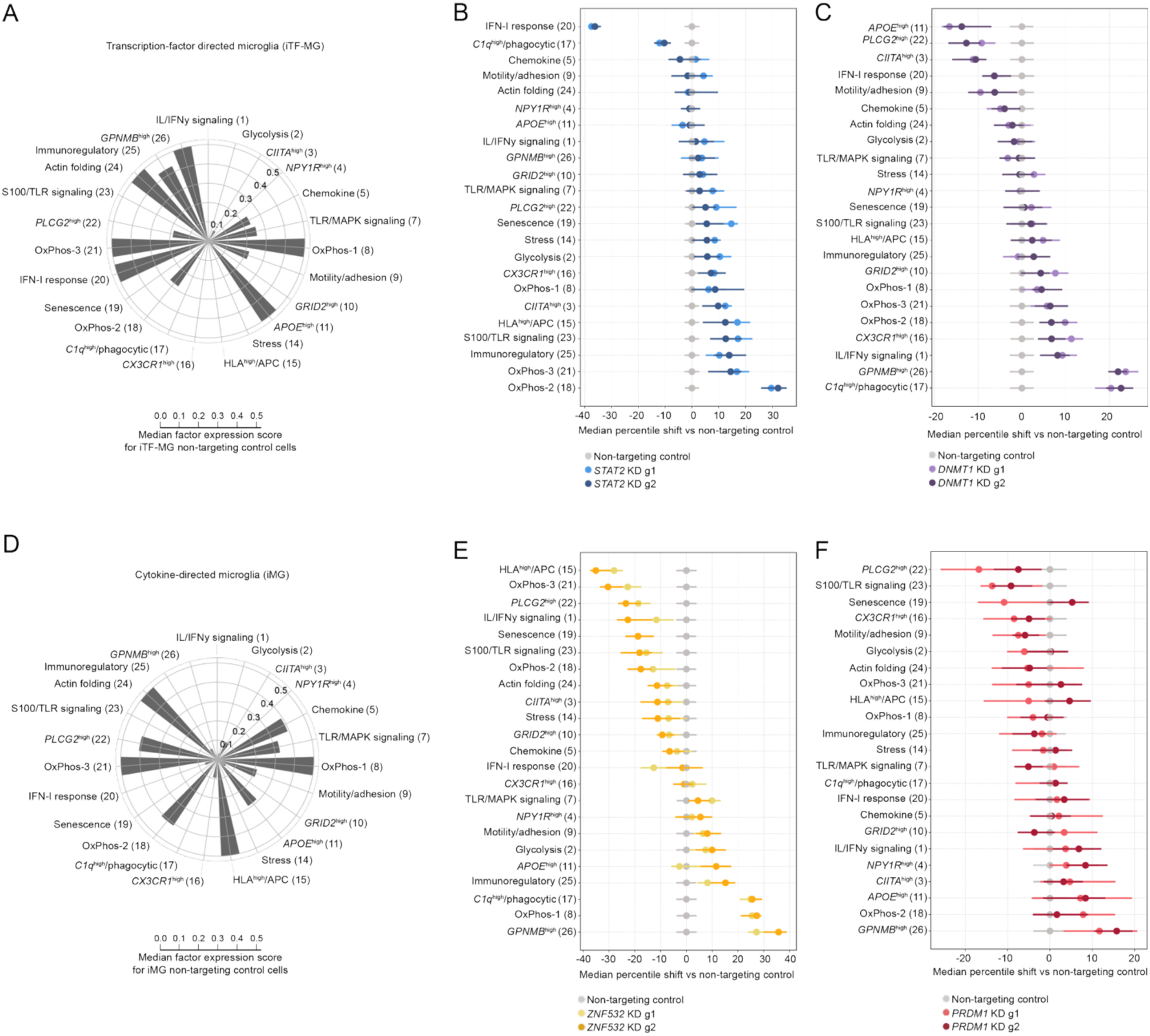
scHPF 26-factor model from human brain-derived microglia. Alignment of single-cell analysis to scHPF factor model from Marshe et al. 2025. (A) Median factor expression for each factor in iTF-MG non-targeting control cells. (B, C) Median percentile shift of state signature scores in (B) *STAT2* KD (blue) or (C) *DNMT1* KD (violet) versus non-targeting control iTF-MG (grey). Separate sgRNAs are represented as g1 and g2. Error bars represent 95% confidence interval. (D) Median factor expression for each factor in iMG non-targeting control cells. (B, C) Median percentile shift of state signature scores in (E) *ZNF532* KD (yellow) or (F) *PRDM1* KD (salmon, red) versus non-targeting control iMG (grey). Separate sgRNAs are represented as g1 and g2. Error bars represent 95% confidence interval. See also table S2.

**Figure S5.**
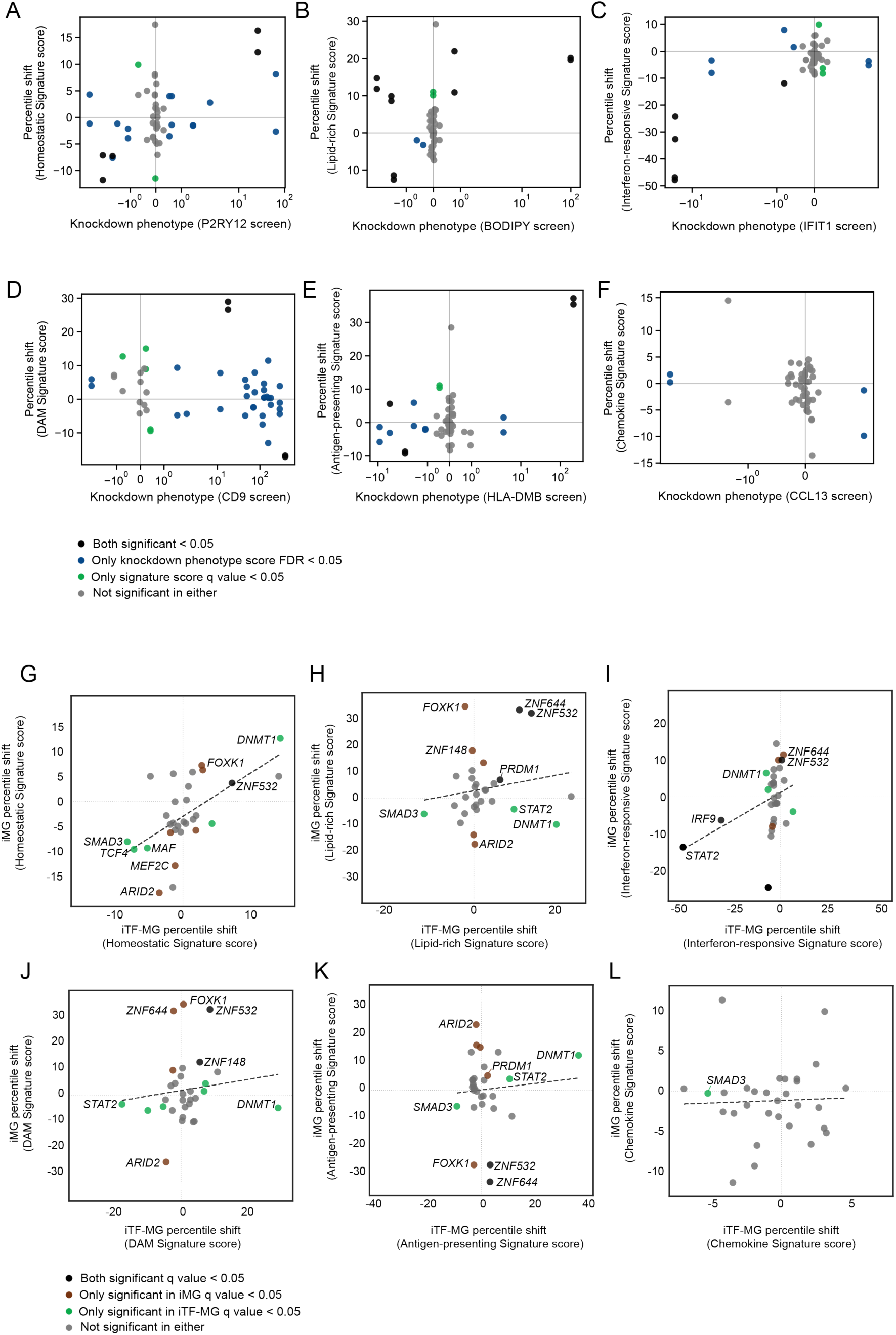
Cross-screen comparisons. (A-F) Correlation of FACS-based screens and signature scores from CROP-seq based screen. X-axis represents the knockdown phenotype from FACS-based screens in Figure 1. Knockdown phenotype is calculated as a log_2_ ratio of counts in the high and low marker expression populations normalized to the standard deviation of non-targeting control cells. Y-axis represents the median percentile shift of state signature scores from CROP-seq in each knockdown versus non-targeting controls. Individual points represent a specific guide RNA. If the shift in FACS-based phenotype score is significant (FDR < 0.05), points are colored blue. If the shift in CROP-seq based signature scores is significant (q value < 0.05), points are colored green. If both are significant, points are colored black. If neither are significant, points are colored grey. (G-L) Correlation of CROP-seq based screens in iTF-MG (x-axis) and iMG (y-axis). Data is presented as percentile shift of state signature scores from CROP-seq in each knockdown versus non-targeting controls. Individual points represent a perturbed gene (average of 2 gRNAs). If the shift in iTF-MG signature scores is significant (q value < 0.05), points are colored green. If the shift in iMG signature scores is significant (q value < 0.05), points are colored brown. If both are significant, points are colored black. If neither are significant, points are colored grey.

**Figure S6.**
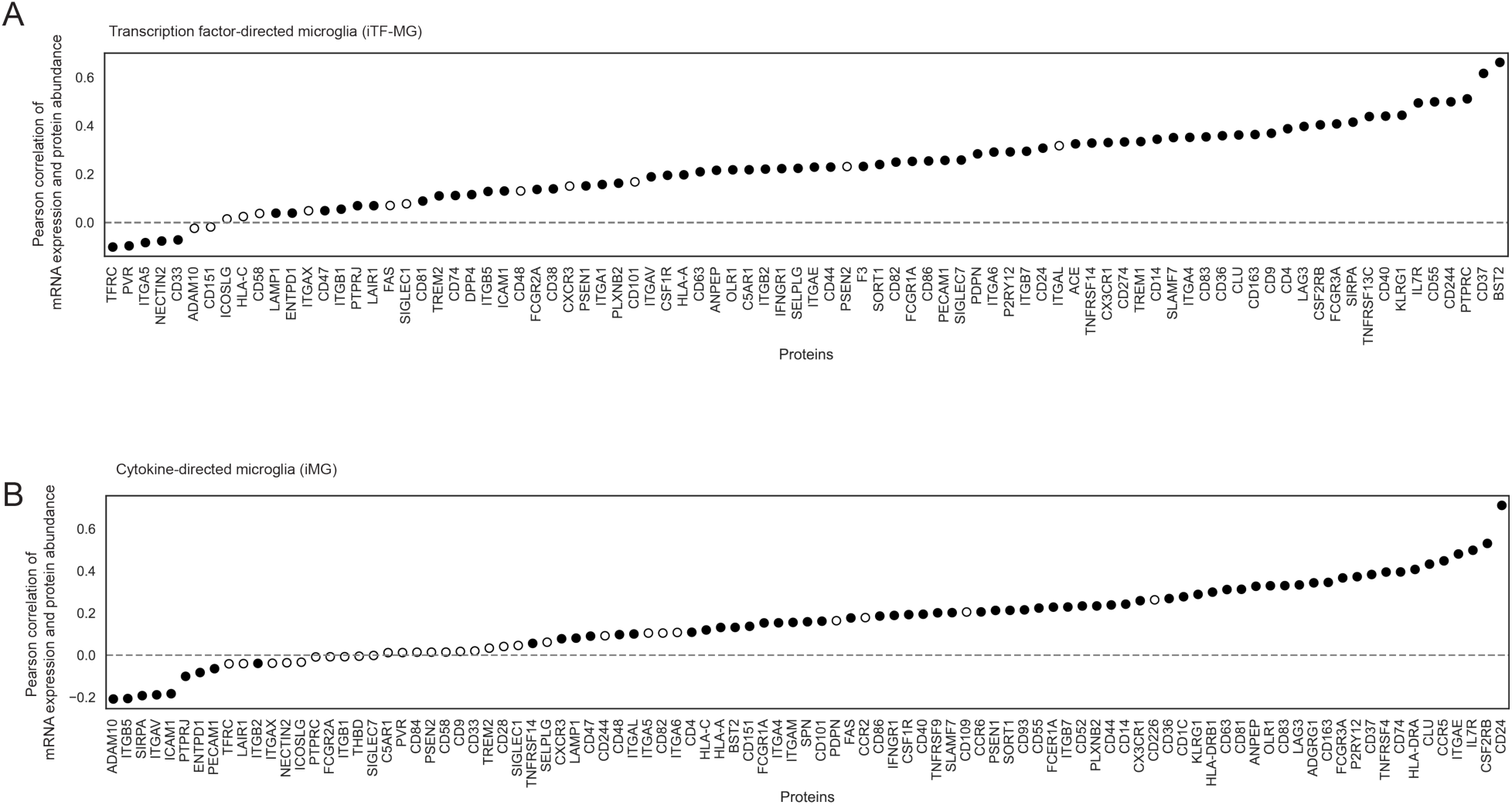
Correlation of mRNA and protein from paired CROP-seq and CITE-seq dataset. Pearson correlation of mRNA expression and protein abundance per cell in all non-targeting control cells from (A) iTF-MG or (B) iMG. Filled circles represent significant correlation. Open circles represent non-significant correlation. Adjusted p value from Pearson correlation P < 0.05.

**Figure S7.**
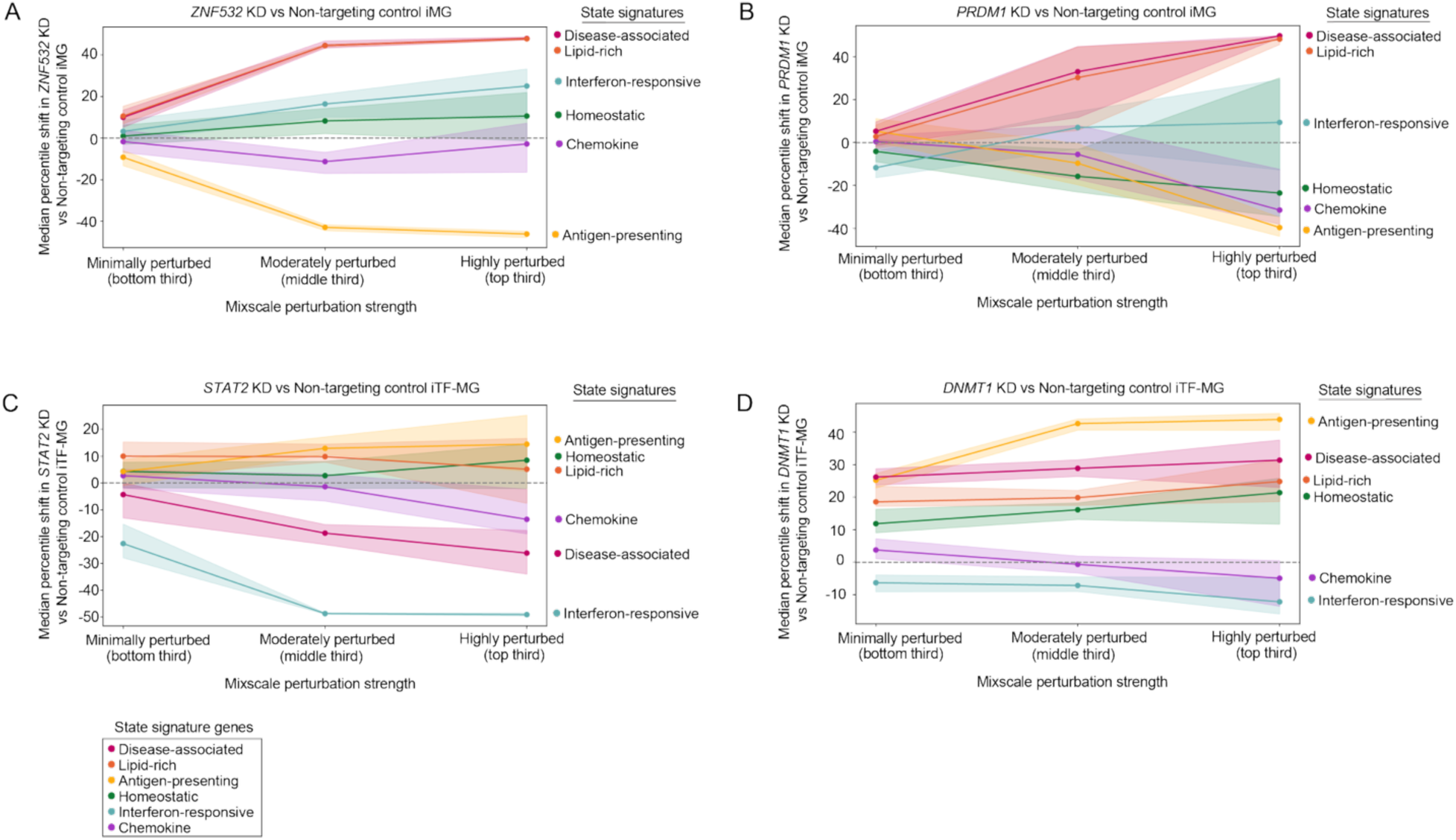
Stratification of signature score enrichment by perturbation strength. Median percentile shift of state signature scores in (A) *ZNF532* KD vs non-targeting control iMG (B) *PRDM1* KD vs non-targeting control iMG (C) *STAT2* KD vs non-targeting control iTF-MG (D) *DNMT1* KD vs non-targeting control iTF-MG. For each perturbation, cells were stratified by their Mixscale perturbation score into the bottom, middle, and top third of perturbation scores (see methods; Cell counts per grouping (minimally, moderately, highly perturbed) *ZNF532* KD: 432, 217, 78; *PRDM1* KD: 446, 82, 13; *STAT2* KD: 373, 252, 76; *DNMT1* KD: 718, 649, 58). The median percentile shift for each state signature was calculated for each perturbation strength against the non-targeting control cells (iMG: 637 cells; iTF-MG: 1311 cells) and plotted with a 95% confidence interval. State signatures are shown in pink: disease-associated, orange: lipid-rich, yellow: antigen-presenting, green: homeostatic, teal: interferon-responsive, purple: chemokine. See also table S2.

**Figure S8.**
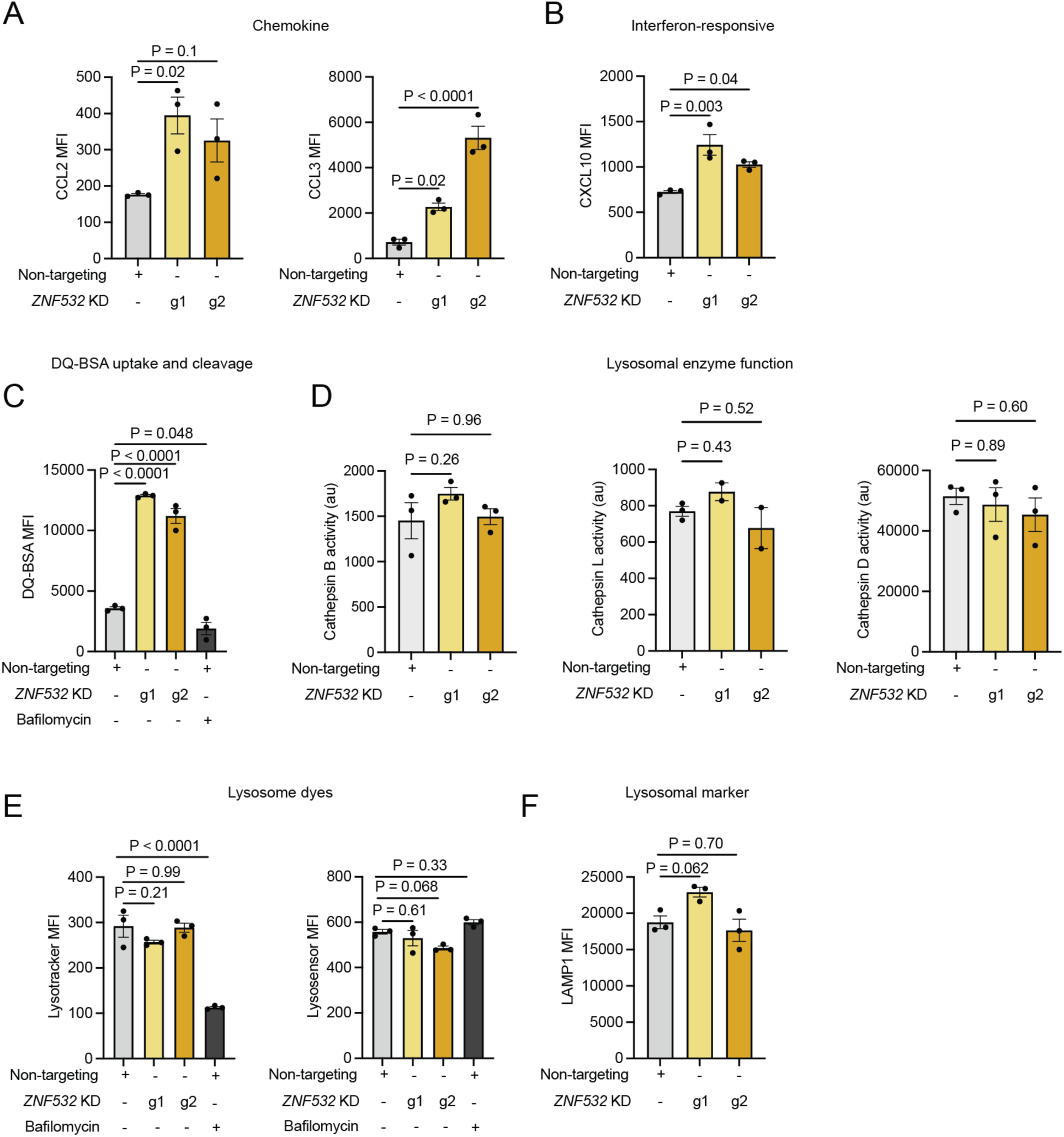
Quantification of additional protein state markers and lysosomal function in *ZNF532* KD. (A-B) Median fluorescence intensity (MFI) of state marker proteins by flow cytometry. *ZNF532* KD microglia (yellow) were compared to non-targeting controls (grey). Points represent one well, n ≥ 10,000 cells analyzed per well. One way ANOVA. (C) Median fluorescence intensity (MFI) of DQ-BSA by flow cytometry. DQ-BSA was added for two hours before analysis to allow for uptake. Bafilomycin A was added four hours before analysis to control wells. Points represent one well, n ≥ 10,000 cells analyzed per well. One way ANOVA. (D) Cathepsin activity measured in *ZNF532* KD microglia (yellow) and non-targeting control cells (grey). Points represent independent wells, n ≥ 70,000 cells. One way ANOVA. (E) Median fluorescence intensity (MFI) of lysotracker and lysosensor in *ZNF532* KD microglia (yellow) and non-targeting controls (grey). Bafilomycin A was added four hours before analysis to control wells. Points represent one well, n ≥ 10,000 cells analyzed per well. One way ANOVA. (F) Median fluorescence intensity (MFI) of LAMP1 by flow cytometry. *ZNF532* KD microglia (yellow) were compared to non-targeting controls (grey). Points represent one well, n ≥ 10,000 cells analyzed per well. One way ANOVA.

**Figure S9.**
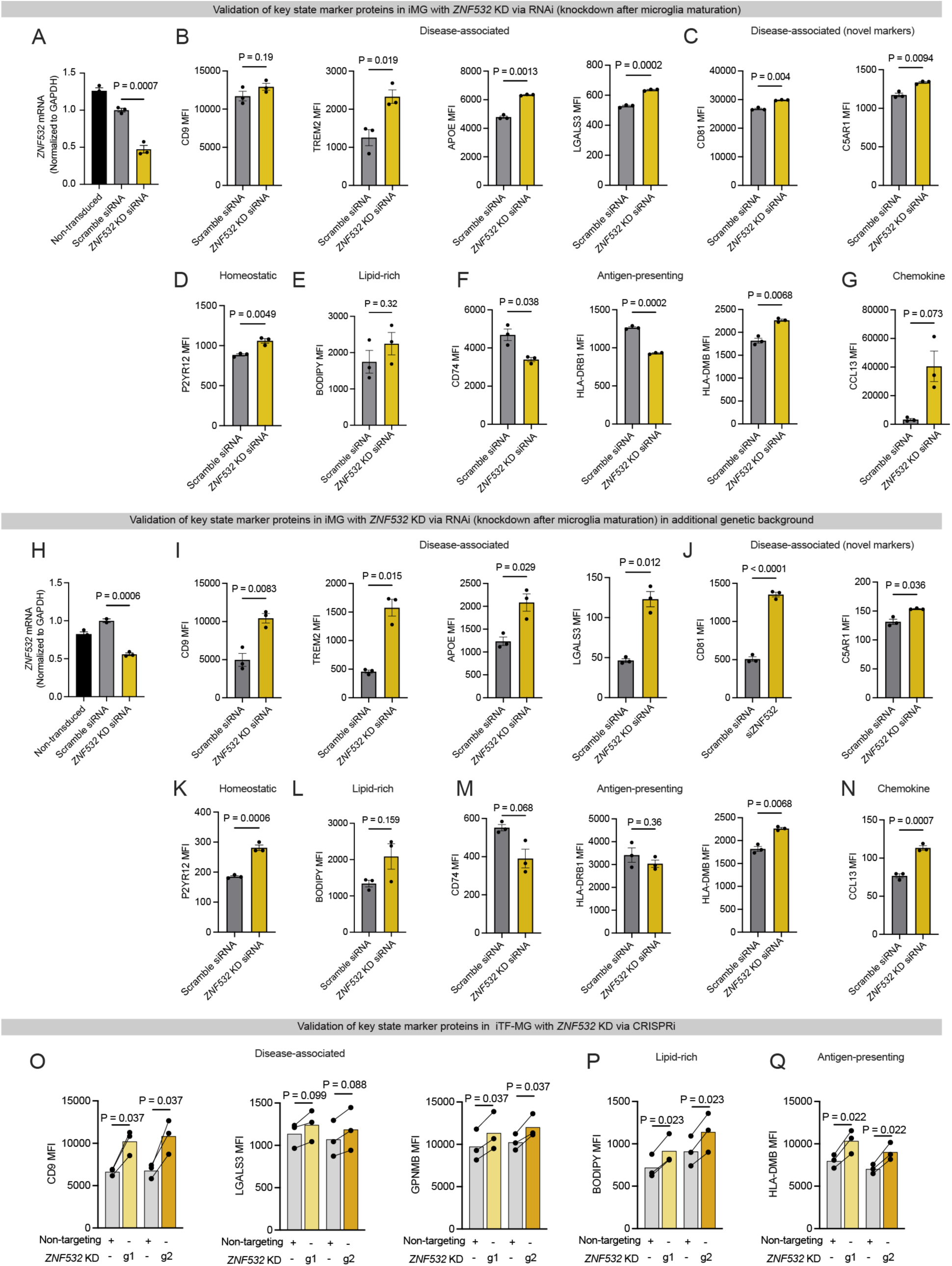
Validation of ZNF532 phenotypes using alternate methods, timing, patient backgrounds, and differentiation protocol. (A) Reverse-transcription quantitative PCR analysis of *ZNF532* expression after siRNA knockdown in WTC11 background. Cells not transduced with siRNA (black), transduced with scramble siRNA (grey) and *ZNF532* KD with siRNA (yellow). P value from one-way ANOVA. (B-G) Median fluorescence intensity (MFI) of state marker proteins by flow cytometry. *ZNF532* KD with siRNA (yellow) and scramble siRNA (grey). Points represent one well, n ≥ 10,000 cells analyzed per well. P value T-test. (H) Reverse-transcription quantitative PCR analysis of *ZNF532* expression after siRNA knockdown in KOLF2.1J background. Cells not transduced with siRNA (black), transduced with scramble siRNA (grey) and *ZNF532* KD with siRNA (yellow). P value from one-way ANOVA. (I-N) Median fluorescence intensity (MFI) of state marker proteins by flow cytometry. *ZNF532* KD with siRNA (yellow) and scramble siRNA (grey). Points represent one well, n ≥ 10,000 cells analyzed per well. P value T-test. (O-Q) *ZNF532* KD in iTF-MG differentiation protocol. Median fluorescence intensity (MFI) of state markers by flow cytometry. *ZNF532* KD iTF-MG microglia (yellow) are compared to in-well non-targeting controls (grey). Points represent one well, n ≥ 10,000 cells analyzed per well. P value paired T-test.

**Figure S10.**
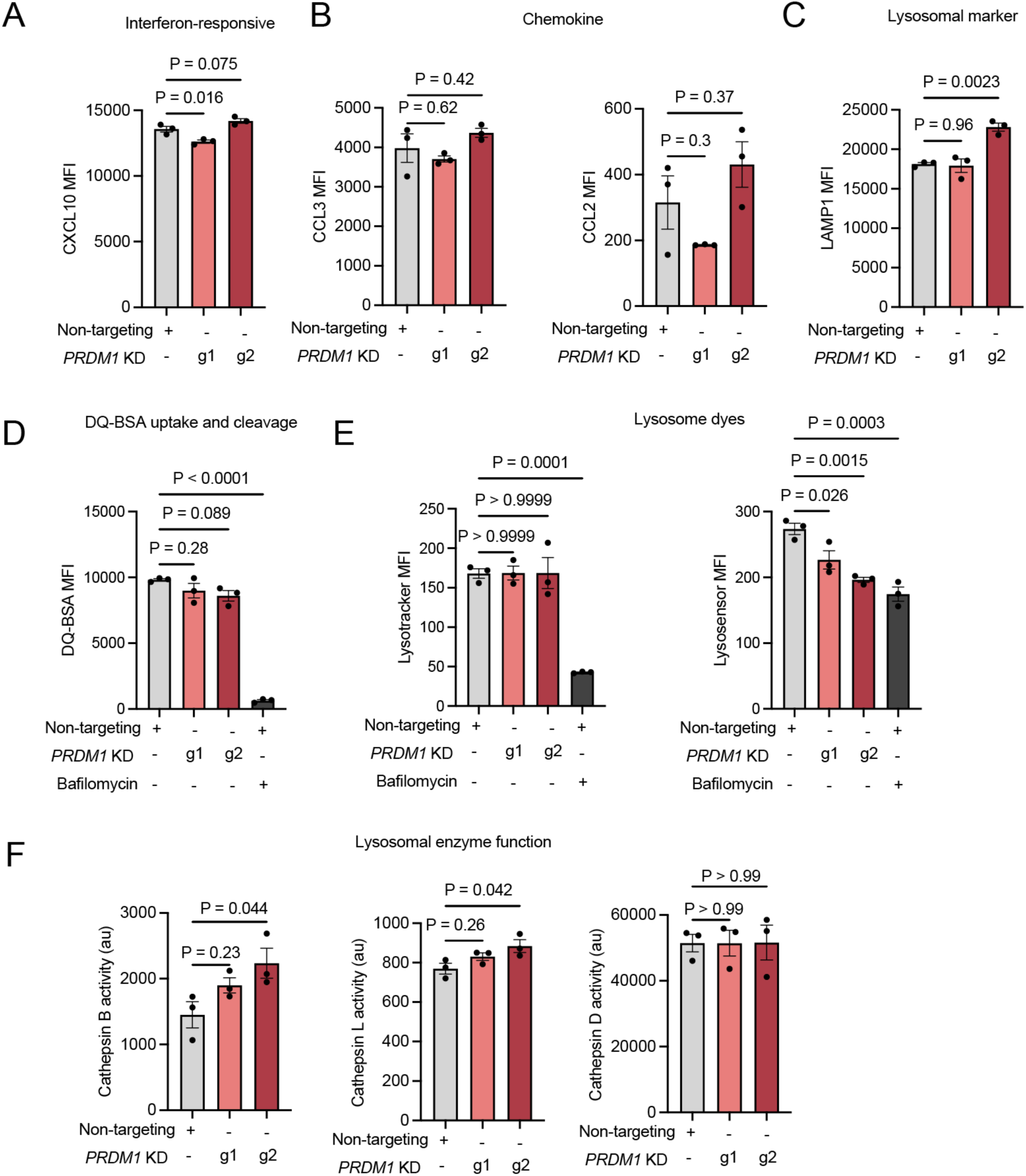
Quantification of additional protein state markers and lysosomal function in *PRDM1* KD. (A-C) Median fluorescence intensity (MFI) of state marker proteins by flow cytometry. *PRDM1* KD microglia (salmon, red) were compared to non-targeting controls (grey). Points represent one well, n ≥ 10,000 cells analyzed per well. One way ANOVA. (D) Median fluorescence intensity (MFI) of DQ-BSA by flow cytometry. DQ-BSA was added for two hours before analysis to allow for uptake. Bafilomycin A was added four hours before analysis to control wells. Points represent one well, n ≥ 10,000 cells analyzed per well. One way ANOVA. (E) Median fluorescence intensity (MFI) of lysotracker and lysosensor in *PRDM1* KD microglia (salmon, red) and non-targeting controls (grey). Bafilomycin A was added four hours before analysis to control wells. Points represent one well, n ≥ 10,000 cells analyzed per well. One way ANOVA. (F) Median fluorescence intensity (MFI) of LAMP1 by flow cytometry. *PRDM1* KD microglia (salmon, red) were compared to non-targeting controls (grey). Points represent one well, n ≥ 10,000 cells analyzed per well. One way ANOVA.

**Figure S11.**
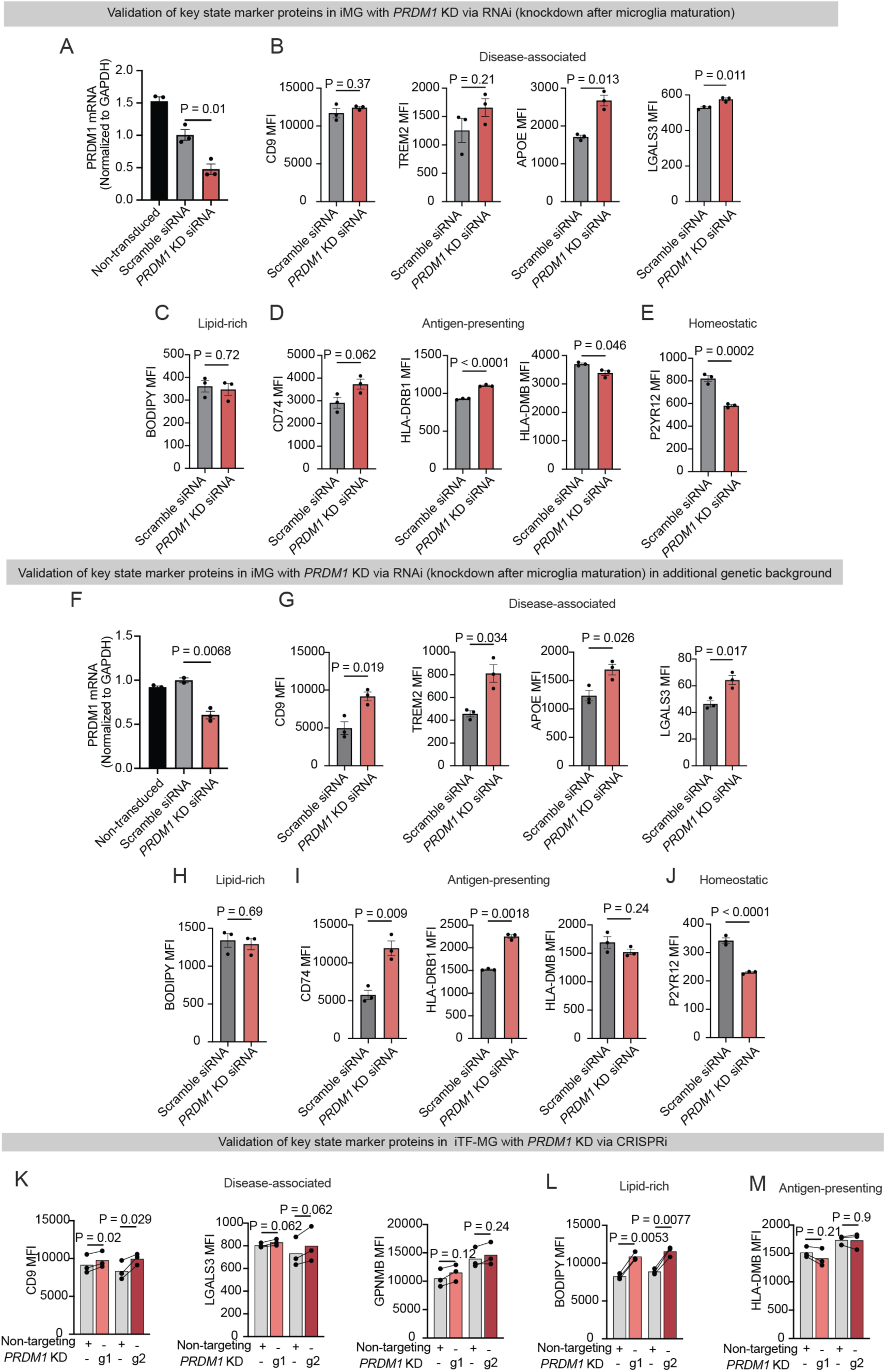
Validation of PRDM1 phenotypes using alternate methods, timing, and patient backgrounds. (A) Reverse-transcription quantitative PCR analysis of *PRDM1* expression after siRNA knockdown in WTC11 background. Cells not transduced with siRNA (black), transduced with scramble siRNA (grey) and *PRDM1* KD with siRNA (red). P value from one-way ANOVA. (B-E) Median fluorescence intensity (MFI) of state marker proteins by flow cytometry. *PRDM1* KD with siRNA (red) and scramble siRNA (grey). Points represent one well, n ≥ 10,000 cells analyzed per well. P value T-test. (F) Reverse-transcription quantitative PCR analysis of *PRDM1* expression after siRNA knockdown in KOLF2.1J background. Cells not transduced with siRNA (black), transduced with scramble siRNA (grey) and *PRDM1* KD with siRNA (red). P value from one-way ANOVA. (G-J) Median fluorescence intensity (MFI) of state marker proteins by flow cytometry. *PRDM1* KD with siRNA (red) and scramble siRNA (grey). Points represent one well, n ≥ 10,000 cells analyzed per well. P value T-test. (K-M) *PRDM1* KD in iTF-MG differentiation protocol. Median fluorescence intensity (MFI) of state markers by flow cytometry. *PRDM1* KD iTF-MG microglia (red) are compared to in-well non-targeting controls (grey). Points represent one well, n ≥ 10,000 cells analyzed per well. P value paired T-test.

**Figure S12.**
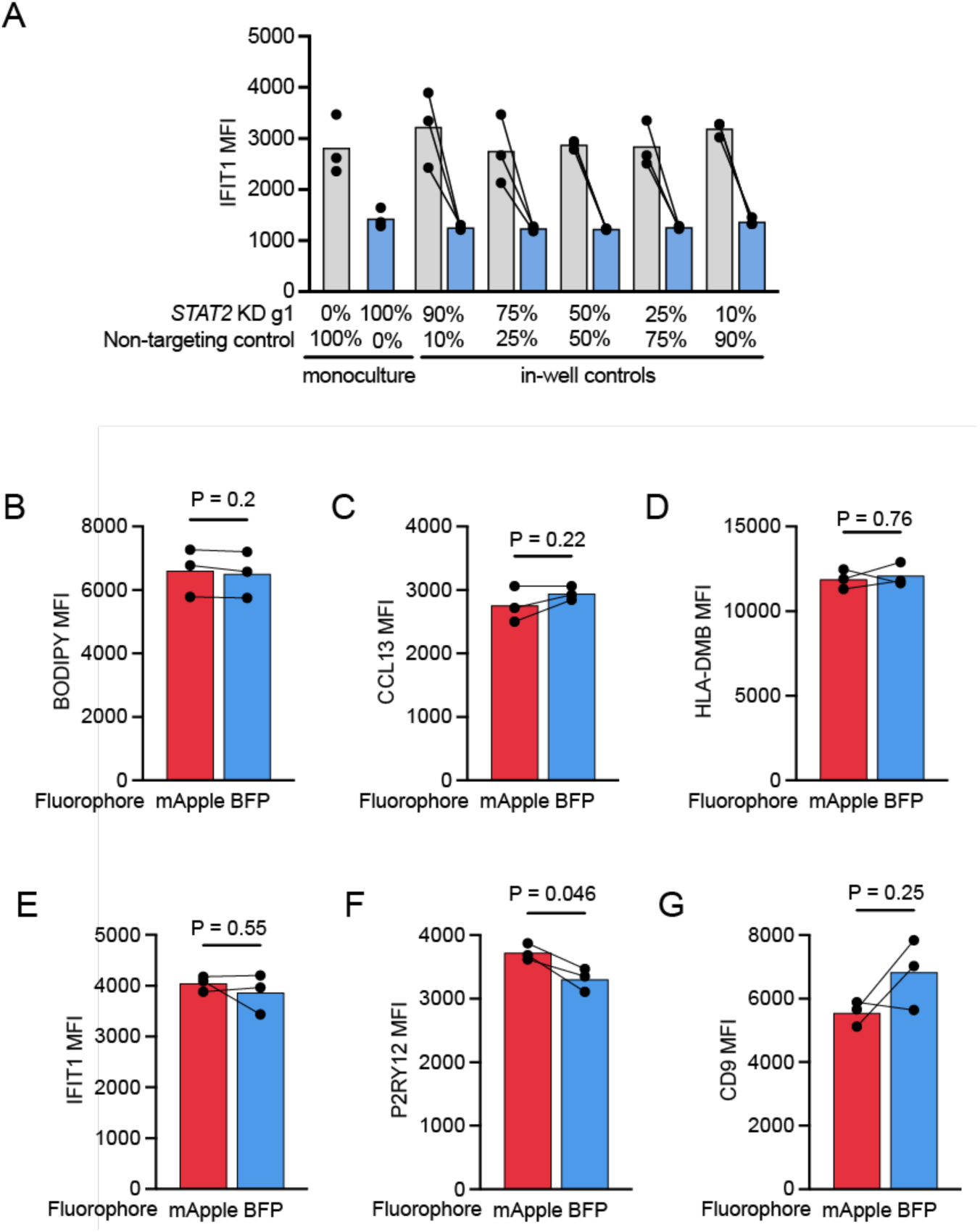
Neither co-culture of knockdown and control microglia nor expression of fluorophores to identify mixed populations change the state signature. (A) Median fluorescence intensity (MFI) of IFIT1 by flow cytometry. *STAT2* KD microglia (blue) were compared to in-well non-targeting controls (grey) distinguished by nuclear fluorescent proteins, or cultured in separate wells for the first two columns. At the beginning of differentiation (d0) cells with *STAT2* KD and non-targeting control guides were co-cultured at the stated percentage. Each point represents an independent well, n ≥ 10,000 cells analyzed per well. (B-G) Median fluorescence intensity (MFI) of state markers by flow cytometry. Non-targeting control cells expressing mApple (red) were compared to in-well non-targeting control cells expressing BFP (blue) distinguished by nuclear fluorescent proteins. Each point represents an independent well, n ≥ 10,000 cells analyzed per well. P values from paired Student’s T-test.

**Figure S13.**
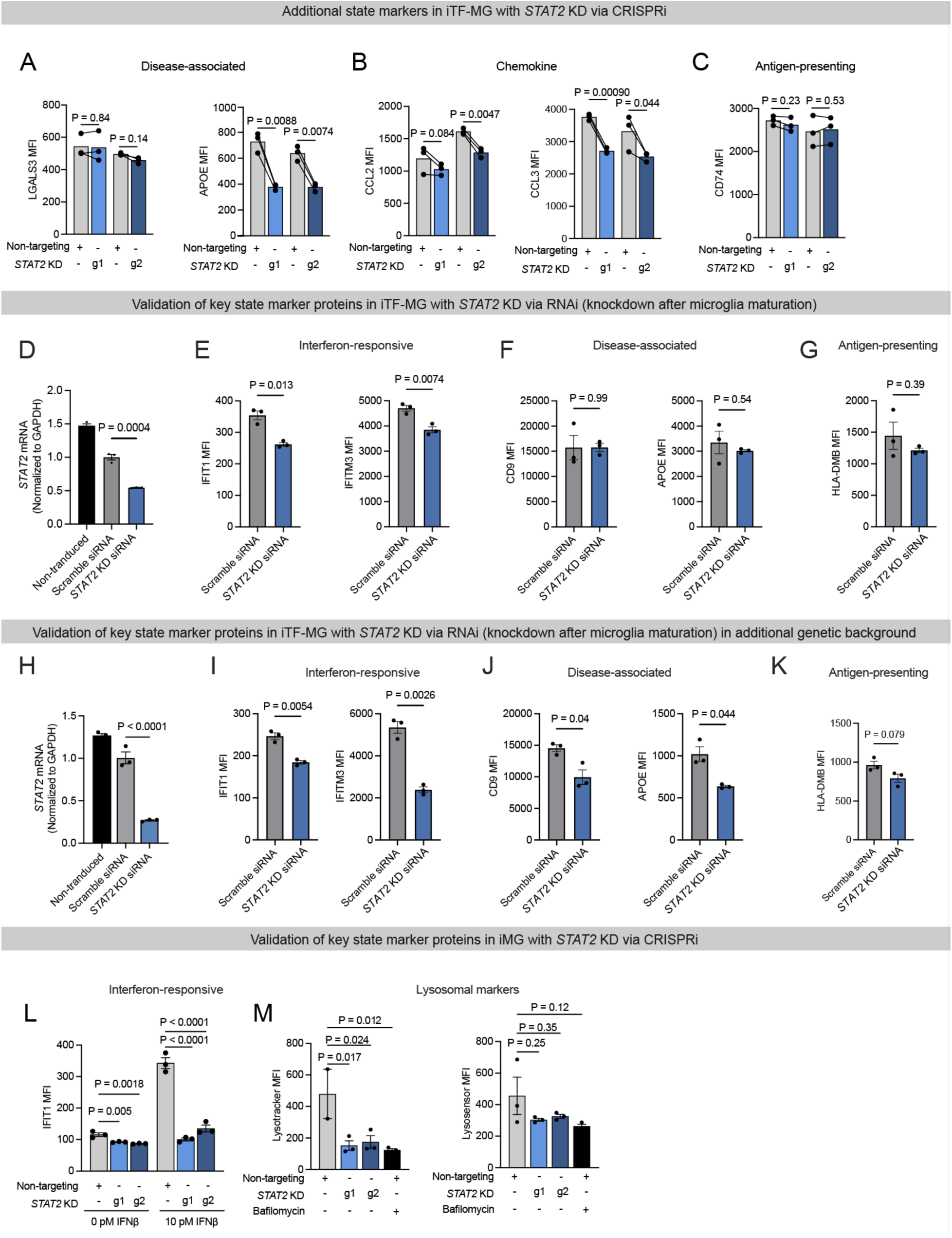
Validation of additional protein state markers, knockdown method, knockdown timeline, patient background, and differentiation protocol for *STAT2*. (A-C) Median fluorescence intensity (MFI) of state marker proteins by flow cytometry. *STAT2* KD microglia (blue) were compared to in-well non-targeting controls (grey) distinguished by nuclear fluorescent proteins. Points represent one well, n ≥ 10,000 cells analyzed per well. P value paired T-test. (D) Reverse-transcription quantitative PCR analysis of *STAT2* expression after siRNA knockdown in WTC11 background. Cells not transduced with siRNA (black), transduced with scramble siRNA (grey) and *STAT2* KD with siRNA (blue). P value from one-way ANOVA. (E-G) Median fluorescence intensity (MFI) of state marker proteins by flow cytometry. *STAT2* KD microglia (blue) and scramble siRNA (grey). Points represent one well, n ≥ 10,000 cells analyzed per well. P value paired T-test. (H) Reverse-transcription quantitative PCR analysis of *STAT2* expression after siRNA knockdown in KOLF2.1J background. Cells not transduced with siRNA (black), transduced with scramble siRNA (grey) and *STAT2* KD with siRNA (blue). P value from one-way ANOVA. (I-K) Median fluorescence intensity (MFI) of state marker proteins by flow cytometry. *STAT2* KD microglia (blue) and scramble siRNA (grey). Points represent one well, n ≥ 10,000 cells analyzed per well. P value paired T-test. (L,M) *STAT2* KD in iMG differentiation protocol. Median fluorescence intensity (MFI) of (L) IFIT1 and (M) lysosomal markers by flow cytometry. *STAT2* KD iMG microglia (blue) and non-targeting controls (grey). Points represent one well, n ≥ 10,000 cells analyzed per well. P value one-way ANOVA.

**Figure S14.**
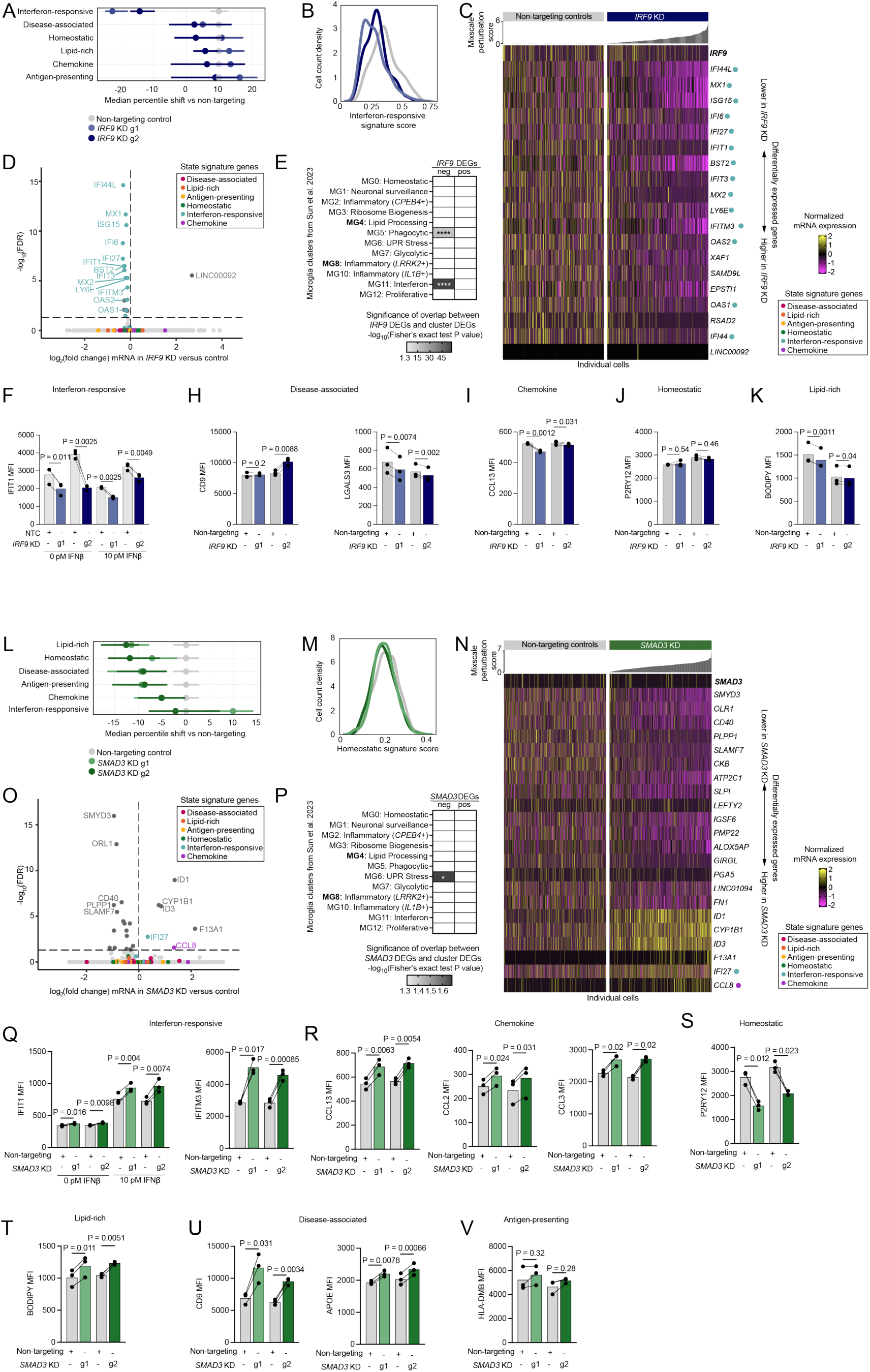
Loss of IRF9 and SMAD3 in iTF-MG. (A) Median percentile shift of state signature scores in *IRF9* KD (blue, navy) versus non-targeting controls (grey). Separate sgRNAs are represented as g1 and g2. Error bars represent 95% confidence interval. (B) Density plot of interferon-responsive signature score for *IRF9* KD (blue, navy) versus non-targeting controls (grey). (C) Heatmap of top differentially expressed genes (DEGs) ranked by FDR (all DEGs) shown. Columns represent individual cells ordered by Mixscale perturbation score (see methods). Scores are graphed above the corresponding column (non-targeting control perturbation scores are all zero by definition). Color scale represents the log-normalized mRNA expression. DEGs that appear in our state signature scores are denoted with a colored dot (pink: disease-associated, orange: lipid-rich, yellow: antigen-presenting, green: homeostatic, teal: interferon-responsive, purple: chemokine). (D) Volcano plot of differentially expressed genes in *IRF9* KD versus non-targeting control. FDR ≤ 0.05. DEGs that appear in our state signature scores colored as above. (E) Heatmap of gene signature overlap analysis of DEGs from *IRF9* KD with human microglia clusters from Sun et al. 2023^54^. Bolded states are increased in AD patients. Color denotes significance of overlap by Fisher’s exact test P value. Non-significant overlap is white. *P < 0.05, **P < 0.01, ***P < 0.001 ****P < 0.0001. (F-K) Median fluorescence intensity (MFI) of state marker proteins by flow cytometry. *IRF9* KD microglia (blue, navy) were compared to in-well non-targeting controls (grey) distinguished by nuclear fluorescent proteins. Each point represents an independent well, n ≥ 10,000 cells analyzed per well. P values from paired Student’s T-test. (L) Median percentile shift of state signature scores in *SMAD3* KD (green) versus non-targeting controls (grey). Separate sgRNAs are represented as g1 and g2. Error bars represent 95% confidence interval. (M) Density plot of homeostatic signature score for *SMAD3* KD (green) versus non-targeting controls (grey). (N) Heatmap of top differentially expressed genes (DEGs) ranked by FDR (all DEGs) shown. Columns represent individual cells ordered by Mixscale perturbation score (see methods for details). Scores are graphed above the corresponding column (non-targeting control perturbation scores are all zero by definition). Color scale represents the log-normalized mRNA expression. DEGs that appear in our state signature scores are denoted with a colored dot (pink: disease-associated, orange: lipid-rich, yellow: antigen-presenting, green: homeostatic, teal: interferon-responsive, purple: chemokine). (O) Volcano plot of differentially expressed genes in *SMAD3* KD versus non-targeting control. FDR ≤ 0.05. DEGs that appear in our state signature scores colored as above. (P) Heatmap of gene signature overlap analysis of DEGs from *SMAD3* KD with human microglia clusters from Sun et al. 2023^54^. Bolded states are increased in AD patients. Color denotes significance of overlap by Fisher’s exact test P value. Non-significant overlap is white. *P < 0.05, **P < 0.01, ***P < 0.001 ****P < 0.0001. (Q-V) Median fluorescence intensity (MFI) of state marker proteins by flow cytometry. *SMAD3* KD microglia (blue, navy) were compared to in-well non-targeting controls (grey) distinguished by nuclear fluorescent proteins. Each point represents an independent well, n ≥ 10,000 cells analyzed per well. P values from paired Student’s T-test.

**Figure S15.**
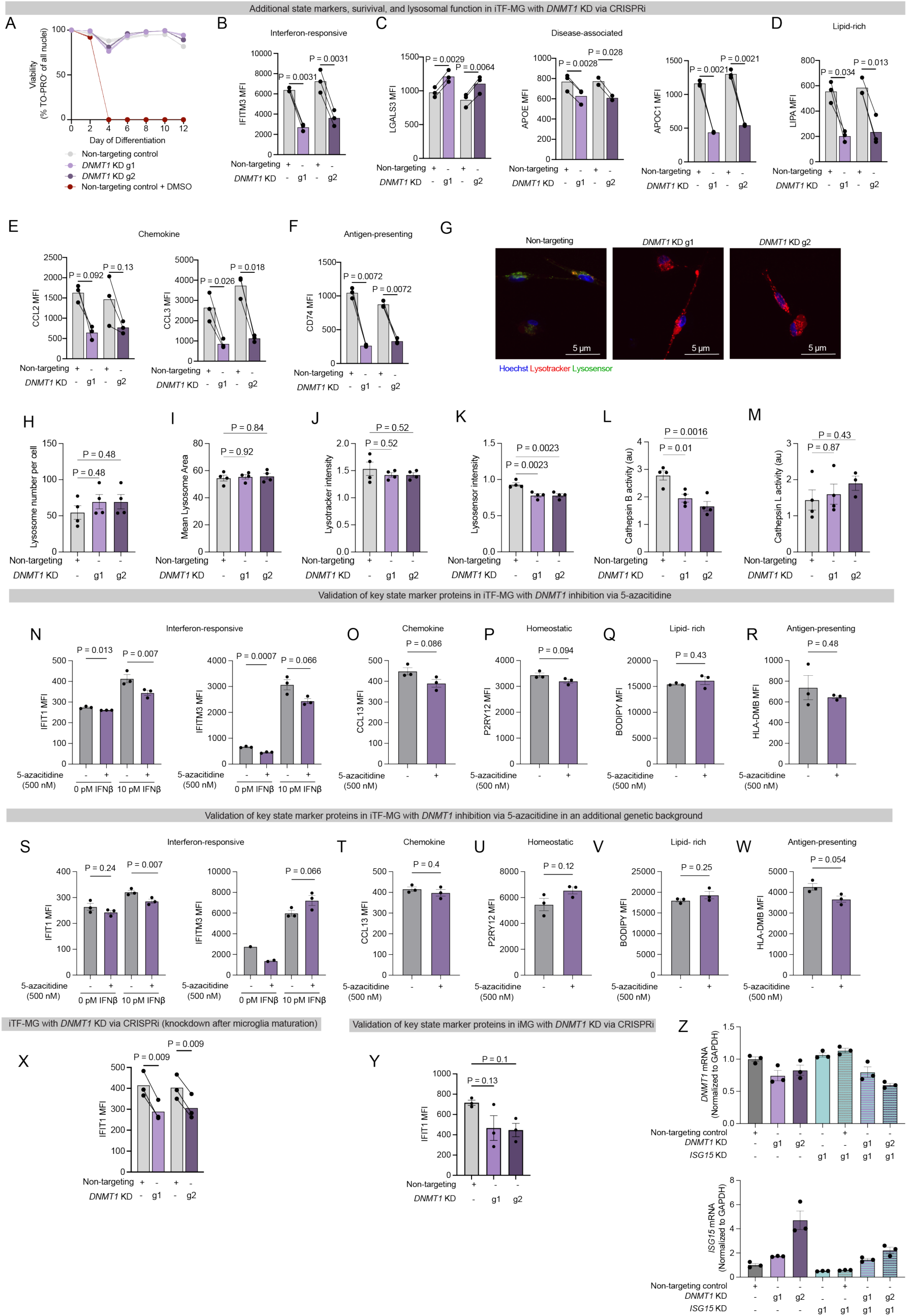
Validation of additional protein state markers, knockdown method, knockdown timeline, patient background, and differentiation protocol for *DNMT1*. (A) Knockdown of *DNMT1* does not alter viability. Viability of non-targeting control (grey) and *DNMT1* KD (violet) cells was measured every day of the differentiation into microglia by TO-PRO staining. n=3 wells, 4 images per well. 1% DMSO was added during normal media changes as a positive control for cell death. Data is represented as mean +/- standard error of the mean shown shaded around the line. (B-F) Median fluorescence intensity (MFI) of state marker proteins by flow cytometry. *DNMT1* KD microglia (violet) were compared to in-well non-targeting controls (grey) distinguished by nuclear fluorescent proteins. Points represent one well, n ≥ 10,000 cells analyzed per well. P value paired T-test. (G) Representative images of lysotracker (red), lysosensor (green), and Hoechst (blue) fluorescence in non-targeting control microglia or *DNMT1* KD microglia. Scale bar = 5 µm. Quantification of (H) lysosomal count per cell, (I) mean lysosomal area per lysosome, (J) lysotracker intensity within lysosomes normalized to Hoechst intensity, and (K) lysosensor intensity within lysosomes normalized to Hoechst intensity. N= 4 images per well in 4 separate wells. P value from One-way ANOVA. (L-M) Lysosomal protease activity in *DNMT1* KD microglia versus non-targeting control microglia measured by fluorescence of MagicRed signal for (L) Cathepsin B or (M) Cathepsin L. n=4 wells, 15,000 cells per well. P value from One-way ANOVA. (N-R) WTC11 background iTF-MG were treated with 500 nM 5-azacitidine from d2-12 to inhibit DNMT1. Median fluorescence intensity (MFI) of state marker proteins by flow cytometry. Microglia with DNMT1 inhibition shown in violet. Vehicle treated control microglia shown in grey. Points represent one well, n ≥ 10,000 cells analyzed per well. P value T-test. (S-W) Kolf2.1J background iTF-MG were treated with 500 nM 5-azacitidine from d2-12 to inhibit DNMT1. Median fluorescence intensity (MFI) of state marker proteins by flow cytometry. Microglia with DNMT1 inhibition shown in violet. Vehicle treated control microglia shown in grey. Points represent one well, n ≥ 10,000 cells analyzed per well. P value T-test. (X) IFIT1 expression in WTC11 background iTG-MG with TMP added from d8-20 to knock down *DNMT1* in mature microglia. *DNMT1* KD microglia (violet) were compared to in-well non-targeting controls (grey) distinguished by nuclear fluorescent proteins. Points represent one well, n ≥ 10,000 cells analyzed per well. P value paired T-test. (Y) *DNMT1* KD in iMG differentiation protocol. Median fluorescence intensity (MFI) of IFIT1. *DNMT1* KD iMG microglia (violet) and non-targeting controls (grey). Points represent one well, n ≥ 10,000 cells analyzed per well. P value T-test. (Z) Reverse-transcription quantitative PCR analysis of *DNMT1* expression (top) or *ISG15* expression (bottom). Non-targeting controls (grey), *DNMT1* KD (purple), *ISG15* KD (teal).

## Legends for Supplemental Tables

Table S1. **Screen results for FACS-based screens.** Gene level results for all six screens. Each tab represents a separate CRISPRi screen: CD9, P2RY12, HLADMB, BODIPY, IFIT1, CCL13. Columns include fold change between high and low sorted cells (fc), log2fc, P value, FDR, and phenotype score which is the product of the log2fc and FDR.

Table S2. **Data related to CROP-seq and CITE-seq analyses**

<u>Tab 1: Primers used in this manuscript</u>

<u>Tab 2: Protospacer sequences and sgRNA identities for CROP-seq, CITE-seq, and downstream analyses.</u> Columns: Gene name, guide number, protospacer name, protospacer sequence

<u>Tab 3: State signature gene definitions</u>. Gene lists used for each state signature score. Each column represents a separate state.

<u>Tab 4: Median percentile shift for each signature score at the level of Genes and sgRNA in iTF-MG and iMG.</u> Columns represent the signature, perturbed gene or guide RNA, median percentile, percentile shift from non-targeting control cells, and q value, differentiation protocol used, and if the calculation is at the gene or guide level.

<u>Tab 5: iTF-MG Pearson’s correlation of CITE-seq proteins with signature scores.</u> First column represents all 166 proteins and their correlation with each mRNA gene signature UCell scores. Values represent Pearson’s correlation.

<u>Tab 6: iMG Pearson’s correlation of CITE-seq proteins with signature scores.</u> First column represents all 166 proteins and their correlation with each mRNA gene signature UCell scores. Values represent Pearson’s correlation.

<u>Tab 7: iTF-MG Weighted differential expression for CROP-seq.</u> Columns represent perturbed gene targeted by CRISPRi, gene transcript name, log2fc, beta weight, P value, and the adjusted P value. Differentially expressed genes for all perturbations that were determined to be significant by Mixscale are included.

<u>Tab 8: iTF Weighted differential expression for CROP-seq.</u> Columns represent perturbed gene targeted by CRISPRi, gene transcript name, log2fc, beta weight, P value, and the adjusted P value. Differentially expressed genes for all perturbations that were determined to be significant by Mixscale are included.

<u>Tab 9: Gene set enrichment analysis</u>. GSEA performed for significant DEGs (top 100 genes) of featured perturbations. Columns represent the differentiation protocol, gene perturbed, direction of DEGs, database used, term name, significance, set of genes present in the term, and adjusted significance.

<u>Tab 10: iTF-MG Weighted differential abundance for CITE-seq.</u> Columns represent perturbed gene targeted by CRISPRi, protein name, log2fc, beta weight, P value, and the adjusted P value.

<u>Tab 11: iTF Weighted differential abundance for CITE-seq.</u> Columns represent perturbed gene targeted by CRISPRi, protein name, log2fc, beta weight, P value, and the adjusted P value.

<u>Tab 12: Overlap analysis with external data.</u> For STAT2, DNMT1, IRF9, SMAD3, ZNF532, and PRDM1, positive and negative DEGs were compared to existing datasets. Columns represent the differentiation model, perturbed gene, direction of the DEG list analyzed (positive DEGs or negative DEGs), the cluster and paper of overlap, the number of genes overlapped between the DEGs and the cluster, the percentage of genes overlapped, the name of genes overlapped, and the fisher’s exact test P value.

<u>Tab 13: Median percentile shift for each scHPF factor at the level of Genes and sgRNA in iTF-MG and iMG.</u> Columns represent the signature, perturbed gene or guide RNA, differentiation protocol used, median percentile, percentile shift from non-targeting control cells, and q value.

<u>Tab 14: Gene Regulatory Network TF targets overlap with differentially expressed genes</u> Columns represent the method of target prediction (SCENIC or SCENIC without pruning for motifs for ZNF532 since no motif exists (grn_znf532)), the transcription factor source (tf), the number of predicted targets, number of significant DEGs, overlap of these values, percentage of predicted that are captured in the DEGs, percentage of the DEGs captured by the predicted, and overlap analyses.

<u>Tab 15: Differential expression of non-targeting controls from iMG and iTF-MG</u> for microglia and macrophage related genes. Expression is shown as normalized values.

Table S3. **Analysis of secreted proteins by nELISA.** First tab shows differential expression of 978 secreted proteins from conditioned media in iTF-MG vs iMG. Columns represent the nELISA protein ID, protein names, test and control cell lines, number of wells used per line, central value, fold change, log2fold change, P value, and adjusted P value. Second tab shows differential expression of 978 secreted proteins in iTF-MG cells with non-targeting guides, *STAT2* KD, or *DNMT1* KD with or without IFNβ treatment for 24 hrs. Columns represent the CRISPRi guide and treatment used, fold change, log2 fold change, P value, adjusted q value, number of wells used per line, and protein detected.

Table S4. **Differential methylation at gene promoters from EM-seq.** Columns represent ensembl gene identification numbers, P value, FDR, and the mean difference where negative values are hypo-methylated in *DNMT1* KD cells versus non-targeting control guides.

## Methods

### iPSC maintenance

We cultured human induced pluripotent stem cells in either StemFlex (Gibco, A33493-01, before transcription factor based differentiation protocol^57^) or mTeSR1 (Stemcell Technologies, 85850, before cytokine-driven differentiation protocol^58,116^). iPSCs were cultured on Matrigel (Corning, 356231) coated cultureware. Cells were passaged once per week via ReLeSR (Stemcell Technologies, 100-0483). Human iPSC studies at the University of California, San Francisco were approved by the Human Gamete, Embryo and Stem Cell Research Committee.

### Transcription factor-directed differentiation of iPSC-Microglia (iTF-iMG)

iTF-MG were differentiated as previously described^57^ with a modified media composition optimized for increased expression of P2RY12, TREM2, and CD33 as follows. Induced pluripotent stem cells containing six doxycycline-inducible transcription factors: CEPBA, CEBPB, IRF5, IRF8, MAFB, PU.1, were passaged as a single-cell suspension using accutase (Thermo Fisher Scientific, A11105-01) for 7 minutes at 37 °C. Cells were plated to dual-coated Matrigel (Corning, 356231) and PDL plates (Corning, 356470) and cultured for 48 hours in Essential 8 (Gibco, A1517001) stem cell maintenance media supplemented with 2 µg/mL doxycycline (Clontech, 631311) and 10 nM Y-27632 (Tocris 1254). Two days post passage, cells were moved to microglia differentiation media: BrainPhys (StemCell Technologies, 05790), 0.5x N2 Supplement (Gibco, 17502048), 0.5x B27 (Gibco, 17504044), 10 ng/mL NT-3 (Peprotech, 450-03), 10 ng/mL BDNF (Peprotech, 450-02), 1 µg/mL mouse laminin (Thermo Fisher Scientific, 23017-015) supplemented with 2 ug/mL doxycycline, 100 ng/mL IL-34 (Peprotech 200-34), and 10 ng/mL GM-CSF (Peprotech 300-03). On day 4, day 8, and day 12, media was replaced with microglia differentiation media supplemented with 2 µg/mL doxycycline, 100 ng/mL IL-34, 10 ng/mL GM-CSF, 50 ng/mL M-CSF (Peprotech 300-25), and 50 ng/mL TGF-β1 (Peprotech100-21C). For experiments with CRISPRi, cells were supplemented with 50 nM trimethoprim (TMP) (MP Biomedical, 195527) every other day to maintain strong knockdown.

### Cytokine-directed differentiation of iPSC-Microglia (iMG)

iMG were differentiated as previously described^58,116^. iPSCs were passaged in small colonies ∼50-100 cells onto Matrigel-coated plates using ReLeSR (Stemcell Technologies, 100-0483) at a density suitable to achieve ∼40 colonies per well the next day. Wells with the correct size and density of iPSCs were differentiated to hematopoietic progenitor cells (HPCs) using StemDiff Hematopoietic kit (Stemcell Technologies, 05310). HPCs were collected on days 10, 12, and 14 and frozen in BamBanker (Bulldog Bio, BB05). To generate iMG, HPCs were thawed at 7-10K cells/cm^2^ into microglia differentiation media: DMEM/F12, 2X insulin-transferrin-selenite, 2X B27, 0.5X N2, 1X glutamax, 1X non-essential amino acids, 400 μM monothioglycerol, 5 μg/mL insulin. Immediately before use, microglial medium was supplemented with 100 ng/mL IL-34, 50 ng/mL TGF-β1, and 25 ng/mL M-CSF (Peprotech) taken from single-use frozen aliquots. Media with fresh cytokines was supplemented 1 mL/well of a 6-well plate every 48 hours for 28 days.

### Lentivirus generation for individual sgRNAs

Lentivirus was generated as described^117^. HEK-293T cells were plated to achieve 80-95% confluence after 24 hours. For 2 mL of media, 1 µg transfer plasmid and 1 µg third generation packaging mix were added to 100 µL Optimem (Gibco, 31985088) and 12 µL TransIT-Lenti Transfection Reagent (Mirusbio, MIR 6604). Transfection mix was incubated for 10 min at room temperature before addition to cells. After 48 hours, conditioned media was collected and filtered through a 0.45 µm PVDF filter. Lentivirus Precipitation solution (Alstem, VC125) was used per manufacturer’s protocols to isolate lentivirus.

iPSCs were infected with lentivirus immediately after single-cell passaging with accutase and treated for 48 hours. After transduction, cells were again single-cell passaged and selected with 1 µg/mL puromycin for two passages or until populations with >95% BFP-positive cells were achieved.

### Pooled CRISPRi screening

For screens of transcription factors and transcriptional regulators, we used a previously develope^94^ sgRNA library targeting 1619 transcription factors and transcriptional regulators, with five independent single-guide RNAs (sgRNAs) per gene and 250 non-targeting sgRNAs. sgRNAs were chosen using the design of the Horlbeck et al. CRISPRi_v2 library^118^ and cloned into pLG15 vector^117^. To create lentivirus, HEK-293T cells were transduced as described above at an MOI of 0.3 to minimize multiple transduction of the same cell with multiple sgRNAs. To ensure equal distributions of gRNAs, transduction was performed at a coverage of 3000x such that every sgRNA would be represented in 3,000 iPSCs on average. For all cell culture, freezing, and differentiation, transduced iPSCs were maintained with at least 1000x average coverage of sgRNAs.

For each screen of microglial activation states, 20M iPSCs were differentiated using the iTF-MG protocol described above, with addition of TMP to induce CRISPRi activity starting at day 0. On day 12 of differentiation, iTF-MG were lifted with TrypLE for 10 min at 37 °C and pelleted at 300 x *g* for 5 min. To stain dead cells for removal, iTF-MG were resuspended in Zombie Red Fixable Viability kit (Biolegend, 423110, 1:200) for 10 min at 4 °C.

For intracellular markers (IFIT1, CCL13), iTF-MG were fixed with zinc fixation buffer (0.1M Tris-HCl, pH 6.5, 0.5% ZnCl_2_, 0.5% Zn Acetate, 0.05% CaCl_2_) overnight at 4 °C as in Samelson et al^119^. Following fixation, cells were stained with anti-IFIT1 (1:100, Cell Signaling, 20329S) or anti-CCL13 (1:50, R&D Systems, IC327G) in TBS-based FACS buffer (TBS + 3% BSA + 0.5 uM EDTA) for 30 min at 4 °C. Cells were washed in TBS-based FACS buffer and strained through a 20 µm filter.

For extracellular markers, antibody was added to live cells in PSB-based FACS buffer (DPBS + 3% BSA + 0.5 µM EDTA) for 30 min at 4 °C. The following antibodies or dye was used: anti-CD9 (1:200, Biolegend, 312104), anti-P2RY12 (1:50, Biolegend, 392108), anti-HLA-DMB (1:200, Proteintech, 82922-1-RR) BODIPY (1:1000, ThermoFisher, D3922). Following staining, cells were washed in PBS-based FACS buffer and strained through a 20 µm filter.

iTF-MG were sorted using a BD FACSAria Fusion flow cytometer. Cells were gated on live, single cells, and sorted into top 30% and bottom 30% of marker expression. After sorting, genomic DNA was isolated with NucleoSpin Blood L kit (Macherey-Nagel, 740954.20). sgRNA cassettes were isolated and amplified as previously described^117^. Screening libraries were sequenced on Illumina MiSeq.

### Single-cell CRISPRi screening (CROP-seq and CITE-seq)

Library Generation: The CROP-seq library was generated targeting 31 genes that had interesting phenotypes in the original six FACS-based screens. The two sgRNAs with the strongest phenotypes in the FACS-based screens were selected for each gene for inclusion in the CROP-seq library. In total, 31 genes with 2 sgRNAs each were pooled alongside 5 non-targeting sgRNAs that did not influence phenotypes in any of the original FACS-based screens, resulting in a final library size of 65 elements. The crop-seq library was prepared as previously described^57^ from arrayed oligonucleotides ordered from Integrated DNA Technologies. Top and bottom oligonucleotides for each sgRNA were annealed and then pooled. This pool was cloned into pMK1334^117^ after restriction digest with BstXI and BlpI. The pooled plasmid library was cloned into Stellar Competent cells such that each plasmid was estimated to have been transformed into >500 bacteria. We performed sgRNA enrichment PCR and sequenced this plasmid pool using MiSeq to confirm normal distribution of all sgRNAs. Lentivirus was prepared and added to iPSCs as described above.

Cell preparation: iMG and iTF-MG containing the CROP-seq library were differentiated at >1000x coverage of the library. HPC differentiations were carried out at ∼5000x coverage to reduce the impact of bottlenecking at the HPC collection stage. iTF-MG and iMG differentiations were timed to reach day 12 and day 28 respectively on the same day. iTF-MG were lifted with TrypLE for 10 min at 37 °C. iMG are grown non-adherently and lifted in the media. Cells were treated with TruStain FcX Fc block (1:200, Biolegend, 422301) for 10 min prior to addition of CITE-seq antibody pools as described in Haage et al. 2025^83^. The Total-Seq A Universal (Biolegend, 399907) and custom^83^ antibody pools were centrifuged for 30 sec at 10,000 x *g* before reconstitution in 27.5 µL Cell Staining Buffer (Biolegend, 420201) total for both antibody panels and incubated at room temperature for 5 min. Before adding to iMG, the antibody pools were centrifuged for 14,000 x *g* for 10 min at 4 °C. 500,000 iMG and iTF-MG were incubated with the antibody panel for 30 min at 4 °C. After staining, cells were washed with 1 mL Cell Stating Buffer four times with centrifugation of 300 x *g* for 10 min. Cells were filtered with 40 µm filter, counted, and diluted to 1.5M cells per mL. Cells were loaded into the 10x Chromium Controller (10X Genomics v4) at 40,000 cells per lane for one lane per sample according to the manufacturer’s instructions. Single-cell sequencing libraries for CROP-seq were prepared as previously described^57^. CITE-seq libraries were prepared using ADT primers per manufacturer’s instructions. Libraries were sequenced on NovaSeq X 10B.

For iTF-MG, CROP-seq was performed with two biological replicates that were differentiated, stained, and sequenced separately and merged as described in the computational methods.

### Generating sgRNA knockdown lines

sgRNAs with strong phenotypes in the original screens were selected for follow up studies. sgRNAs were cloned into pMK1334 as previously described^74^. Individual gRNAs were added to iPSCs and selected as described above in lentivirus production. For samples with in-well non-targeting control, the plasmid backbone blue fluorescent protein (BFP) was swapped for mApple as previously described^74^. sgRNA sequences are listed in Table S2.

### Small interfering RNA (siRNA) knockdown

As an orthogonal method of knockdown, we used siRNA. In iTF-MG, microglia were nucleofected at d8 of differentiation. In iMG, microglia were nucleofected at d23. In both protocols, cells were nucleofected with 45 pmol siRNA (ON-TARGETplus 2.0 SMARTpool, Horizon Discovery) per million cells using the Human Macrophage Nucleofection kit (Lonza, VPA-1008) Y-001 program. Cells were allowed to recover until maturity (d12-13 for iTF-MG, d28 for iMG).

### Reverse-transcriptase quantitative polymerase chain reaction

To confirm RNA knockdown for independent sgRNA knockdown lines, RNA was extracted from mature microglia using the Quick-RNA Microprep Kit (Zymo, R1050). cDNA was generated using SensiFAST cDNA Synthesis Kit (Meridian Bioscience, BIO-65053). For quantitative PCR, SensiFAST SYBR Lo-ROX 2X Master Mix (Bioline; BIO-94005) was used with custom qPCR primers from Integrated DNA Technologies. Amplification was quantified using the Applied Biosystems QuantStudio 6 Pro Real-Time PCR System using QuantStudio Real Time PCR software (v.1.3). Fold change in RNA expression was calculated using ΔΔCt. Data normalized to *GAPDH* to control for RNA concentration.

### Flow cytometry

iTF-MG were lifted with TrypLE (Gibco, 12604021) for 10 min at 37°C. iMG are cultured in a non-adherent format and do not require dissociation. For both cell types, iPSC-Microglia were pelleted at 300 x *g* for 5 min. iPSC-Microglia were resuspended in Zombie Live/Dead stain (1:200, Biolegend, multiple cat#) for 10 min at room temperature and washed in DPBS.

For extracellular markers, iPSC-Microglia erre stained for 30 min at 4 °C in FACS buffer (DPBS + 3% BSA + 0.5 µM EDTA). Extracellular antibodies: anti-CD9 (1:200, Biolegend, 312104), anti-P2RY12 (1:50, Biolegend, 392108), anti-HLA-DMB (1:200, Proteintech, 82922-1-RR), anti-LGALS3 (1:200, Biolegend, 125410), anti-GPNMB (1:200, Invitrogen, 740041MP488), anti-HLA-DRB1 (1:200, Biolegend, 327010), anti-CD11b (1:200, Biolegend, 301329,), anti-CD45 (1:200, Biolegend, 368515), anti-TMEM119 (1:200, Biolegend, 853318), anti-CD86 (1:200, Biolegend, 305406), anti-CD55 (1:50, Biolegend, 311306), anti-CD81 (1:200, Biolegend, 349504), anti-CD28 (1:50, Biolegend, 302906), anti-C5aR (1:50, Biolegend, 344306).

For intracellular markers, iPSC-Microglia were first fixed with with eBioscience Intracellular Fixation and Permeabilization Buffer Set (Invitrogen, 888-8824-00) for 20 min at room temperature and washed in permeabilization buffer. Antibodies were diluted in permeabilization buffer and added to cells for 30 min at 4 °C. Intracellular antibodies: anti-IFIT1 (1:100, Cell Signaling, 20329S), anti-CCL13 (1:50, R&D Systems, IC327G), anti-IFITM3 (1:200, Proteintech, CL488-11714), anti-APOE (1:200, Invitrogen, MA5-15852), anti-CCL2 (1:50, Biolegend, 505910), anti-CCL3 (1:200, Biolegend, 934603), anti-CD74 (1:50, Biolegend, 326808), anti-SPP1 (1:50, eBioscience, 50-9096-42), anti-TREM2 (1:100, R&D Systems, MAB18281), anti-CXCL10 (1:50, Biolegend, 519504), anti-LAMP1 (1:200, Biolegend, 328610), anti-APOC1 (1:200, Invitrogen, PA5-145261), anti-LIPA (1:200, Proteintech, 12956-1-AP), anti-PU.1 (1:200, Cell Signaling, 2266S).

For secreted markers (CCL13, CXCL10, SPI1), cells were pre-treated with GolgiPlug (1:1000, BD Biosciences, 555029) 6 hours prior to lifting.

Samples were analyzed on a BD FACSCelesta or BD LSR Fortessa X14 using BD FACSDiva software. Median fluorescence intensity was calculated using FlowJo analysis software after gating for live, single cells. Data were collected from 3 independent differentiations with 3 replicate wells per experiment (n ≥ 10,000 cells).

### nELISA secreted protein analysis

Soluble protein levels were measured using the nELISA assay (Nomic Bio) for high-throughput protein profiling, which quantified 1000 proteins per well. Samples were sent to Nomic Bio (Montreal, Canada) for analysis with the Omni 1000 panel, using previously described protocols^82^. Briefly, the Nomic platform pre-assembles antibody pairs on spectrally encoded microparticles^120^, resulting in spatial separation between non-cognate antibodies, preventing the rise of reagent-driven cross-reactivity, and enabling multiplexing of >1000 immunoassays in parallel using flow cytometry. Standard curves were included for each protein, enabling quantification of pg/mL concentrations.

### Timelapse imaging phagocytosis assay

iMG were collected on day 28 and plated at 50% confluence on Matrigel-coated 96-well plates and allowed to recover overnight. After addition of pHrodo tagged human synaptosomes (10 ng/mL) or pHrodo-tagged beta-amyloid (500 nM), images were collected every hour on an IncuCyte S3 Live-Cell Analysis System (Sartorius) with 4 images per well in 4 independent wells per condition. IncuCyte 2023A software was used to mask fluorescent substrate signal normalized to cell body area.

### Beta-amyloid fibrillization and labelling

Beta-amyloid (1-42) monomer (rPeptide, A-1170-02) was resuspended in 100 µM NaOH and incubated at 37 °C overnight. Fibrillarized beta-amyloid was tagged with pHrodo-Red, Succinimidyl Ester (ThermoFisher Scientific, 36600) in 0.1M sodium bicarbonate for 1 hour at room temperature, washed and resuspended at 100 µM in DPBS.

### Isolation and labelling of human synaptosomes

Synaptosomes were isolated using Method 2 from Hahn et al.^121^ Briefly, brain samples were dounce-homogenized in 320 mM sucrose, 0.1 mM CaCl_2_, and 1 mM MgCl_2,_ with protease and phosphatase inhibitors. Synaptosomes were isolated using a sucrose gradient centrifugation at 100,000 x *g* at 4 °C for 3 hours. Synaptosomes were collected at the interface, diluted in 0.1 mM CaCl_2_ and centrifuged for 15,000 x *g* at 4 °C for 20 min to remove larger membrane fragments. Synaptosomes were tagged with pHrodo-Red, Succinimidyl Ester (ThermoFisher Scientific, 36600) per manufacturer’s instructions.

### Flow cytometry analysis of lysosomal markers

To analyze lysosome markers and activity, control cells were pre-treated with Bafilomycin A (200 nM, Sigma, SML1661) for 4 hours. For uptake analysis, DQ Green BSA (10 µg/mL, Invitrogen, D12050) was added for 2 hours. For lysosome tracking, Lysotracker Deep Red (20 nM, Invitrogen, L12492) or Lysosensor Green (500 nM, Invitrogen, L7535) were added for 30 minutes. Samples were analyzed on BD FACSCelesta using BD FACSDiva software. Median fluorescence intensity was calculated using FlowJo analysis software after gating for live, single cells. Data were collected from 3 independent differentiations with 3 replicate wells per experiment (n ≥ 10,000 cells).

### Microscopy

Mature microglia were treated with Lysotracker Deep Red (20 nM, Invitrogen, L12492), Lysosensor Green (500 nM, Invitrogen, L7535), and Hoechst (ThermoFisher, R37165) for 30 minutes. Live cells were imaged on a spinning disk high-speed confocal microscope with super-resolution mode (Leica DMi8 with Yokogawa CSU-W1 SoRa) at 40x magnification plus 1.5x optical zoom. 4 wells were imaged per cell line and 4 z-stack images were taken per well. Images were analyzed blind on CellProfiler^122^.

### Cathepsin Activity assay

Activity of cathepsin B (Abcam, ab270772) and cathepsin L (Abcam, ab270774) was measured by fluorometric Magic Red cleavage assay per manufacturer’s instructions for three wells per cell line. Activity of cathepsin D was measured by fluorometric substrate cleavage kit (Abcam, ab65302) per manufacturer’s instructions for three wells per cell line. Fluorescence was quantified on an Agilent BioTek Synergy H1 plate reader.

### TO-PRO viability assay

To assess viability, cells were treated with TO-PRO-3 (Thermo Fisher, R37113) and Hoechst (Thermo Fisher, R37165) for 15 min before imaging on an IN Cell Analyzer 6000 (Cytiva). Images were analyzed using CellProfiler^122^ by masking all nuclei with the Hoechst, and performing a binary classification of TO-PRO-3 negative (live) cells and TO-PRO-3 positive (dead) cells.

### Enzymatic-methylation sequencing

Whole-genome enzymatic methyl sequencing (WGEMS) libraries were generated for DNMT1 knockdown or non-targeting control microglia. Genomic DNA was isolated using NucleoSpin Blood L kit (Macherey-Nagel, 740954.20). 200 ng of gDNA was diluted to 24 μL with TE buffer. Control DNA Unmethylated Lambda (NEB, E7122AVIAL) and Control DNA CpG-methylated pUC19 (NEB, E7123AVIAL) were diluted 1:50 with TE buffer and 1 µL of each was added to the genomic DNA. All samples were sheared to 240-280 bp using NEBNext UltraShear® (NEB, M7634L) following manufacturer recommendations. End prep and adaptor ligation was performed using 44 μL of fragmented DNA following the NEBNext UltraShear® manual (version 2.0, section 5). NEBNext® Enzymatic Methyl-seq Kit (NEB, E7120S/L) was used, with formamide denaturation for enzymatic methyl conversion. 5 ng of deaminated DNA was PCR-amplified for 6 cycles, followed by PCR-clean up and eluted in TE buffer according to protocol. Amplified libraries were quantified on an Agilent 2100 TapeStation system (Agilent). Samples were pooled at equimolar concentration and sequenced with paired-end 150 bp reads on an Illumina NovaSeq X 10B. We obtained between 183.6M to 299M paired-end reads across all libraries.

### EM-sequencing analysis

Reads were trimmed using cutadapt (version 4.4)^123^, filtering empty resulting reads (-m 1) and specifying both forward (-a AGATCGGAAGAGCACACGTCTGAACTCCAGTCA) and reverse (-A AGATCGGAAGAGCGTCGTGTAGGGAAAGAGTGT) adapters. The resulting reads were aligned using Bismark (version 0.24.2)^124^ to the hg38 reference genome with methylated and unmethylated control DNA sequences provided by NEBNext (https://neb-em-seq-sra.s3.amazonaws.com/grch38_core%2Bbs_controls.fa). The Bismark tool deduplicate_bismark was used to deduplicate aligned reads. Next, bismark_methylation_extractor and bismark2bedGraph were then used to create a methylation report for CpG sites for the top and bottom strands. Additionally, coverage2cytosine was used to generate a cytosine report aggregating top and bottom strand reads. The top and bottom strand bedgraph files were combined (on the basis of CpG symmetry), and loci that had less than 3 reads after combining strands were filtered out. To ensure that samples within each condition were statistically independent, technical replicates for each sgRNA were combined by summing the total number of reads and number of methylated reads at each locus, resulting in one sample per sgRNA, for a total of two samples per condition.

Differential methylation analysis was performed using bsseq (version 1.40.0)^125^ and DSS (version 2.52.0)^126^ in R (version 4.3.3) by calling DMLtest with smoothing=TRUE. The analysis was performed between the non-targeting and the *DNMT1* KD cells each with two independent sgRNAs, for a total of 4 samples per differential analysis.

For aggregation, promoter regions were defined as +/- 500 bp from the TSS sites, and methyl sites were aggregated by mean difference of *DNMT1* KD vs non-targeting control, and selecting the minimum P value and minimum FDR. Significant regions were determined by minimum FDR < 0.05, and hypo- and hyper-methylated regions were determined by mean difference of less than 0 or greater than or equal to 0, respectively. Signature overlap was calculated by counting the number of genes for each signature that appear in the significant hypo or hyper-methylated regions. Overlap with the genome-wide interferon-responsive screen from McQuade et al. 2025^73^ was conducted by overlapping significant positive or negative DEGs with the significant hypo-methylated or hyper-methylated regions, with some genes labelled on the volcano plot.

### DNMT1-low versus DNMT1-high differential gene-score analysis in human microglia

To assess whether DNMT1 activity is associated with altered microglial cell states in human tissue, we analyzed microglia from a single-nucleus ATAC-seq dataset comprising Alzheimer’s disease, progressive supranuclear palsy, and Pick’s disease across three brain regions with distinct vulnerability (precentral gyrus, insula, and calcarine cortex)^127^. Data were processed using ArchR (v1.0.3)^128^. High-quality nuclei were retained based on transcription start site (TSS) enrichment > 2 and ≥1,000 fragments per nucleus. Putative doublets were removed, and dimensionality reduction, batch correction, and clustering were performed to identify major brain cell types. Cell identities were annotated using canonical marker genes and validated by integrating with matched snRNA-seq data. A small number of low-confidence clusters and outlier samples were excluded. Microglial cells were then extracted for downstream analyses.

Differentially accessible regions (DARs) between DNMT1-low and DNMT1-high microglia were identified from the ArchR PeakMatrix using a Wilcoxon rank-sum test, and regions with FDR < 0.1 and |log2FC| > 0.5 were retained. CpG island annotations were obtained from the UCSC Genome Browser (GRCh38/hg38) using the CpG Islands track^129^. Enrichment of DNMT1-low DARs within CpG islands was assessed using Fisher’s exact test, with all tested ATAC peaks as the background. DNMT1-low DARs were strongly enriched in CpG islands, with 10,121 of 13,367 DARs overlapping CpG islands (odds ratio = 88.73, FDR < 2.2 × 10⁻¹⁶).

DNMT1 activity was quantified using the ArchR GeneScoreMatrix. For each microglial nucleus, the DNMT1 gene score was extracted, and cells with nonzero scores were retained. The 70th percentile of the nonzero DNMT1 gene-score distribution (1.003) was rounded to 1 and used as the cutoff. Cells with DNMT1 gene scores greater than 1 were classified as DNMT1-high, whereas those with scores less than or equal to 1 were classified as DNMT1-low. This yielded 15,592 microglia, including 4,688 DNMT1-high and 10,904 DNMT1-low cells.

Differential gene-score analysis was conducted using a pseudobulk strategy within the limma-voom framework^130^, with normalization performed using edgeR^131^. Gene scores were aggregated by sample, brain region, diagnosis, and DNMT1 group to generate pseudobulk profiles. Linear models were fit with the design: ∼ 0 + dnmt1_group + sample_batch + region + diagnosis. Differential gene scores between DNMT1-low and DNMT1-high microglia were assessed using empirical Bayes moderation. Multiple testing correction was applied using the Benjamini-Hochberg method, and genes with a false discovery rate (FDR) < 0.1 were considered significant.

### CRISPRi screen analysis

Analysis of CRISPR screens was as previously described^132^. Raw sequencing results were mapped with ‘sgcount’^132^. After, ‘crispr_screen’^132^ was used to perform differential enrichment analysis using a ‘drop first’ weighting strategy and Benjamini-Hochberg multiple hypothesis correction. Non-targeting control guides were grouped into “amalgam” genes in random groupings of five guides to mirror the five guides per targeted gene knockdown in our CRISPRi library. A z score cutoff of 5 was implemented to remove spurious non-targeting control cells.

### Single-cell RNA Sequence Alignment

The raw single-cell sequencing reads of gene expression libraries were processed using kb-python (version: v0.29.1) from kallisto-bustools^133^. First, using the GRCh38 human reference transcriptome with Ensembl annotations (release 113), a kallisto index was generated with the default k-mer size (k=31). The 10x reads were then pseudo-aligned to this index using ‘kb count’ from kallisto-bustools with the ‘standard’ workflow and the cell barcode whitelist for Single Cell 3’ v3.1 chemistry.

### Single-cell ADT Protein Assignment

The raw single-cell sequencing reads of antibody-derived tag (ADT) libraries were processed using the Cell Ranger software from 10x Genomics (version 9.0.0). The computational analysis was conducted on a high-performance computing (HPC) cluster configured with 48 GB of virtual memory and eight local cores. A custom feature reference file^83^ was used for detecting unique molecular identifiers (UMIs) associated with antibody-derived tags.

### CRISPRi knockdown demultiplexing (sgRNA assignment)

The raw sequencing reads of the PCR amplified sgRNA library were processed using kallisto-bustools^133^. First, using ‘kb ref’, a kallisto index of the sgRNA library was generated with k-mer size of 9. The reads were then pseudo-aligned to this index using ‘kb count’ from kallisto-bustools with the ‘standard’ workflow and the cell barcode whitelist for Single Cell 3’ v3.1 chemistry. sgRNA assignment was then conducted using geomux (version: v0.2.5)^132^, which conducts a hypergeometric test to calculate the significance of observing each sgRNA in each barcode. Cells were assigned to the majority sgRNA if they had a log-odds ratio above 1, Benjamini-Hochberg corrected P-value below 0.05, minimum UMI count to consider a barcode was 3, and minimum number of barcodes to consider a sgRNA was the default, 100. Once cells were assigned to a sgRNA, the barcodes were used to intersect the gene expression and ADT count matrices to store both the RNA and protein expression counts for each sgRNA-assigned cell in MuData format from muon (version: v0.1.7)^134^. Cells with no barcodes or multiple barcodes were not included for further analysis. Both the iTF-MG and iMG samples were processed this way. Of the two biological replicates from iTF-MG samples, only one includes the CITE-seq library and was included for this additional analysis.

### Quality Control and Integration

All QC was conducted using scanpy (version: v1.11.3)^135^. For both, the iTF-MG and iMG dataset with RNA and ADT counts, cells were filtered to include those with a minimum count of 200, and a mitochondrial count of less than or equal to 8%. Genes were filtered to include those with minimum cells of 10. For the iTF-MG experiment with only RNA counts, cells were filtered to include those with a minimum count of 1000. All other QC was conducted in the same way as the other datasets. For protein expression levels, four different normalization methods were initially tested and log_1_p-normalize was chosen as it had the highest median correlation to RNA expression as described in Haage et al. 2025^83^. Thus, for both RNA and protein expression levels, cells in each dataset were log_1_p-normalized to a target sum of 10,000. The top 5,000 highly variable genes were selected to conduct principal component analysis (PCA). Nearest neighbors were then calculated using the default 15 nearest neighbors, to then build UMAPs. Clustering was performed using different resolutions of the Leiden (version: v0.10.2)^136^ algorithm, and after setting the resolution to 1. Clusters with low *AIF1* expression (non-microglia cells) and low total counts were removed. 6 out of 18 clusters were removed from the iTF-MG replicate with only RNA data, whereas only 2 out of 13 clusters were removed from the iTF-MG replicate with both RNA and ADT data. Similarly, only 2 out of 9 clusters were removed from the iMG dataset. The two iTF-MG replicates were then concatenated and batch corrected using Harmony (version: v0.0.10)^137^, to get 3 final datasets used in analysis the iTF-MG and iMG MuData with both RNA and ADT counts, and the merged iTF-MG AnnData with the batch-corrected RNA counts. Two sgRNAs, FOXK1 sgRNA 2 (FOXK1_g2) and non-targeting control sgRNA 5 (non-targeting_g5) were also removed from analysis due to functional validation of FOXK1_g2 not producing knockdown (Figure S3H), and rigorous testing of non-targeting_g5 showing a perturbation effect through the Mixscale^87^ analysis described below. After QC, the resulting iTF-MG had 23,826 cells with RNA expression for 16,398 genes, and 9,040 cells with ADT expression for 180 proteins (166 proteins + 14 antibody controls). The iMG had 8,247 cells with RNA expression for 13,939 genes, and 8,247 cells with ADT expression for 180 proteins with 14 of those proteins included in the panel as controls.

### Integration of iTF-MG and iMG

Non-targeting cells were extracted from both iTF-MG and iMG and combined into one AnnData^135^ object for a total of 1,948 cells. Highly variable genes were re-calculated on joint dataset and principal component analysis (PCA) was conducted. Harmony^137^ was used to integrate the two models using parameters max_iter_harmony = 30, theta = 4, and sigma = 0.3. Nearest neighbors were then calculated using the default 15 nearest neighbors to then build UMAPs as depicted in Figure S3D.

### Mixscale Analysis

#### Perturbation Score Calculation-

To quantify strength of knockdown, a custom-modified version of Mixscale (modified from Jiang et al.^87^ https://github.com/reetm09/mixscale) was used to get a Z-score normalized perturbation score for each cell. The modification replaces Mixscale’s Bonferroni correction method for multiple hypothesis correction with the Benjamini-Hochberg method in the initial calculation of differentially expressed genes to determine if a targeted knockdown had a perturbation effect. In Mixscale, this DEG effect vector is then used to calculate a per-cell perturbation score, taking advantage of the full transcriptome for assessing effect of knockdown, and providing a continuous score more representative of CRISPRi knockdown in comparison to binary classification of perturbed vs not-perturbed which is more representative of CRISPR knockout. This was similarly done with the protein modality to get Mixscale perturbation scores at the protein level.

#### Weighted Differential Expression Analysis for genes and proteins-

The per-cell perturbation score calculated above is used as a weight such that cells with higher perturbation scores (stronger knockdown effect) are given a higher weight into a negative binomial regression model for differential expression analysis. For perturbations that don’t have an initial differential expression effect as classified by Mixscale, they underwent standard differential expression analysis. Similarly, differential abundant proteins (DAPs) were also calculated. All visualizations of DEGs and DAPs, including heatmaps (Figure 4C, 5C, 6C, 7C, S14C, S14N) and volcano plots (Figure 3D, 3E, 3F, 3J, 3K, 3L, 4D, 5D, 6D, 7D, S14D, S14O), were conducted in R, utilizing the dpylr^138^, ggplot2^139^, ComplexHeatmap^140^, and EnhancedVolcano^141^ packages. For heatmaps, individual cells (columns) were ordered by Mixscale score with a minimum score of 0, as depicted on histogram above heatmap. Each row depicts the log-normalized mRNA expression of the top 50 DEGs (top 25 downregulated and top 25 upregulated, FDR < 0.05). Non-targeting cells are unordered as they do not have Mixscale scores. For each perturbation, either non-targeting control or perturbed cells are downsampled to depict a balanced number of cells on the heatmap plot. Volcano plots are plotted with thresholds for significance at FDR < 0.05, with genes colored by signature. Full DEG and DAP tables can be found in Table S2, Tab 7, and Tab 8 for iMG and iTF-MG.

### Calculation of difference for Signature and Factor scores

Each signature was curated by literature (Table S2). Per-cell scores for each signature were calculated using a custom python implementation of the UCell^142^ algorithm, which is a robust method to calculate scores using Mann-Whitney U statistic test based on a ranked gene list. The signature score heatmap (Figure 2D) depicts median percentile shift of signature scores between perturbations and non-targeting control cells with a Mann-Whitney U test for significance and FDR < 0.05 for identifying perturbations that significantly affect signatures. For per-perturbation point plots (Figure 4A, 5A, 6A, 7A, S14A, S14L) error bars were calculated by 95% confidence intervals. Distributions of signature scores for each sgRNA were also plotted against non-targeting control cells normalized to AUC = 1 (Figure 4B, 5B, 6B, 7B, S14B, S14M). All median percentile shifts at gene and sgRNA level for iTF-MG and iMG can be found in Table S2, Tab 4, and Tab 5.

### Protein-Signature Correlations

For each protein-signature pair, the Pearson correlation was computed across all cells by comparing per-cell protein expression (from protein modality) and per-cell U-cell signature scores (from RNA modality). P-values were corrected for multiple testing per signature using the Benjamini-Hochberg FDR procedure. Resulting correlations for interferon, chemokine, disease-associated, and antigen-presenting signature scores plotted in Figure 3B and 3H, results for other signatures can be found in Table S2 Tab 6 and Tab 7.

### Stratification of Signature Score Enrichment by Perturbation Strength

Mixscale scores were binned by extracting bottom third, middle third, and top third of scores per perturbation, as they have different ranges. Median percentile shifts from NTC were calculated for each signature within each binned subset, error bars depict 95% confidence intervals and plotted in Figure S7A-D.

### Overlap of Differentially Expressed Genes (DEG) with External Datasets

A Fischer’s exact test was used to test for significance when conducting an overlap of DEGs for each perturbation compared to external datasets pulled from literature. Significant DEGs were chosen with a FDR < 0.05.

### scHPF Factor projection onto human microglia model

Both single-cell iTF-MG and iMG RNA were separately projected into a pre-existing consensus scHPF^143^ model from Marshe et al. 2025. ^80^ This model captures 26 non-orthogonal, latent factors representing microglial gene expression signatures defined in a dataset of 378,543 live human microglia from 127 donors and 17 unique brain tissue types, sequenced using 10x v3 chemistry. Three factors were excluded as they were related to sequencing batch effects. This is described in Marshe et al. 2025. Broadly, the model represents key biological processes in microglia such as metabolism, phagocytosis, antigen presentation, disease-associated states. For each query library (iTF-MG and iMG RNA), the raw UMI counts were extracted and downsampled to match the sequencing depth of the model using scHPF’s ‘downsample_loom.py’. All 8,478 genes in the scHPF model were present in the projected libraries. The downsampled counts were then projected into the model (scHPF’s ‘prep-like’, ‘project’, ‘score’) to get a per-cell factor score for each of the 26 factors in both microglia models. Radar plot of raw projected factor scores were plotted after subletting to non-targeting cells only (Figure S4A, S4D). Median percentile shift of factor scores were calculated with 95% confidence intervals per perturbation (Figure S4B, S4C, S4E, S4F). All median percentile shifts of factor scores at gene and sgRNA level for iTF-MG and iMG can be found in Table S2 Tab 11 and Tab 12.

### Predicting Knockdown Targets using SCENIC

The python implementation of the SCENIC^48^ algorithm (pySCENIC version 0.12.1)^144^ was used to assess the predicted gene-regulatory networks in our iPSC cells. Following the standard pipeline, two separate GRNs were constructed for iTF-MG and iMG respectively. Following the standard pipeline, raw expression matrices were inputted to the ‘grn’ command which utilizes GRNBoost2^145^ to build a network of adjacencies using co-expression of genes. Next, the default human motif databases (hg38_10kbp_up_10kbp_down_full_tx_v10_clust.genes_vs_motifs.rankings.feather and hg38_500bp_up_100bp_down_full_tx_v10_clust.genes_vs_motifs.rankings.feather), and default human annotations (motifs-v10nr_clust-nr.hgnc-m0.001-o0.0.tbl) were used for the motif pruning step using the ‘ctx’ command. This was used to infer enriched transcription factor (TF) binding motifs in co-expression modules, enabling the creation of regulons with source TFs and their predicted targets. During this step, ZNF532 dropped out as source TF so for the overlap analysis, predicted targets from GRN step were used instead of from the motif pruning step. For each constructed GRN, the set of predicted targets of PRDM1 in iMG and STAT2, DNMT1, IRF9 in iTF-MG was overlapped with the set of significant DEGs for those knockdowns. See Table S2 Tab 14 for results.

### Gene-set enrichment analysis

Gene-set enrichment analysis was conducted for positively and negatively differentially expressed genes in iMG: ZNF532 KD, PRDM1 KD and in iTF-MG, STAT2 KD, and DNMT1 KD against GO: Biological Processes, GO: Molecular Function, and GO: Cellular Component libraries using idea (https://github.com/noamteyssier/idea)^146^, which uses Enrichr^147^. Top 100 DEGs were selected when greater than 100 differentially expressed genes per perturbation. See Table S2 Tab 9 for results.

